# MosaicSim: A Novel Mosaic Variant Simulator Reveals Diminishing Returns of Ultra-High Coverage for Mosaic Variant Detection

**DOI:** 10.64898/2025.12.03.692191

**Authors:** Erik Stricker, Farhang Jaryani, Michal Izydorczyk, Chi-Lam Poon, Philippe Sanio, Adam Alexander, Sontosh K. Deb, Fritz Sedlazeck, Jeffrey Rogers, Elizabeth G. Atkinson

## Abstract

Genetic mutations within select cells of a tissue, termed mosaic variants (MV), are being increasingly recognized for their role in human disease. This growing interest underscores the need for specialized tools to detect and analyze MVs. However, such detection methods still lack thorough evaluation, largely due to missing benchmarking datasets that are large, reliable, and reflective of the complexity of biological samples. To address this gap, we developed MosaicSim, a tool for simulating variants in realistic sequencing data. The TweakVar workflow is at the tool’s core and represents a unique simulation pipeline that layers simulated MVs onto empirical whole genome sequencing data, generating a large, realistic ground truth dataset that combines the strengths of both simulation and biological data. To demonstrate the functionality of the workflow, we simulated 1,000 mosaic single nucleotide polymorphisms using TweakVar within whole genome sequencing files of different coverages. MVs were called with Illumina’s DRAGEN and compared to the ground truth. Our results show 150×–445× coverage performed comparably, with a true-positive rate between 50.4% (300×) and 54.9% (150×) and no false-positives detected. Across all samples, increasing variant allele frequency had a significant positive effect on call success. Additionally, we observed that call rates for variants in lower complexity regions improved with increasing read depth. We did not find significant effects attributable to specific mutation patterns or mean read map quality. MosaicSim fills a critical unmet need by providing representative, customizable ground truth datasets for MV benchmarking, enabling systematic evaluation and optimization of variant calling methods.

## Introduction

An increasing number of studies show that somatic variants, also termed mosaic variants (MVs), have significant clinical effects (Carli et al. 2022; Kingdom and Wright 2022; Truty et al. 2023). Recognizing the importance of these variants, the Somatic Mosaicism across Human Tissues (SMaHT) Network was established to systematically investigate the prevalence, distribution, and functional impact of somatic mutations across diverse human tissues (Coorens et al. 2025). As part of this initiative, our work aims to address a critical gap in the field: the lack of scalable, realistic benchmarking tools for MV detection. While recent improvements in high-coverage, high-accuracy whole genome sequencing (WGS) technologies have been instrumental in identifying disease associations, the success of these analyses fundamentally depends on the performance of mosaic variant callers, the computational cornerstone of any such somatic variation study. To ensure the optimal functionality of these callers, realistic validation datasets are essential (Ewing et al. 2015). Yet, scaling up MV detection methods remains challenging due to the scarcity of realistic, high-quality ground-truth datasets for benchmarking. In addition, the limited availability of comprehensive, cost-effective, and reliable validation resources hampers methodological innovation and drives up research costs by necessitating less efficient study designs.

To close this gap, we conceptualized MosaicSim, a tool allowing the simulation of mosaic variants in sequencing data, during the Fifth Baylor College of Medicine & DNAnexus hackathon in 2023 (Deb et al. 2024) and constructed it in its rudimentary form. Now, we present the production-ready TweakVar workflow central to the MosaicSim tool. The TweakVar workflow combines empirical data with ground truth MV simulations. In the past, mosaic variant callers have been validated using two different approaches with notable limitations. Firstly, a widely used technique in the field involves the physical mixing of samples. Almost all mosaic variant callers including MosaicHunter (Huang et al. 2017), MosaicForecast (Dou et al. 2020), DeepMosaic (Yang et al. 2023), Mutect2 (Cibulskis et al. 2013), Strelka2 (Kim et al. 2018), GATK’s HaplotypeCaller (McKenna et al. 2010), and Illumina’s DRAGEN (Dynamic Read Analysis for GENomics) (Scheffler et al. 2023; Allen et al. 2025) were at one point tested with ultra-deep sequenced physically mixed samples (Ha et al. 2023). While mixing tissue samples offers better biological relevance, it remains costly and is limited to variants present in the individuals. Therefore, this approach often overlooks rare disease-associated loci. In addition, high-complexity regions may still be inaccessible or lack sufficient confidence due to technical constraints.

In contrast, the most cost-effective and precise approach involves the use of simulators that create completely synthetic reads with variants based on a reference genome. Examples of simulation packages are PySim-sv (Xia et al. 2017), SCNVsim (Qin et al. 2015), VarSim (Mu et al. 2015), or SURVIVOR (Jeffares et al. 2017) with their use outlined in benchmarking papers such as DeepMosaic (Yang et al. 2023). While *in silico* simulations offer unmatched control and are considered the gold standard for benchmarking variant callers, even the most advanced error models for sequencing platforms cannot fully capture the biological complexity present in empirical samples, e.g., sample heterogenicity, lower coverage of repeat regions, and technical artifacts.

TweakVar addresses these limitations, combining the best of both approaches, by creating simulated MVs through modification of a fraction of existing reads within empirical WGS data BAM files, therefore modeling MVs in a realistic way. As a result, the workflow produces a simulated WGS file alongside a ground truth file. TweakVar provides access to the complete genomic landscape, including complex and low coverage regions, while being cost-effective capturing realistic technical errors, noise, and biological bias. Further, it allows users full control over simulated variant location, number, frequency, structure, and base, and can convert virtually any WGS file into a simulated dataset.

To validate the functionality and value of MosaicSim and its TweakVar workflow, we present our benchmarking results of Illumina’s DRAGEN variant caller applied to empirical WGS files with differing coverages and each of 1,000 simulated mosaic single nucleotide polymorphisms (SNPs). Our results demonstrate that variant detection success is strongly influenced by variant allele frequency and region-specific complexity. Most strikingly, a comparison of files with different coverages showed no significant differences between high and ultra-high coverage, and surprisingly, found that 150× read depth performed better than 450× and 300× due to technical features of the variant caller. Overall, our application of TweakVar demonstrates its potential to test variant callers at their performance limits in a cost-effective manner, uncovering error patterns that have so far remained hidden. While applied here on a small scale, this work highlights the clear possibility for TweakVar’s integration into large-scale MV research, which will enable more precise variant detection and, in turn, greater clinical relevance and impact.

## Results

### MosaicSim: A Framework for Simulating Mosaic Variants in Empirical Data

To address the need for realistic benchmarking datasets for mosaic variant callers, we developed MosaicSim, a software tool enabling the generation of simulated mosaic variant datasets incorporating empirical sequencing data. At the core of this tool is the TweakVar workflow, a three-stage process that generates a ground-truth variant list based on user-defined parameters, edits a corresponding fraction of reads within an existing BAM file according to the list, and merges the modified reads back into the original dataset (Fig. 1). This approach preserves the natural noise and complexity of the biological sample while providing a precise ground truth for performance evaluation. A full technical description of the workflow is provided in the Methods section.

**Figure 1.**
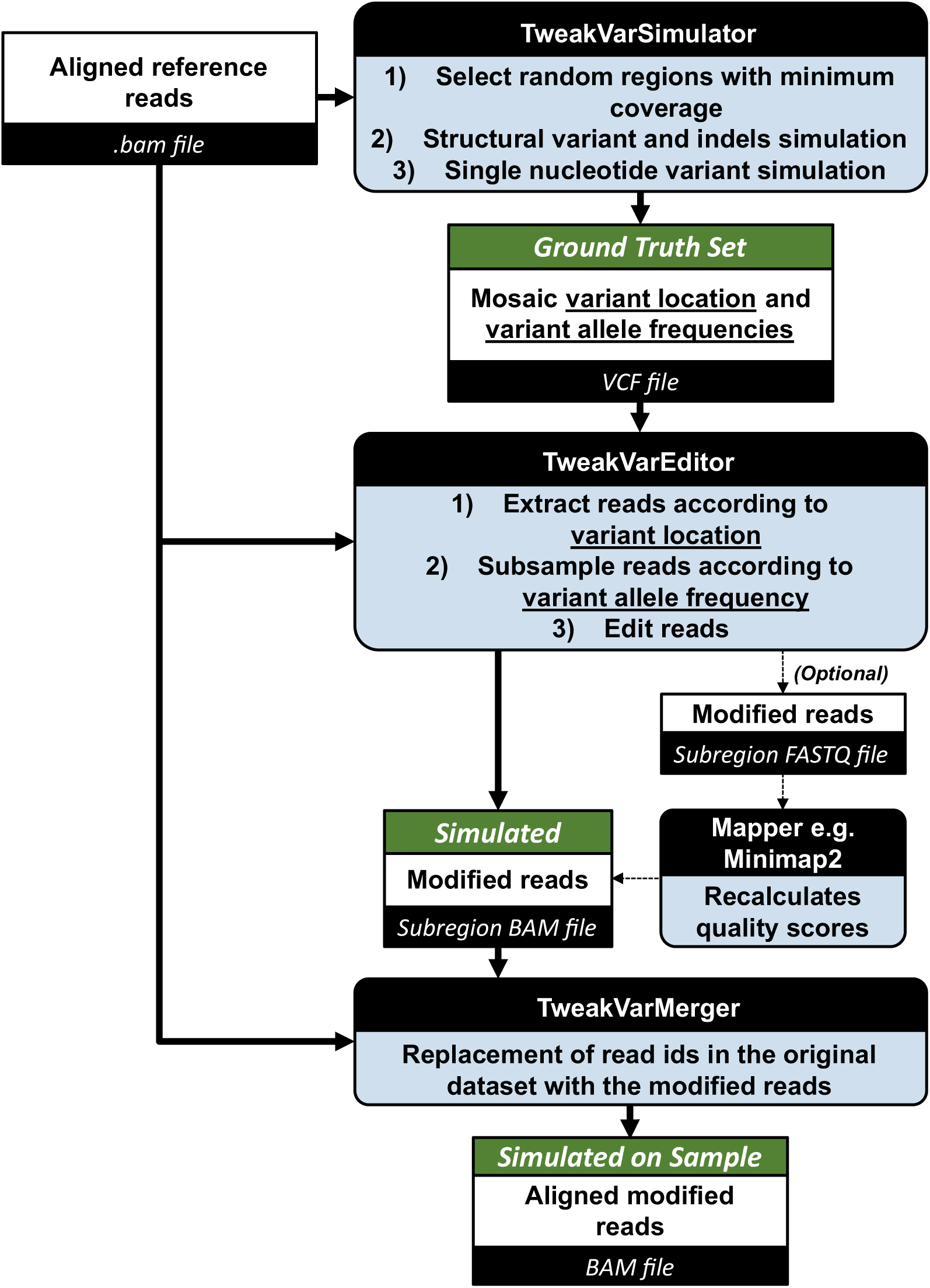
The TweakVarSimulator generates a random set of mosaic variants, including single nucleotide variants and structural variants. These variants can be customized by adjusting parameters such as variant allele frequency, number of variants to simulate, and variant type or size. The output of the TweakVarSimulator is a VCF file with variant location and allele frequencies. The TweakVarEditor uses the VCF file to extract reads from the input BAM file that overlap with the variant locations. Only a subset of the reads determined by the allele frequency are edited. The output BAM file retains the alignment info and now contains the edited reads keeping the query name of each read the same. Alternatively, a FASTQ instead of a BAM output file can be generated, which allows the remapping of the modified reads. The TweakVarMergers replaces the reads in the original dataset with the modified reads. This step is the computationally most intensive part, as it involves resorting reads.

### Creation of a Simulated Ground Truth Dataset for Benchmarking

We applied the TweakVar workflow of the MosaicSim tool to a high coverage (approximately 450× average coverage) WGS file and its down-sampled (300×, 150×, 100× and 30× average coverage) derivatives to verify the functionality of the pipeline and subsequently assess MV call success with Illumina’s DRAGEN MV caller. Using TweakVar, we successfully simulated 1,000 SNPs at the same locations in each of the five WGS files (see Supplemental Fig. S1 for an IGV browser screenshot of a modified region). We targeted simulation variant allele frequencies (VAFs) spanning 1%–5% for all coverage levels; however, local coverage fluctuations caused the final observed VAFs in the 450× file to range from 1% to 5.4% (Supplemental Table S1). This variability was, as expected, more pronounced in the lower-coverage files due to reduced read availability (Supplemental Table S2–S5). Overall, read depths at variants were normally distributed around the average coverage of the whole genome with an average coverage of 446 (SD = 128) for the 450× file, 301 (SD = 87) for the 300× file, 150 (SD = 45) for the 150× file, 101 (SD = 31) for the 100× file, and 30 (SD = 11) for the 30× file (Supplemental Figure S2). In addition, variants with VAF of 0.2%–0.6% were simulated within the highest-coverage file to mimic errors according to published technical error rates (Pfeiffer et al. 2018; Stoler and Nekrutenko 2021). We verified that all MVs were equally distributed across the genome (Supplemental Fig. S3). To ensure variants were placed into distinct reads, a minimum separation of 400 bp, twice the length of an Illumina read, was required. The observed distances were significantly larger, with the closest variants being 520 bp apart, and the average distance exceeding 2.9 Mb.

### Variant Detection Performance Plateaus Above 150× Coverage

After their creation, we performed variant calling on all five benchmarking BAM files with simulated MVs using Illumina’s DRAGEN pipeline. To quantify call success, we assessed true-positive rates by determining locus and allele match for each variant based on the TweakVar provided ground truth file. However, due to high inconsistencies in predicted and actual VAFs, we evaluated the quantification accuracy of DRAGEN through VAF estimation independently. Strikingly, 150× through 450× coverages performed comparably, with 150× actually having the highest overall true-positive rate of 54.9%, followed by 51.9% for 450×, and 50.4% for 300× (Tab. 1, Fig. 2A). Variant calling with 100× and 30× average coverage resulted in significantly lower call success. In contrast to variants on the autosomes, none of the 53 simulated SNPs on chromosomes X and Y were detected by DRAGEN even though the DRAGEN output included 171,548 variants on the sex chromosomes X and Y. Removal of MVs on chromosomes X and Y from the analysis increased the true-positive rate to 58% for 150×, followed by 54.8% for 450×, and 53.1% for 300× (Supplemental Table S6).

**Figure 2.**
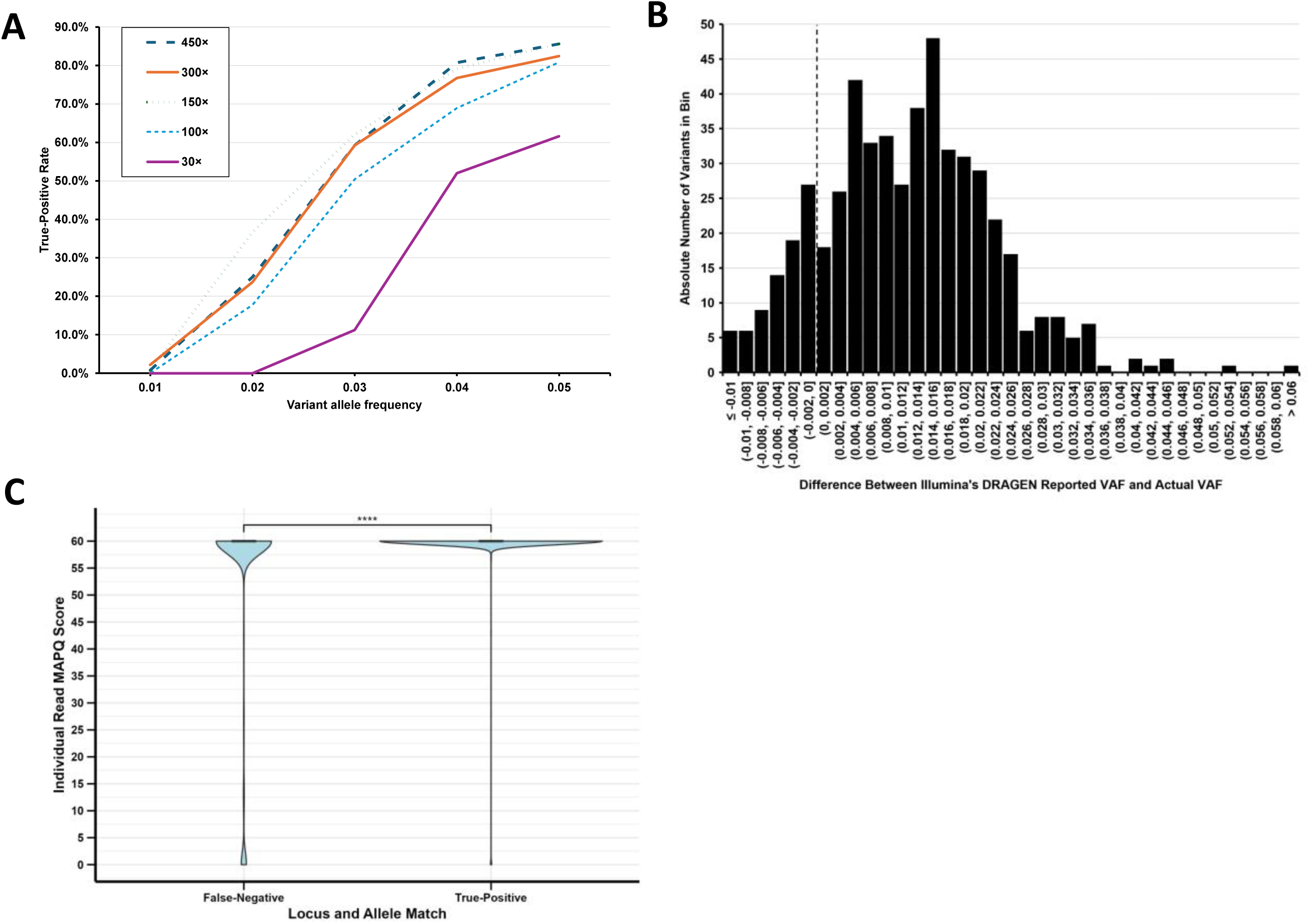
Evaluation of 1,000 simulated SNPs configured to 1%*–*5% VAF using the TweakVar workflow. MVs were variants called with Illumina’s DRAGEN variant caller and compared to the ground truth provided by the TweakVar workflow. (A) Comparison of true-positive files with approximately 450×, 300×, 150×, 100× and 30× average coverage. Note that actual VAF can vary based on read availability at the locus and therefore, indicated VAFs should be considered more of a range with minimum VAF as indicated. (B) Differences in the Illumina DRAGEN predicted and actual variant allele frequencies of the 520 simulated mosaic variants detected in the 450× file. The DRAGEN output produced an average difference of 1.35% (SD = 4.37%). The vertical dashed line indicates no VAF difference between DRAGEN and truth set. (C) Violin plot showing the distribution of high read map quality (MAPQ = 60) and low read map quality (MAPQ <5). Reads with unknown map quality score (MAPQ = 255) were removed. A statistical comparison was made using an unpaired, two-tailed Student’s t-test: **** p ≤ 0.0001.

**Table 1.** Overall true-positive, false-negative and false-positive rates were determined for MV calling applying DRAGEN to the 1,000 SNP benchmarking dataset.

| Coverage | True-Positive | False-Negative | False-Positive | P-Value |
| --- | --- | --- | --- | --- |
| 450× (original) | 51.9% | 48.1%<br>(<sup>‡</sup>0.1% wrong allele) | 0% | Referent |
| 1% VAF | 0.7% | 99.3% | 0% | Referent |
| 2% VAF | 25.0% | 75.0%<sup>‡</sup> | 0% | Referent |
| 3% VAF | 59.2% | 40.8% | 0% | Referent |
| 4% VAF | 80.7% | 19.3% | 0% | Referent |
| 5% VAF | 85.6% | 14.4% | 0% | Referent |
| 300× (down-sampled) | 50.4% | 49.6%<br>(0% wrong allele) | NA | 0.50 |
| 1% VAF | 2.2% | 97.8% | NA | 0.31 |
| 2% VAF | 23.7% | 76.3% | NA | 0.74 |
| 3% VAF | 59.2% | 40.8% | NA | 1 |
| 4% VAF | 76.8% | 23.2% | NA | 0.27 |
| 5% VAF | 82.4% | 17.6% | NA | 0.49 |
| 150× (down-sampled) | 54.9% | 45.1%<br>(0.1% wrong allele) | NA | 0.18 |
| 1% VAF | 0.7% | 99.3% | NA | 1 |
| 2% VAF | 36.6% | 63.4%<sup>‡</sup> | NA | 0.006** |
| 3% VAF | 62.0% | 38.0% | NA | 0.52 |
| 4% VAF | 79.1% | 20.9% | NA | 0.65 |
| 5% VAF | 85.6% | 14.4% | NA | 1 |
| 100× (down-sampled) | 44.3% | 55.7%<br>(0.1% wrong allele) | NA | 0.0006*** |
| 1% VAF | 0.0% | 100.0% | 0% | 0.32 |
| 2% VAF | 17.7% | 82.3%<sup>‡</sup> | NA | 0.054 |
| 3% VAF | 50.4% | 49.6% | NA | 0.048* |
| 4% VAF | 68.9% | 31.1% | NA | 0.002** |
| 5% VAF | 80.8% | 19.2% | NA | 0.31 |
| 30× (down-sampled) | 23.7% | 76.3%<br>(0.1% wrong allele) | NA | 2.84E-40**** |
| 1% VAF | 0.0% | 100.0% | NA | 0.32 |
| 2% VAF | 0.0% | 100.0%<sup>‡</sup> | NA | 3.32E-17**** |
| 3% VAF | 11.2% | 88.8% | NA | 2.37E-33**** |
| 4% VAF | 52.0% | 48.0% | NA | 2.49E-12**** |
| 5% VAF | 61.6% | 38.4% | NA | 1.27E-05**** |

True-positive and false-negative rates were determined based on locus and allele matches of simulated SNPs set to 1%–5% VAF with DRAGEN reported SNPs; false-positive rate was determined based on simulated SNPs set to 0.2%–0.6% VAF in 450× file. <sup>‡</sup>Includes one variant with correct locus but wrong allele. Asterisks indicate significant p-value for unpaired, two-tailed Student t-test conducted with 450× as referent: \* $p \leq 0.05$ , \*\* $p \leq 0.01$ , \*\*\* $p \leq 0.001$ , \*\*\*\* $p \leq 0.0001$ .

### Allele Frequency is a Stronger Determinant of Detection than Sequencing Depth

Next, we investigated the influence of VAF (Fig. 3.A–C), coverage at variant locus (Fig. 3.D–F), and read count with alternative allele (Fig. 3.G–I) on true-positive rates. Interestingly, we were able to detect a unified VAF cutoff of <2% for detection at all depths barring 30x, with only single outliers. The 30x file had a higher VAF cutoff of <4% (Fig. 3.A–C). In addition, we observed a clear positive effect of higher VAFs on call success across coverages (450× file p-value = 1.06E-88; 300× file p-value = 3.76E-77; 150× file p-value = 4.91E-69; 100× file p-value = 2.63E-79; 30× file p-value = 9.07E-58) across all variants. While coverage at the variant locus displayed significant effects for lower coverage files, such as the 30× file (p-value = 3.66E-09), 100× file (p-value = 0.002), and 150× file (p-value = 0.03), and no significant effect on call success for the 300× file (p-value = 0.226) and 450× file (p-value = 0.05), clear detection cutoffs could also be observed for locus coverage. Across all samples tested, we noted that variants in regions with coverages two standard deviations above and one standard deviation below were detected with a below 0.5% success rate (Fig. 3.D–F). As a combination of both observed effects, alternative allele counts had significant effects on true-positive rates across the board, including for the 450× file (p-value = 2.23E-52), 300× file (p-value = 4E-61), 150× file (p-value = 7.22E-43), 100× file (p-value = 6.8E-59), and 30× file (p-value = 1.68E-180). However, cutoffs differed between the coverages suggesting a stronger effect of allele frequency than absolute counts on detection. We also noticed a substantial discrepancy of true VAFs, which were calculated as the ratio of alternative to total read counts, and the VAF reported by DRAGEN with an average difference of 1.34% (SD = 4.38%) (Fig. 2B).

**Figure 3.**
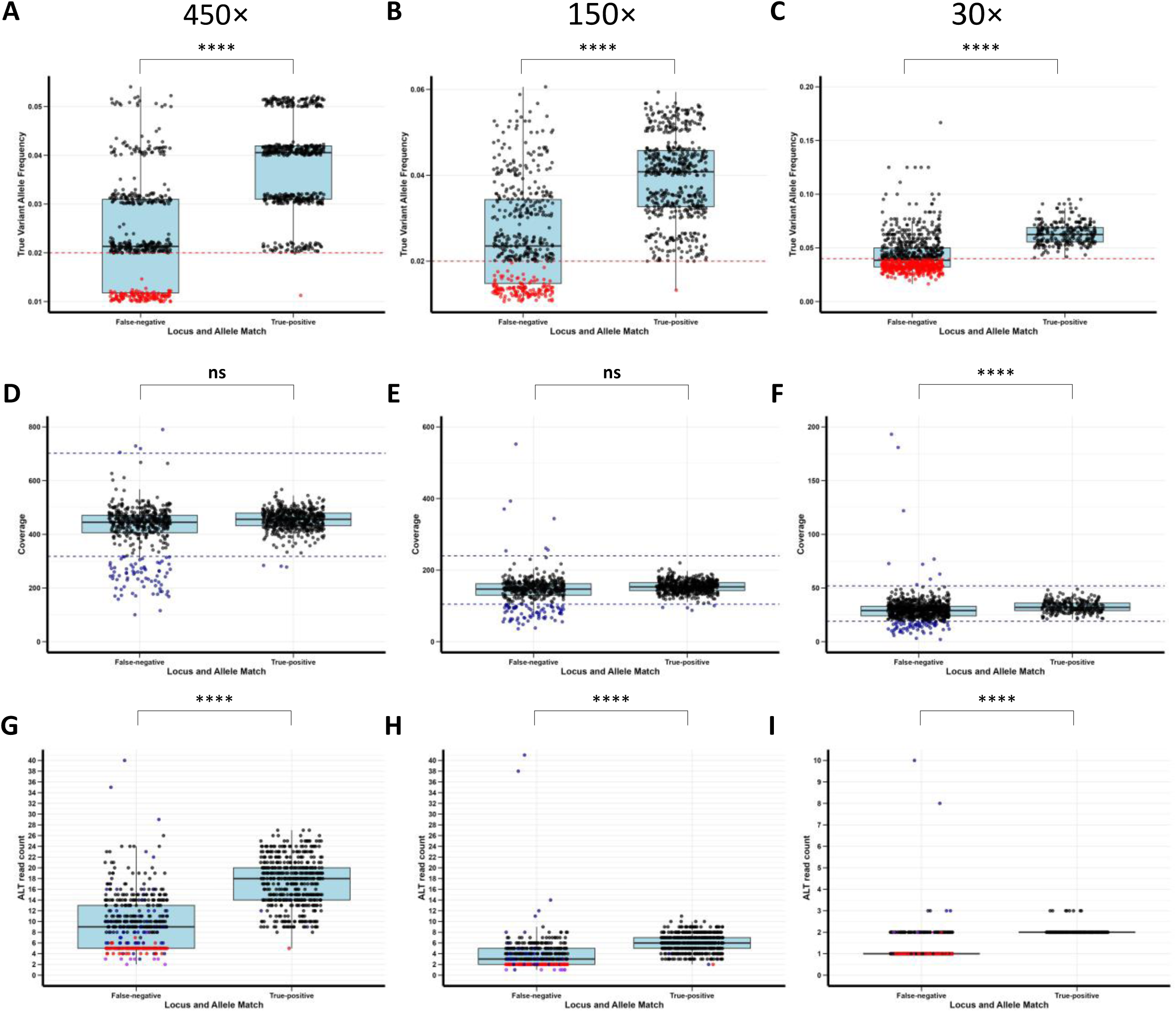
Comparison of true-positive and false-negative variant calls across different sequencing depths and metrics. Panels show the distribution of true-positive (locus and allele matched) versus false-negative (locus or allele unmatched) calls by: True variant allele frequency - (A) 450×, (B) 150×, and (C) 30× coverage files; coverage at variant location - (D) 450×, (E) 150×, and (F) 30× coverage files; and by alternative allele count for (G) 450×, (H) 150×, and (I) 30× coverage files. Variants with VAF below detection threshold were colored red (for 450× and 150× coverage panels A–B and G–H, <2% VAF; for 30× coverage panels C and I, <4%). Variants with very high (above two standard deviations) or low (below one standard deviation) coverage indicating a detection threshold were colored blue (for 450× files in panels D and G >702 and <318; for 150× files in panels E and H, >240 and <105; for the 30× files in panels F and I, >52 and <19). In panels G–I, variants that satisfied both coloring requirements were colored purple. Asterisks indicate significant p-value for unpaired, two-tailed Student t-test: ns = non-significant (p > 0.05), **** p ≤ 0.0001.

### Complexity Region-Specific Characteristics Influence Variant Calling

To assess locus-specific influences on call success, we initially compared true-positive rates of variants with known rsID and *de novo* variants without rsID (Supplemental Tables S1-S5). Statistical comparisons showed significantly higher true positive rates for annotated variants with a 7–8% increase for 100× (p = 0.045) and 450× files (p = 0.016). Since highly repetitive or structurally complex regions are difficult to map, they also tend to have a lower density of variants with assigned rsIDs. Therefore, we applied the genome region categorization masks by Ha *et al*. (publication in preparation) to our simulated variants (Fig. 4). The ‘easy’ category includes the most confidently mappable and variant-callable regions across population samples with minimal error in short-read calling. The ‘difficult’ mask contains moderately mappable regions across genomes, which are less confident for short-read variant calling. Lastly, the ‘extreme’ mask includes structurally complex and highly repetitive regions that are poorly resolved in pan-genome assemblies. Overall, we observed that easily mappable regions contained 75.9% of the bulk of the simulated variants and were increasingly mapped successfully with growing read depth (Fig. 4A). In addition, true-positive rates significantly increased from 23.3% in ‘difficult’ complexity regions for 30× coverage to 48.8% for 150× coverage (Fig. 4B). A similar, although smaller effect on true-positive rates could be observed for ‘extreme’ complexity regions with and increase from 10.9% for 30× coverage to 22.6% for 150× coverage (Fig. 4C). Lastly, we assessed other potential confounders and found that substitution type, *i.e.*, REF>ALT combinations (p = 0.783) had no significant effect on call success (Supplemental Figure S4).

**Figure 4.**
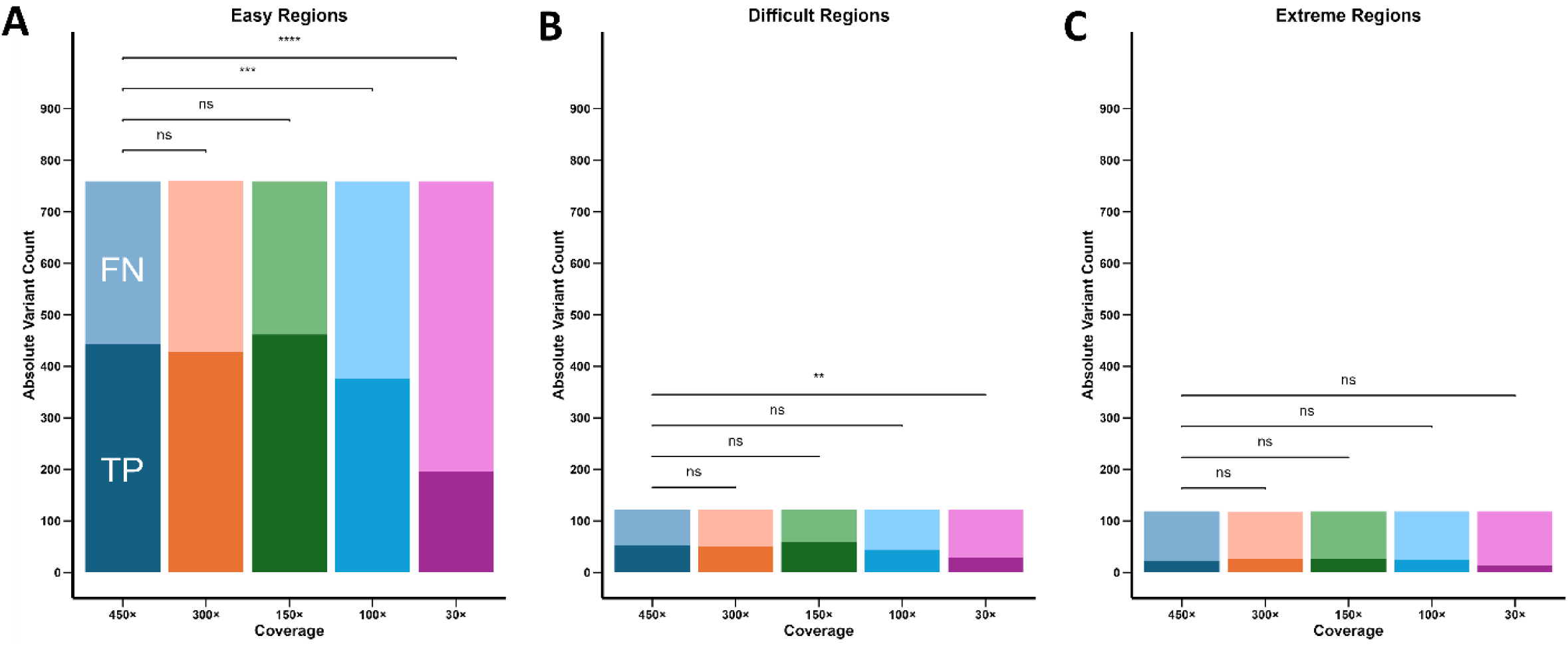
TweakVar simulated variants stratified by complexity region and different coverage files. (A) ‘Easy’ regions include most confidently mappable variants, while (B) ‘difficult’ and (C) ‘extreme’ contain increasingly complex and repetitive loci. Darker shaded areas represent true-positive (TP, locus and allele matched) variants, while lighter shaded areas annotate false-negative (FN, non-matched) variants. Comparisons were made using an unpaired, two-tailed Student’s t-test: ns = non-significant (p > 0.05), ** p ≤ 0.01, *** p ≤ 0.001, **** p ≤ 0.0001.

## Discussion

Here, we introduce MosaicSim, a new tool able to generate large, realistic ground truth datasets for MV detection by inserting simulated MVs into empirical WGS data with its TweakVar workflow. We demonstrated TweakVar’s capabilities by simulating 1,000 SNPs in WGS files of 30× to 450× coverage and used the resulting datasets to assess Illumina’s DRAGEN variant detection pipeline. In evaluating DRAGEN’s performance using TweakVar-simulated datasets, we observed several informative patterns that highlight both the strengths and limitations of current mosaic variant detection approaches. Our results showed that 150×, 300×, and 450× coverage files all performed similarly with 51.9%–54.9% true-positive rates. These findings highlight the need for variant calling pipeline assessments with extensive truth datasets as the MV detection rates we observed at 150× not only suggest a large potential for cost savings by opting out of ultra-high coverage approaches but also represented the best performance with existing variant calling tools. In addition, TweakVar’s implementation and dataset generation is cost-effective and highly flexible, making it well suited for optimizing study design of projects involving MV detection.

Secondary to overall coverage, we detected VAF as a critical factor in MV calling success. Our benchmarking datasets included variants targeted between 1% and 5% VAF distributed evenly across the genome. Using its default parameters for MV detection, Illumina’s DRAGEN v4.3 failed to detect MVs with a VAF below 2%–including variants we simulated with 0.2% to 0.6% VAF to emulate technical errors–with the exception of a few isolated cases. This reflects a known shortcoming of the machine learning model used by Illumina’s DRAGEN (Scheffler et al. 2023; Allen et al. 2025). Such limitations concerning rarer MVs might be overcome by using MV callers such as GATK (McKenna et al. 2010) which rely on absolute read cutoffs rather than on read and context evidence for low VAF calls. In addition, Illumina is in the alpha testing phase for high sensitivity mosaic detection in high-depth samples (>200×) which theoretically lowers the limit of detection to approximately 1.5% VAF (Illumina 2025).

Beyond the challenge of low-frequency VAF detection, we also observed that true-positive rates increased nearly linearly with rising VAF across all coverage levels, but plateaued at 85.6%. This is notably lower than the published single nucleotide variant (SNV) sensitivities of 92–99% established based on ultra-deep sequenced biological samples (Scheffler et al. 2023; Allen et al. 2025). These findings suggest that even at higher VAFs, detection performance may be constrained by factors not captured in traditional mixed empirical benchmarking datasets. A more comprehensive approach using simulated data as provided by MosaicSim in combination with existing benchmarking datasets is necessary to robustly validate caller performance and drive improvements in overall MV detection accuracy.

Upon stratification of the simulated MVs by genomic region complexity, we observed a direct correlation between false-negative rates and mapping difficulty, highlighting the challenges of variant detection in repetitive or structurally complex regions. Overall sequencing coverage had the largest influence on low complexity regions with easy mappability, more than doubling call success from 25.8% in the lowest coverage to 61.7% for 150× coverage. These findings matched our observations that known variants with rsIDs have significantly higher true-positive rates, and they are less abundant in low complexity and hard-to-map regions. This also emphasizes another shortcoming of relying exclusively on biological mixed samples for MV caller benchmarking, as these datasets typically simulate mosaicism by diluting high-frequency germline variants (Ha et al. 2022). As a result, they may not fully capture the complexity or technical challenges associated with detecting various true low-frequency mosaic variants. Our analyses of MAPQ scores further emphasized the problems caused by low quality regions. In our data, reads with low MAPQ scores were significantly enriched in false-negative loci. Encouragingly, newer MV callers, such as Sniffles2, have started to address this issue by implementing purely simulated datasets into their benchmarking. However, the lack of technical and biological noise still limits their ability to fully replicate the challenges of empirical data (Smolka et al. 2024). With TweakVar, we were able to overcome these limitations by simulating variants in complex regions within a realistic biological background.

The results of this study should be considered in light of certain limitations. First, we only tested a single MV calling pipeline with standard parameters which might not have been optimized to our biological sample or types of MVs. Nonetheless, Illumina’s DRAGEN is considered one of the leading MV callers and is intended to be used for all other samples generated by the SMaHT network. Considering our presented results, we would recommend further optimization and adjustment of parameters and/or benchmarking of other callers such as GATK’s HaplotypeCaller (McKenna et al. 2010), MosaicHunter (Huang et al. 2017), MosaicForecast (Dou et al. 2020), DeepMosaic (Yang et al. 2023), Mutect2 (Cibulskis et al. 2013), or Strelka2 (Kim et al. 2018). Secondly, we simulated a limited number of MVs and a single MV type on top of one singular sequencing background. To broaden its applicability, we have already incorporated capabilities into TweakVar to simulate not only SNVs but additionally indels and structural variants (SVs) of variable length in the current version. Furthermore, MosaicSim is easily scalable and ubiquitously applicable to not only WGS files of various origins but also to different sequencing platforms, including long-read samples. We intend to further investigate issues such as missing sex chromosome detection and apparent detection cutoffs by generating a larger simulated dataset with more different samples. This dataset will be made publicly accessible to supplement existing reference standards.

We consider MosaicSim a valuable and flexible resource that addresses a critical unmet need for both the SMaHT consortium and the broader genomics community. By providing a robust, cost-effective method for generating large-scale, realistic ground-truth datasets, MosaicSim will directly empower users to optimize and systematically benchmark variant detection pipelines. This capability is crucial for developing comprehensive guidelines and informing decisions regarding the most efficient data production strategies (Ha et al. 2023). The tool’s ability to layer MVs onto varied empirical backgrounds will accelerate the refinement of MV detection tools, allowing developers to test callers at their performance limits and uncover systematic error patterns in a way not possible with physically mixed or purely synthetic samples. Its application will enable systematic comparisons of caller performance across different sequencing depths and variant frequencies (Chen et al. 2020), facilitating the establishment of clear clinical best practices (Koboldt 2020). Ultimately, by enabling more precise and sensitive MV detection, MosaicSim will enhance the translational value of MV research and its future clinical relevance.

## Methods

### TweakVar: The MosaicSim Simulation Workflow

TweakVar creates simulated MVs based on a user-specified or random list of mutations for which it modifies a fraction of existing reads within a user-provided WGS BAM file to result in a desired MV frequency. To accomplish this task, the pipeline is designed to follow a three-step process (see Fig. 1). First, the TweakVarSimulator takes an existing BAM file with aligned sequencing reads and selects random regions with minimum coverage within which it simulates variants. The workflow offers full control over the number of simulated SNPs, indels, or SVs, allele frequency ranges for both SNVs and SVs separately, minimum and maximum length of SVs, and ratio of insertions and deletions. The resulting VCF file provides the truth set with mosaic variant locations and allele frequencies and serves as the map file for the TweakVarEditor, which extracts reads according to the indicated locations, subsamples them giving the provided variant allele frequency, and then edits them. Output can be either an aligned BAM file with the modified reads or a FASTQ file, which can be remapped with a tool such as Minimap2 (Yuan et al. 2017; Danecek et al. 2021). The latter approach carries the risk of losing reads due to the introduced changes, potentially increasing rates of misalignment, but allows the recalculation of read quality scores. In the final step of the pipeline, the TweakVarMerger replaces the old reads in the input BAM file with the modified reads from the simulated BAM file to output a new BAM file with simulated mosaic variants. This simulated BAM file, in combination with the VCF truth file, can be used to benchmark any mosaic variant caller. Prior to the initial step, the user can further down-sample high coverage files to evaluate different read depths. The TweakVar simulation workflow was developed in Python 3.10.

### Generation of a Benchmarking Dataset with Simulated Mosaic Variants

To benchmark Illumina’s DRAGEN (Dynamic Read Analysis for GENomics) (Scheffler et al. 2023; Allen et al. 2025) variant calling pipeline, we obtained the HAPMAP6 benchmark sample mapped to GrCh38 and in CRAM format from the SMaHT Data Analysis Center (DAC) due to its high average read depth of 445× as a foundation for our simulated datasets. We used the samtools (version 1.21) view and index functions to convert the file into BAM format.

Using the TweakVar workflow, we simulated 1,000 SNPs in a WGS file with 445x average coverage and its down-sampled files with 300×, 150×, 100× and 30× coverage. We set VAFs from 1% to 5% to assess false-negative rates across the range of VAFs. For down-sampled files, the VCF SNV file was with the --SNV_truth_file, and --ignore_minimum_cov tags to ensure simulation of MV at identical locations and with the same alternate allele. In addition, we simulated 1,000 SNPs with VAF between 0.2% and 0.6% in the highest coverage file to emulate errors according to published technical error rates (Pfeiffer et al. 2018; Stoler and Nekrutenko 2021). These SNPs were placed 1,500 bp upstream of corresponding higher frequency variants to ensure similar distribution without sequencing read interference. Variants were randomly distributed across all autosomes and X and Y sex chromosomes. The TweakVarSimulator step was executed using a single task allocated 4 CPUs and 32 GB of memory and took 6 minutes and 17 seconds for the simulation of 1,000 SNPs. The TweakVarEditor step was run with identical resource allocation: one task, 4 CPUs, and 32 GB of memory and took 1 minute and 57 second to create the sub-BAM file with 1,000 SNPs. The final TweakVarMerger step was submitted as a batch job configured to use 4 tasks, with 4 CPUs per task, and a total of 64 GB of memory and took nearly 3 days with 72 hours, 43 minutes and 29 seconds to modify the 450× file with 1,000 SNPs. The complete code used can be found in Supplemental File S1. We used the IGV browser application (Consortium 2024) to visually validate the successful modification of sequencing reads for select locations.

### Variant Calling DRAGEN and Performance Assessment

We completed DRAGEN v4.3 variant calling with mosaic detection using the simulated BAM files at different coverage levels as inputs. All parameters used can be found in the Supplemental File S2. To assess the variant calling success, we compared each truth VCF file generated by TweakVar to each hard-filtered VCF file output provided by Dragen. We defined a locus match as having identical chromosome and start position in the truth file and Dragen output. We defined an allele match as having identical reference and alternative allele in the truth file and Dragen output. We evaluated locus and allele match rates based on actual allele frequency according to the truth file, alternative/simulated allele counts, coverage at variant site, base substitution type, mean read map quality score (MAPQ), and *de novo* vs. known variation.

### Genomic Stratification by Complexity Region

Genomic region stratification was performed using version 2 of complexity region masks by Ha *et al*. (publication in preparation). Version 2 adopts a biologically grounded approach using pan-genome assemblies and widely accepted mappability masks to divide the genome into mapping categories: ‘easy’, ‘difficult’, and ‘extreme’. The ‘easy’ region is based on the 1KG strict mask (GRCh38 genome accessibility mask) and includes confidently mappable and variant-callable regions across population samples with minimal error in short-read calling (Genomes Project et al. 2015). The ‘difficult’ regions represent the pm151b (PanMask strict) (Li 2025) but not in 1KG strict mask describing moderately mappable regions across the pangenome with less confidence for short-read variant calling. PanMask is a recent pan-genome–based approach that uses hundreds of high-quality assemblies to define sample-agnostic ‘easy’ regions, which offer a better balance between callability and genome coverage and are less biased by short-read aligner heuristics or reference genome limitations (Li 2025). The ‘extreme’ regions comprise all loci outside the pm151b set and 1KG strict mask, which consist of highly repetitive or structurally complex regions that are poorly resolved by pangenome or mappability filters. Overall, the ‘easy’ mask covered 74% of the genome and 78% of Non-N bases, while the ‘difficult’ mask comprised 10% of the genome (11% non-N bases) and the ‘extreme’ mask contained 16% of the genome (11% of non-N bases).

### Statistical Analysis

For statistical analyses, we compared false-negative variant calls (no locus or allele match in DRAGEN generated VCF file despite presence in the truth set) to true-positive calls (locus and allele match between the truth file and DRAGEN output). We conducted Student’s t-tests and reported the p-value for impact assessment of actual allele frequency according to truth file, alternative/simulated allele counts, coverage at variant site, mean read map quality scores, *de novo* vs. known variation, and comparisons within complexity regions. We performed a Pearson’s Chi-squared test to assess the call success between different base substitution types. Furthermore, we compared different overall coverages to the highest coverage sequencing file separated by simulated variant allele frequencies in Student’s t-tests as well.

## Data and Code Availability

This study was conducted as part of the NIH Common Fund consortium initiative, Somatic Mosaicism across Human Tissues (SMaHT). Benchmarking samples are available to qualified researchers per request to the SMaHT Network. The highest coverage file described in this study was the HAPMAP6 benchmark sample, which is the Coriell HAPMAP mix of 6 samples (83.50% HG005, 10% HG02622, 2% HG002, 2% HG02257, 2% HG02486, and 0.5% HG00438) (Abyzov 2025). The data used in this work was provided by the SMaHT DAC on behalf of the SMaHT Network. More information about the SMaHT Network and data is available online at https://smaht.org and at https://data.smaht.org.

MosaicSim is an open source software available at https://github.com/erikstricker/MosaicSim/ alongside installation instructions, and a detailed user tutorial with publicly available toy data. All analyses presented in this manuscript were performed using TweakVar version [v1.0.0] of the software available as Supplemental File S3.

## Competing Interests Statement

The authors have no competing interests to declare.

## Acknowledgements

We are extremely grateful to the SMaHT donors, and donor families, who have generously provided such precious gifts to support this important work. This research was supported by NIH (R01 HG012869 to E.G.A.), and the NIH Common Fund, through the Office of Strategic Coordination/Office of the NIH Director under awards U24 MH133204, U24 NS132103, UG3 NS132024, UG3 NS132061, UG3 NS132084, UG3 NS132105, UG3 NS132127, UG3 NS132128, UG3 NS132132, UG3 NS132134, UG3 NS132135, UG3 NS132136, UG3 NS132138, UG3 NS132139, UG3 NS132144, UG3 NS132146, UM1 DA058219, UM1 DA058220, UM1 DA058229, UM1 DA058230, UM1 DA058235, and UM1 DA058236.

## Supplemental Material

Supplemental Tables and Figures.docx

Supplemental Table S1.csv

Supplemental Table S2.csv

Supplemental Table S3.csv

Supplemental Table S4.csv

Supplemental Table S5.csv

Supplemental File S1.txt

Supplemental File S2.txt

Supplemental File S3.zip

## Notes

### Competing Interest Statement

The authors have declared no competing interest.

