## Supplemental Tables and Figures for "MosaicSim: A Novel Mosaic Variant Simulator Reveals Diminishing Returns of Ultra-High Coverage for Mosaic Variant Detection"

### Supplemental Tables & Figures

**Supplemental Table S6.** Overall true-positive, false-negative and false-positive rates determined for MV calling applying DRAGEN to the 947 out of 1,000 autosomal SNPs of the benchmarking dataset.

| Coverage | True-positive | False-negative | False-positive |
|---|---|---|---|
| **450× (original)** | 54.8% | 45.2%<br>(0.1% wrong allele) | 0% |
| **300× (down-sampled)** | 53.1% | 46.9%<br>(0% wrong allele) | NA |
| **150× (down-sampled)** | 58.0% | 42.0%<br>(0.1% wrong allele) | NA |
| **100× (down-sampled)** | 46.8% | 54.2%<br>(0.1% wrong allele) | NA |
| **30× (down-sampled)** | 25.0% | 75.0%<br>(0.1% wrong allele) | NA |

True-positive and false-negative rates were determined based on locus and allele match of simulated SNPs set to 1% - 5% VAF with DRAGEN reported SNPs; false-positive rate was determined based on simulated SNPs set to 0.2% - 0.6% VAF in 450× file

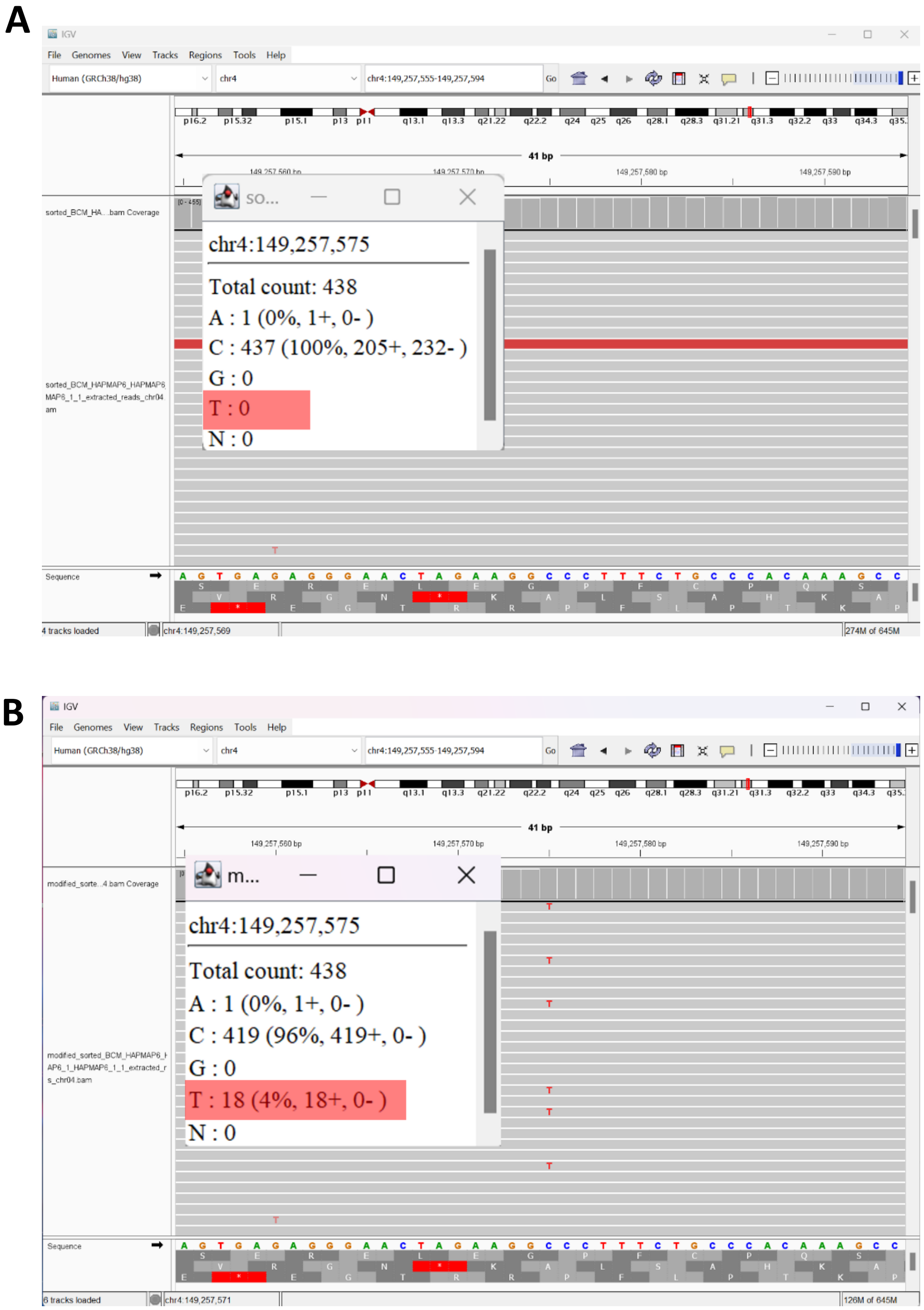


**Supplemental Figure S1.** IGV browser screenshot of the approximately 450× BAM file (A) before and (B) after introduction of a mosaic SNP by modifying approximately 4% of the reads at position chr4: 149,257,575 using the TweakVar workflow.

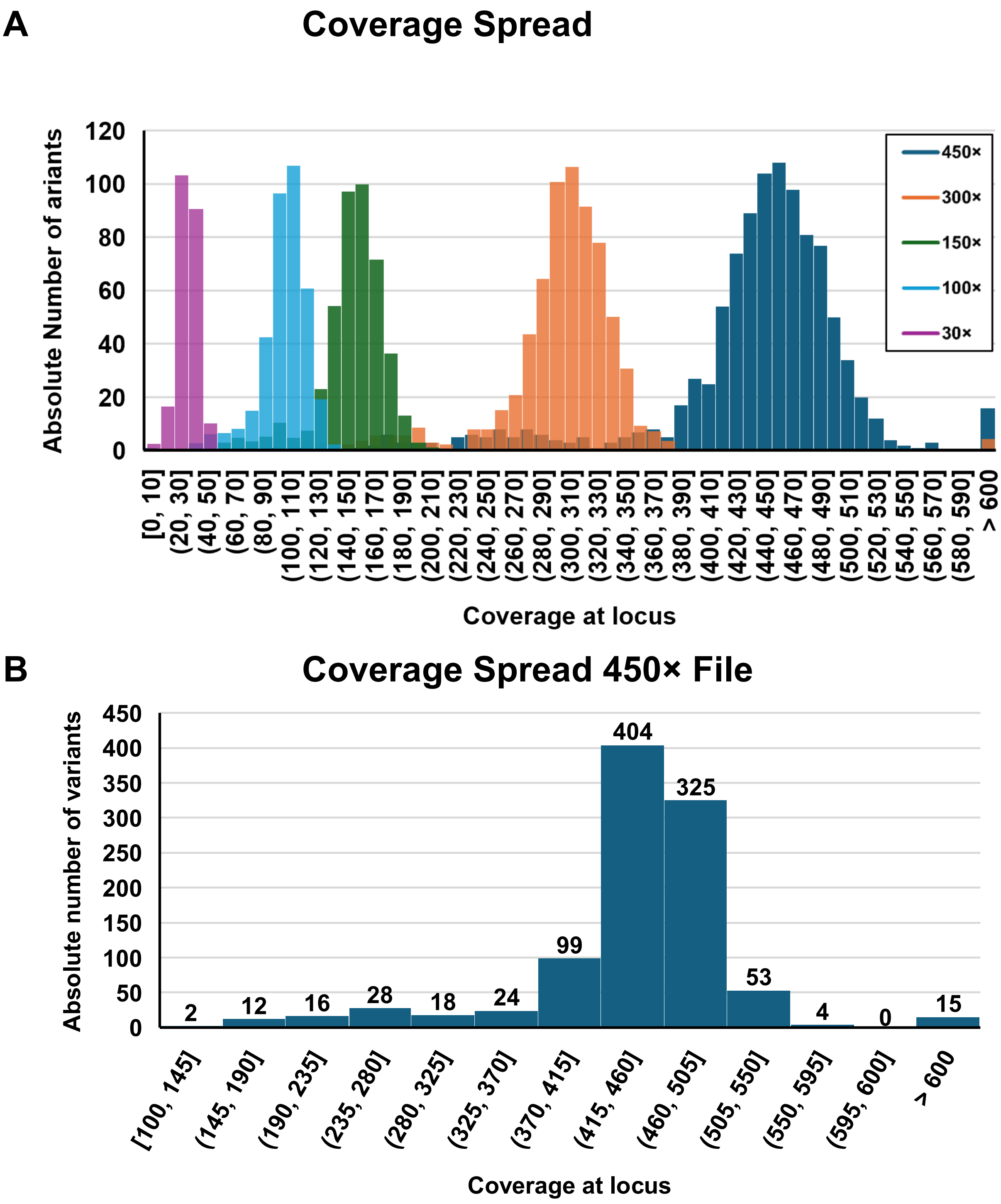
A
Coverage Spread
Absolute Number of ariants
120
100
80
60
40
20
0
450×
300×
150×
100×
30×
[0, 10]
(20, 30]
(40, 50]
(60, 70]
(80, 90]
(100, 110]
(120, 130]
(140, 150]
(160, 170]
(180, 190]
(200, 210]
(220, 230]
(240, 250]
(260, 270]
(280, 290]
(300, 310]
(320, 330]
(340, 350]
(360, 370]
(380, 390]
(400, 410]
(420, 430]
(440, 450]
(460, 470]
(480, 490]
(500, 510]
(520, 530]
(540, 550]
(560, 570]
(580, 590]
> 600
Coverage at locus
B
Coverage Spread 450× File
Absolute number of variants
450
400
350
300
250
200
150
100
50
0
2
12
16
28
18
24
99
404
325
53
4
0
15
[100, 145]
(145, 190]
(190, 235]
(235, 280]
(280, 325]
(325, 370]
(370, 415]
(415, 460]
(460, 505]
(505, 550]
(550, 595]
(595, 600]
> 600
Coverage at locus

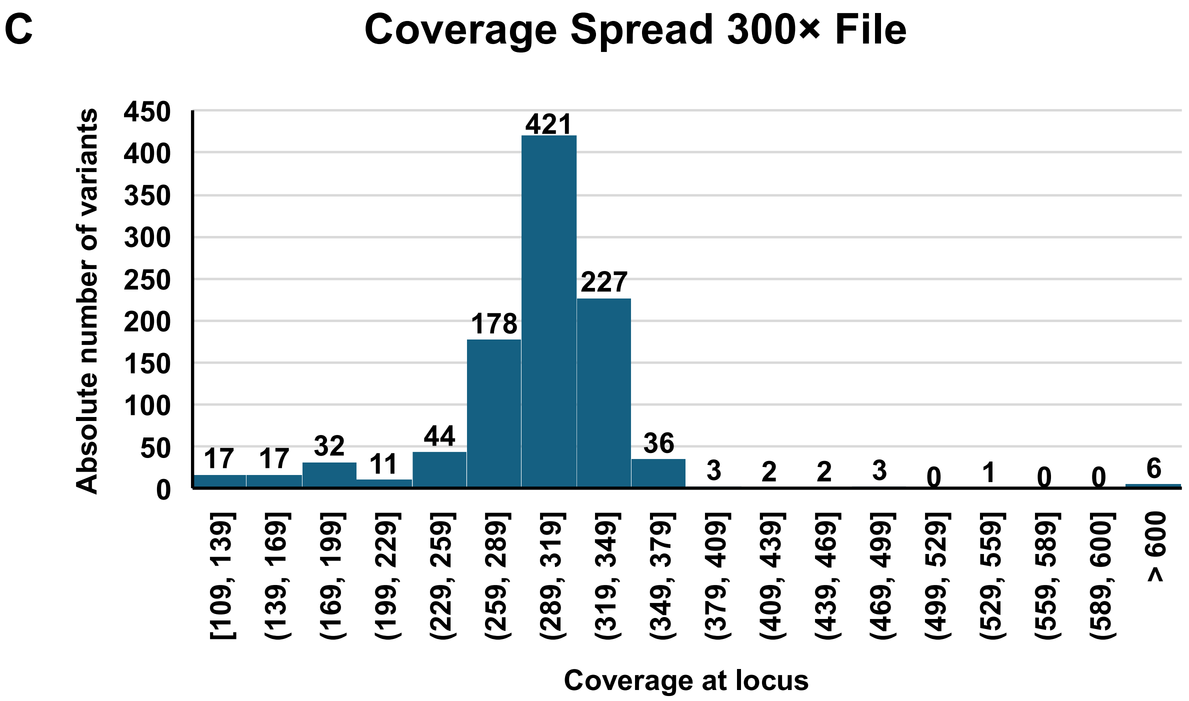
C
Coverage Spread 300× File
Absolute number of variants
450
400
350
300
250
200
150
100
50
0
17
17
32
11
44
178
421
227
36
3
2
2
3
0
1
0
0
6
[109, 139]
(139, 169]
(169, 199]
(199, 229]
(229, 259]
(259, 289]
(289, 319]
(319, 349]
(349, 379]
(379, 409]
(409, 439]
(439, 469]
(469, 499]
(499, 529]
(529, 559]
(559, 589]
(589, 600]
> 600
Coverage at locus

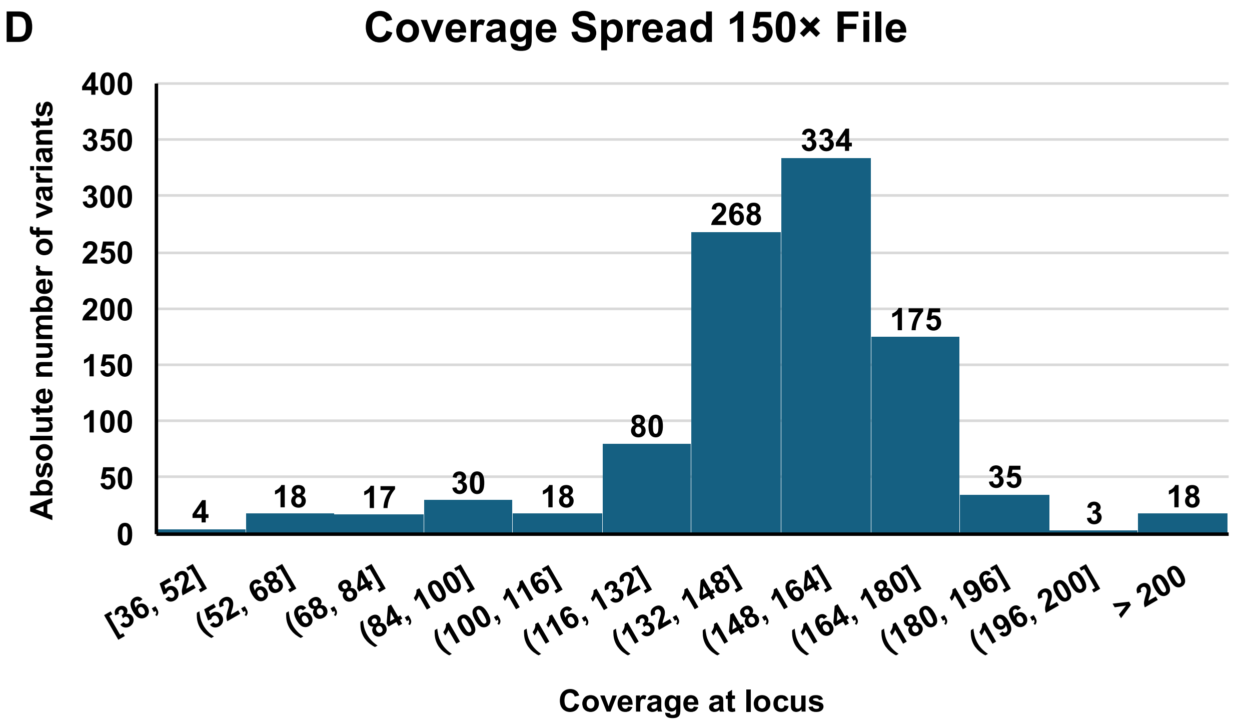
D
Coverage Spread 150× File
Absolute number of variants
400
350
300
250
200
150
100
50
0
4
18
17
30
18
80
268
334
175
35
3
18
[36, 52]
(52, 68]
(68, 84]
(84, 100]
(100, 116]
(116, 132]
(132, 148]
(148, 164]
(164, 180]
(180, 196]
(196, 200]
> 200
Coverage at locus

E

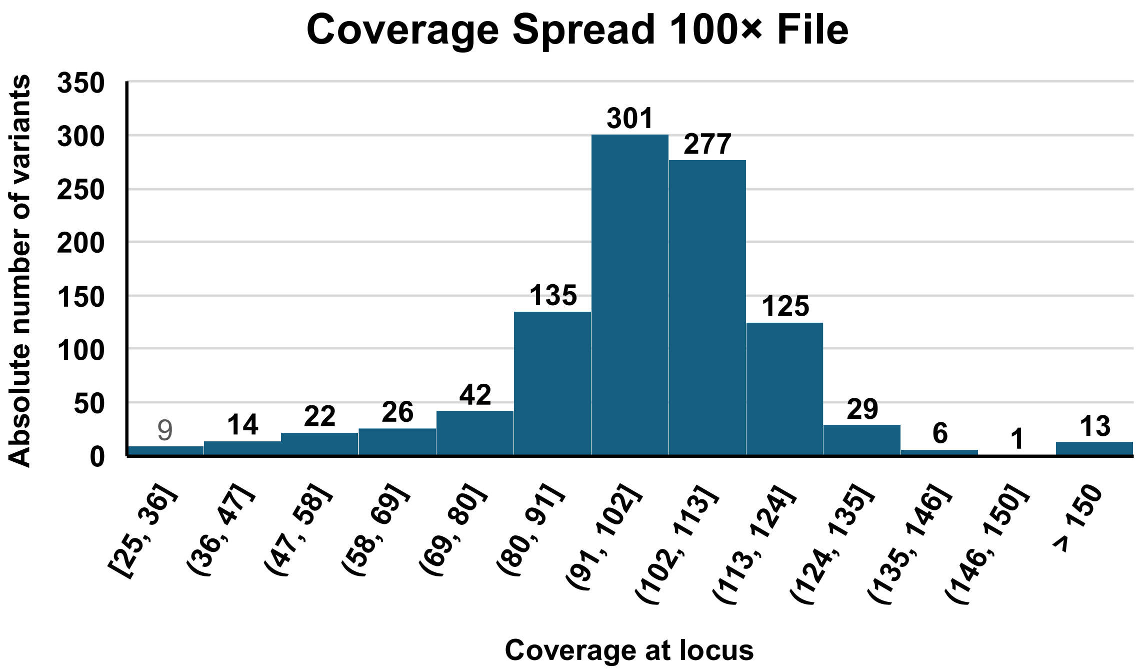


F

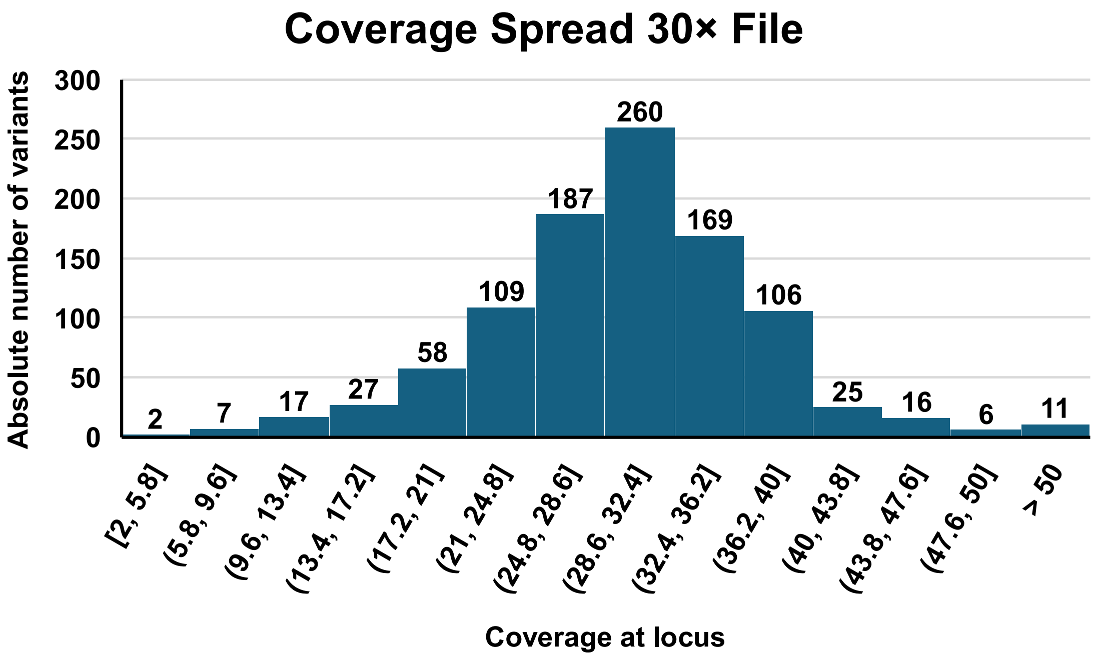


**Supplemental Figure S2.** Coverage/read depth at the loci of the 1,000 variants generated with the TweakVar workflow. Distribution of the (A) all variants, (B) 450× variants with an average coverage of 446 (Standard Deviation = 128), (C) 300× variants with an average coverage of 301 (Standard Deviation = 87), (D) 150× variants with an average coverage of 150 (Standard Deviation = 45), (E) 100× variants with an average coverage of 101 (Standard Deviation = 31), and (F) 30× variants with an average coverage of 30 (Standard Deviation = 11). Numbers of above the bars indicate the absolute number of variants within the bin.

A

B

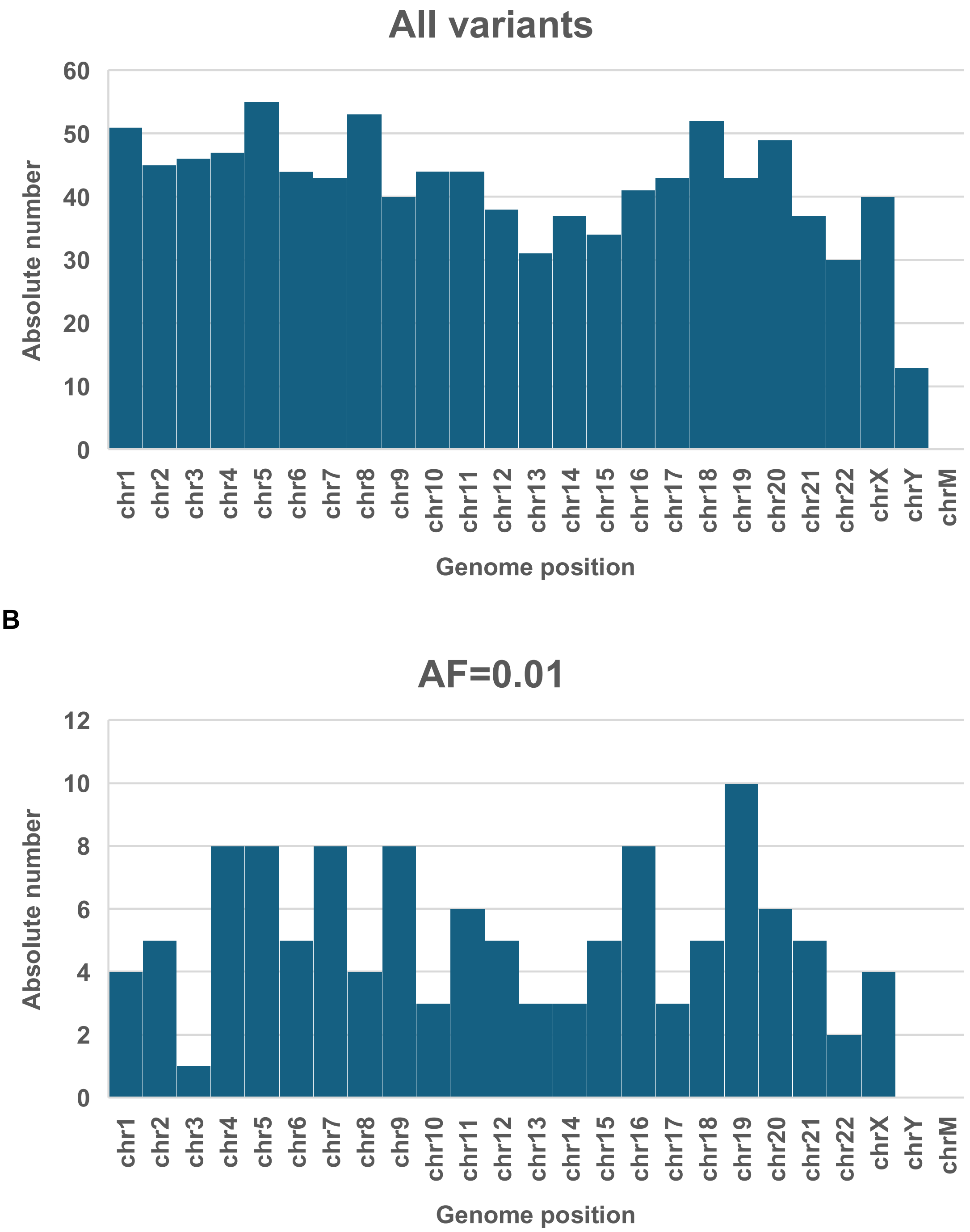
All variants
Absolute number
60
50
40
30
20
10
0
chr1
chr2
chr3
chr4
chr5
chr6
chr7
chr8
chr9
chr10
chr11
chr12
chr13
chr14
chr15
chr16
chr17
chr18
chr19
chr20
chr21
chr22
chrX
chrY
chrM
Genome position
AF=0.01
Absolute number
12
10
8
6
4
2
0
chr1
chr2
chr3
chr4
chr5
chr6
chr7
chr8
chr9
chr10
chr11
chr12
chr13
chr14
chr15
chr16
chr17
chr18
chr19
chr20
chr21
chr22
chrX
chrY
chrM
Genome position

C

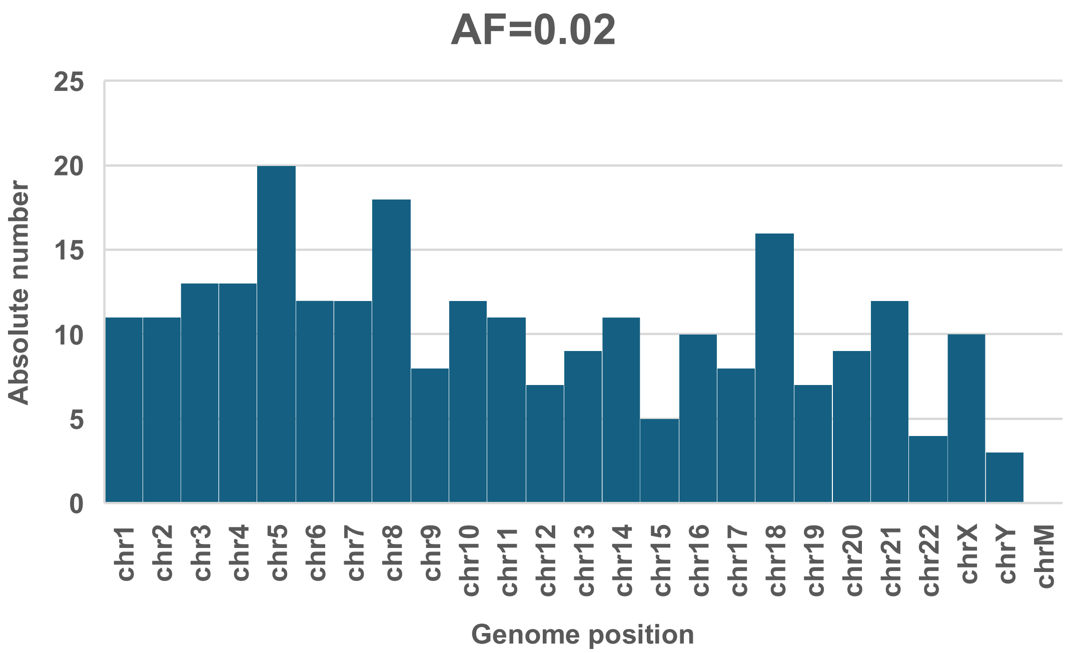
AF=0.02
Absolute number
25
20
15
10
5
0
chr1
chr2
chr3
chr4
chr5
chr6
chr7
chr8
chr9
chr10
chr11
chr12
chr13
chr14
chr15
chr16
chr17
chr18
chr19
chr20
chr21
chr22
chrX
chrY
chrM
Genome position

D

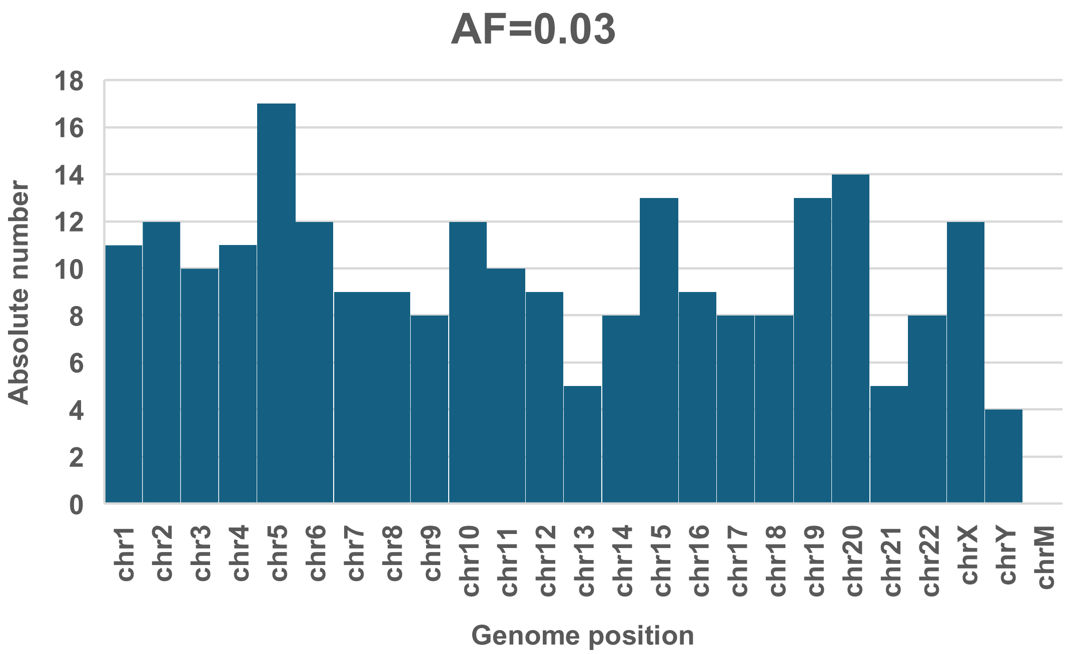
AF=0.03
Absolute number
18
16
14
12
10
8
6
4
2
0
chr1
chr2
chr3
chr4
chr5
chr6
chr7
chr8
chr9
chr10
chr11
chr12
chr13
chr14
chr15
chr16
chr17
chr18
chr19
chr20
chr21
chr22
chrX
chrY
chrM
Genome position

**E**

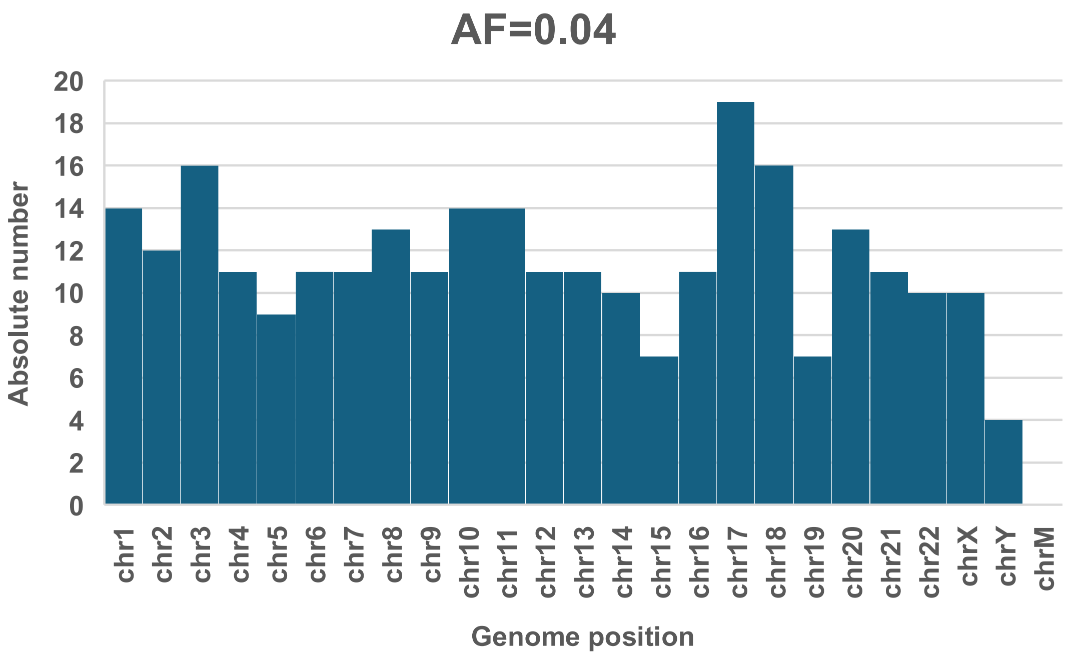


**F**

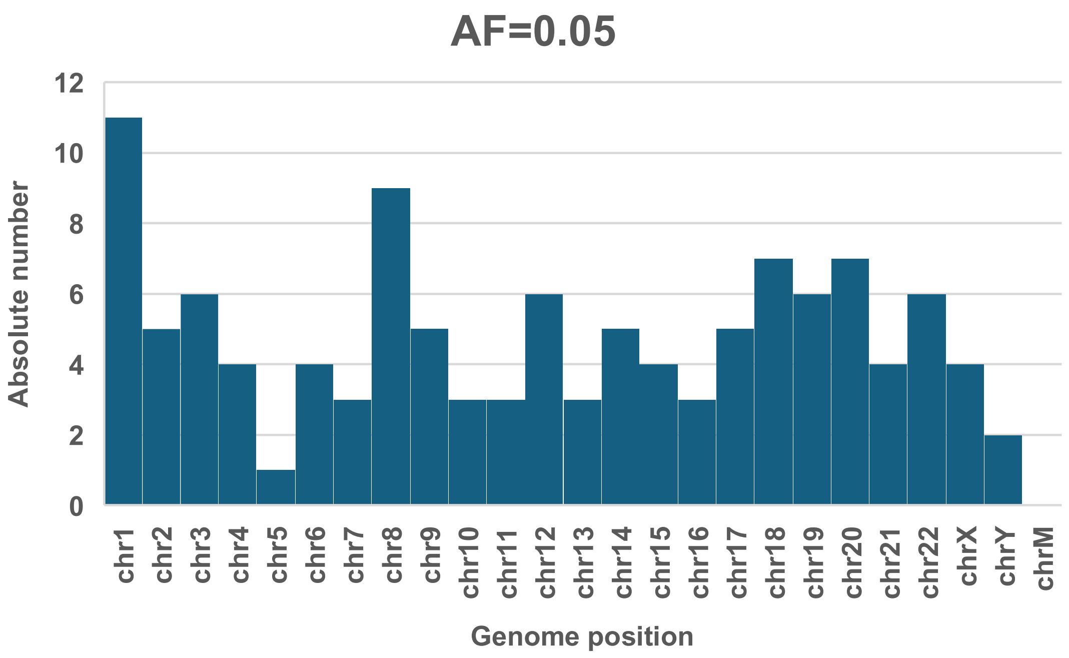


**Supplemental Figure S3.** Distribution of the 1,000 variants generated with the TweakVar workflow across the genome. (A) Overall variants, and variants set to (B) 1%, (C) 2%, (D) 3%, (E) 4%, and (F) 5% variant allele frequency.

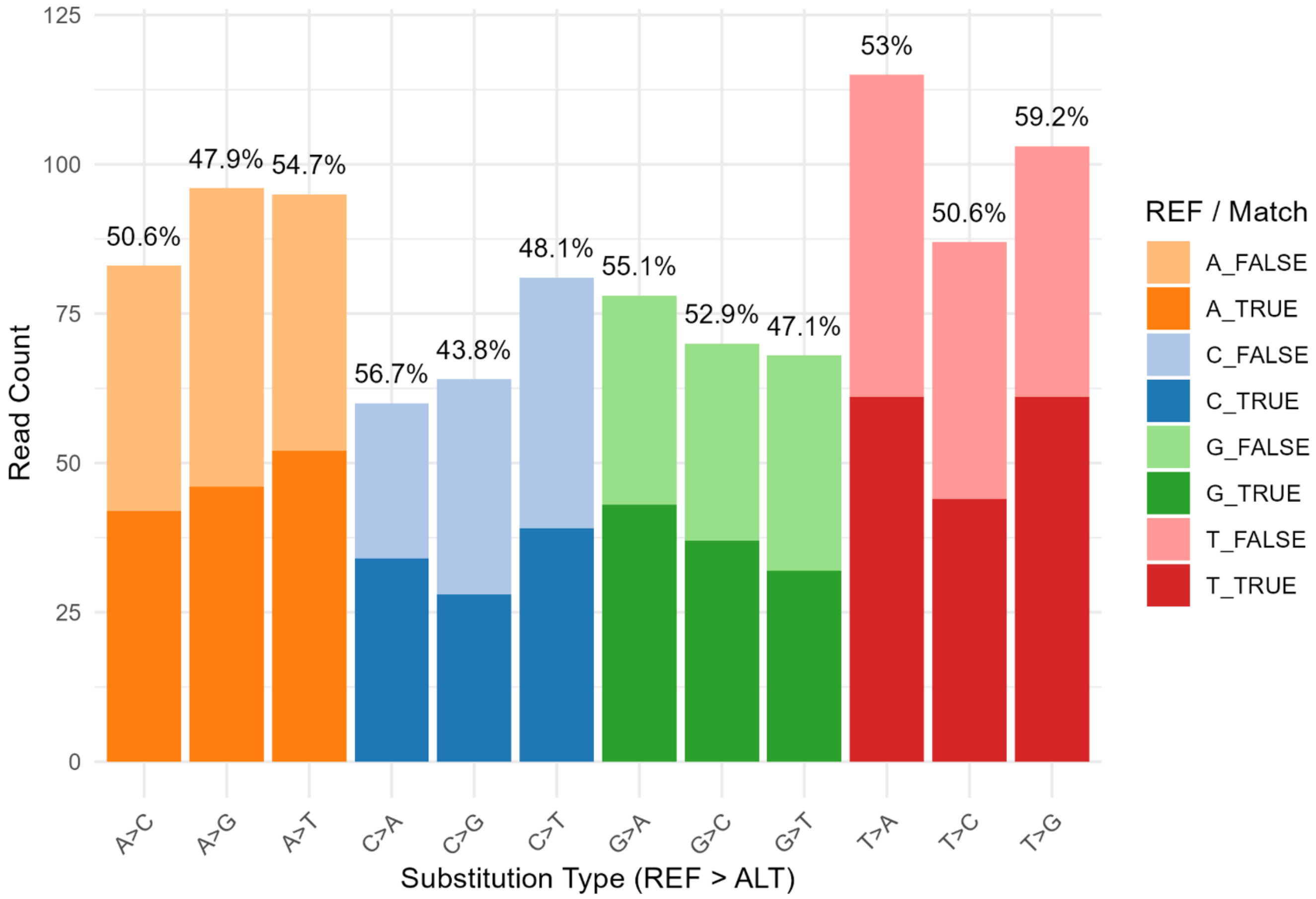


**Supplemental Figure S4.** Allele match distribution per base substitution color-coded by reference base. Percentages above the bars and dark-shaded colors represent true-positive locus and allele matches. Chi-Square test between true-positive percentages resulted in a p-value of 0.783.
