## Supplemental File S3 for "MosaicSim: A Novel Mosaic Variant Simulator Reveals Diminishing Returns of Ultra-High Coverage for Mosaic Variant Detection": mosaicsim.sh

```
#!/bin/bash

# Global variables to store amounts
SEED_TYKE=0
REFERENCE_TYKE=""
PREFIX_TYKE=""
BAM_TYKE=""
TYKE_FALG="0"
SPIKE_FLAG="0"
BASELINE_SPIKE=""
SPIKEIN_SPIKE=""
RATIO_SPIKE=""
OUTPUT_SPIKE=""

display_main_help() {
        echo "Simulation of Mosaic Variants in Sequencing Data"
        echo -e "\nUsage: mosicsim <pipline> [arguments]"
        echo -e "\npipline:"
        echo "  Tyke     Genrated simulated mosaic SV/SNV"
        echo "  Spike    Simulates a sample with potential mosiac variants at a user-specified ratio"
        exit 1
} 
display_Tyke_usage() {
    echo "Options:"
    echo "  -h, --help                     Display this help message"
    echo "  -b, --bam        <bam>         Input bam file"
    echo "  -p, --prefix     <prefix>      output prefix "
    echo "  -s, --seed       <seed>        seed number "
    echo "  -r, --reference  <REF>         reference fasta file"
    exit 1

}

# Function to display usage information for 'samtools Spike' command
display_Spike_usage() {
    echo "Options:"
    echo "  -h, --help             Display this help message"
    echo "  -b, --baseline         bam file to spike data into"
    echo "  -s, --spikein          bam file extract data from"
    echo "  -r, --ratio            spike-in ratio e.g 0.05 (5%)"
    echo "  -o, --output           path to output_dirpath for spikein results"
    exit 1  

}

# Function to parse options and set amounts
parse_options_tyke() {
    while [[ $# -gt 0 ]]; do
        case $1 in
            -b|--bam)
                BAM_TYKE=$2
                shift
                shift
                ;;
            -r|--reference)
                REFERENCE_TYKE=$2
                shift
                shift
                ;;
            -p|--prefix)
                PREFIX_TYKE=$2
                shift
                shift
                ;;
            -s|--seed)
                SEED_TYKE=$2
                shift
                shift
                ;;
            -h|--help)
                display_Tyke_usage
                shift
                shift
                ;;
            *)
                shift
                echo "Unknown1 option: $1"
                exit 1
                ;;
        esac
    done
}

parse_options_spike() {
    while [[ $# -gt 0 ]]; do
        case $1 in
            -b|--baseline)
                BASELINE_SPIKE=$2
                shift
                shift
                ;;
            -s|--spikein)
                SPIKEIN_SPIKE=$2
                shift
                shift
                ;;
            -r|--ratio )
                RATIO_SPIKE=$2
                shift
                shift
                ;;
            -o|--output )
                OUTPUT_SPIKE=$2
                shift
                shift
                ;;
            -h|--help)
                display_Spike_usage
                shift
                shift
                ;;
            *)
                shift
                echo "Unknown1 option: $1"
                exit 1
                ;;
        esac
    done
}

# Main function
main() {
    # Check if any arguments are provided
    if [ $# -eq 0 ]; then
	display_main_help
    fi

    # Parse the command and call respective function
    case $1 in
        tyke|Tyke)
            shift
            if [ $# -eq 0 ]
            then
                display_Tyke_usage
                exit
            fi
            parse_options_tyke "$@"
            TYKE_FALG="1"
            ;;
        spike|Spike)
            shift
            if [ $# -eq 0 ]
            then
                display_Spike_usage
                exit
            fi
            parse_options_spike "$@"
            SPIKE_FLAG=1
            ;;
        *)
            echo "Unknown command: $1"
            echo "Available commands: Tyke, Spike"
            exit 1
            ;;
    esac
}

# Call the main function with provided arguments
main "$@"
if [ $TYKE_FALG == "1" ]
then
    echo "TykeVar pipline"
    echo "${BAM_TYKE}" "${REFERENCE_TYKE}" "${PREFIX_TYKE}" "${SEED_TYKE}"
    python3 scripts/Tyke/TykeVarSimulator/TykeVarSimulator.py "${BAM_TYKE}" "${REFERENCE_TYKE}" "${PREFIX_TYKE}" "${SEED_TYKE}"
    python3 scripts/Tyke/main.py -v SIMULATED_VCF -b BAM -r REF -o OUTPUT_FASTQ
    python3 scripts/Tyke/filter_merge_bam.py -b BAM -m MOD_BAM --primary -o OUT_DIR --prefix PREFIX
    exit
fi 

if [ $SPIKE_FLAG == "1" ]
then
    echo "Spikein pipline"
    echo "$BASELINE_SPIKE" "$SPIKEIN_SPIKE" "$RATIO_SPIKE" "$OUTPUT_SPIKE"
fi
```
