## Supplemental File S3 for "MosaicSim: A Novel Mosaic Variant Simulator Reveals Diminishing Returns of Ultra-High Coverage for Mosaic Variant Detection": README.md

# MosaicSim: Simulation of Mosaic Variants in Sequencing Data

**Hackathon team: Lead: Fritz Sedlazeck - Developers: Erik Stricker, Xinchang Zheng, Michal Izydorczyk, Chi-Lam Poon, Philippe Sanio, Farhang Jaryani, Joyjit Daw, Divya Kalra, Adam Alexander - Writers: Erik Stricker, Sontosh Deb**

*We provide two simulation workflows which output sequencing read files with artificial mosaic variants and a ground truth mosaic variant annotation file for the validation of mosaic variant callers.*


## Table of Contents
|1. [Background](#background)<br>2. [Installation](#installation)<br>3. [Dependencies](#dependencies)<br>4. [Tests](#tests)<br>5. [How to Use It](#how-to-use-it)<br>6. [Example Implementation](#example-implementation)<br>7. [Method Description](#method-description)<br>8. [Contributers](#contributers)<br>9. [References](#references)<br><img width=1000/>|<img src="images/MosaicSim_workflows.png"  style="width: auto; display: block; margin: auto;">|
|:------|-:|


## Background

<img src="images/BackgroundMV.png"  style="width: auto; display: block; margin: auto;">

In the context of individual genome comparison, mutations that appear within a small fraction of the population are considered rare variants[<sup>1</sup>](#1). When assessing a population of cells from a tissue of the same individual in turn, rare variants only present in a small fraction of the cells are defined as mosaic variants (MVs)[<sup>2</sup>](#2). Recent studies have shown that there is potential disease associations of for certain MVs[<sup>2</sup>](#2). However, MVs are challenging to detect because they are mixed in with data from the non-mutated cells and present in the same sequencing file. Therefore, several pipelines have been developed or adjusted to extract mosaic single nucleotide, structural or indel variants from whole genome sequencing data such as Sniffles[<sup>3</sup>](#3), DeepMosaic[<sup>4</sup>](#4), Mutect2[<sup>5</sup>](#5), DeepVariant[<sup>6</sup>](#6). To benchmark and validate the efficiency and accuracy of these methods, sequencing files with known MVs are necessary. We developed two simulation workflows called SpikeVar (*Spike* in *Var*iants from a second individual) and TweakVar (*Tweak* *Var*iants within existing reads of one individual), which output sequencing read files with artificial MVs and a ground truth annotation file for the MVs. SpikeVar accomplishes this by spiking in real reads from a sample at user-defined ratio into the sequencing file from a second sample. In contrast, TweakVar creates a list of random mutations and modifies a fraction of existing reads to match the user-defined MV frequency.


## Installation
These instructions are valid for **Linux x86**. For other platforms (e.g. MacOS), instructions will need to be adapted.

Load conda, e.g.
```
module load miniconda/3
```
or
```
module load anaconda3 ##sometimes python has to be loaded before starting an environment)
module load python
```

Installing two conda environments with python 3.10
```
conda create -n MosaicSim python=3.10
conda init
conda activate MosaicSim
```

Unload python if previously loaded so that python 3.10 from conda environment will be used
```
module unload python
```

Obtain the MosaicSim from github (replace $HOME with your preferred installation director)
```
cd  $HOME
git clone https://github.com/erikstricker/MosaicSim.git
```

To install the relevant python dependencies, run
```
REPO_ROOT="$HOME/MosaicSim/"

pip install -r $REPO_ROOT/requirements.txt
```
Ensure to also load mosdepth>0.3.2 (for SpikeVar), samtools >1.15.1, and bcftools>1.19

Once the requirements are installed, please install or load the following additional packages

_Installation_
```
conda install -c bioconda samtools bcftools mosdepth
```
_Loading (e.g.)_
```
export PATH=/path/to/software/mosdepth/mosdepth-0.3.2/bin:$PATH
export PATH=/path/to/software/samtools/samtools-1.21/bin:$PATH
export PATH=/path/to/software/bcftools/bcftools-1.19/bin:$PATH
```
or
```
module load mosdepth-0.3.2
module load samtools-1.21
module load bcftools-1.19
```

## Dependencies

### SpikeVar
- mosdepth 0.3.2
- samtools 1.15.1
- bcftools 1.19
- Python 3.6.8
- bcftools
  
### TweakVar
- pysam (0.21.0) 
- numpy (1.25.2)
- biopython (1.81)
- samtools 1.15.1
- bcftools 1.19

## Contributers


|<img src="images/Erik Stricker.jpg" width="150"/><br>Erik Stricker|<img src="images/Chi-Lam Poon.jpg" width="150"/><br>Chi-Lam Poon|<img src="images/Philippe Sanio.jpg" width="150"/><br>Philippe Sanio|<img src="images/Xinchang Zheng.jpg" width="150"/><br>Xinchang Zheng|<img src="images/Farhang Jaryani.png" width="150"/><br>Farhang Jaryani|
|:-:|:-:|:-:|:-:|:-:|

|<img src="images/Joyjit Daw.png" width="150"/><br>Joyjit Daw |<img src="images/Michal Bogumil Izydorczyk.png" width="150"/><br>Michal Izydorczyk|<img src="images/Sontosh K Deb.jpg" width="150"/><br>Sontosh Deb|<img src="images/Fritz Sedlazeck.jpg" width="150"/><br>Fritz Sedlazeck |<img src="images/Alexander Adam.jpg" width="150"/><br>Adam Alexander|
|:-:|:-:|:-:|:-:|:-:|

|<img src="images/Divya Kalrai_placeholder.jpg" width="150"/><br>Divya Kalra|
|:-:|


## References

<a id="1">[1]</a> Sariya S, Lee JH, Mayeux R, Vardarajan BN, Reyes-Dumeyer D, Manly JJ, Brickman AM, Lantigua R, Medrano M, Jimenez-Velazquez IZ, Tosto G. Rare Variants Imputation in Admixed Populations: Comparison Across Reference Panels and Bioinformatics Tools. Front Genet. 2019;10:239. Epub 20190403. doi: 10.3389/fgene.2019.00239. PubMed PMID: 31001313; PMCID: PMC6456789.  

<a id="2">[2]</a> Miller CR, Lee K, Pfau RB, Reshmi SC, Corsmeier DJ, Hashimoto S, Dave-Wala A, Jayaraman V, Koboldt D, Matthews T, Mouhlas D, Stein M, McKinney A, Grossman T, Kelly BJ, White P, Magrini V, Wilson RK, Mardis ER, Cottrell CE. Disease-associated mosaic variation in clinical exome sequencing: a two-year pediatric tertiary care experience. Cold Spring Harb Mol Case Stud. 2020;6(3). Epub 20200612. doi: 10.1101/mcs.a005231. PubMed PMID: 32371413; PMCID: PMC7304353.  

<a id="3">[3]</a> Sedlazeck FJ, Rescheneder P, Smolka M, Fang H, Nattestad M, von Haeseler A, Schatz MC. Accurate detection of complex structural variations using single-molecule sequencing. Nat Methods. 2018;15(6):461-8. Epub 20180430. doi: 10.1038/s41592-018-0001-7. PubMed PMID: 29713083; PMCID: PMC5990442.  

<a id="4">[4]</a> Yang X, Xu X, Breuss MW, Antaki D, Ball LL, Chung C, Shen J, Li C, George RD, Wang Y, Bae T, Cheng Y, Abyzov A, Wei L, Alexandrov LB, Sebat JL, Network NBSM, Gleeson JG. Control-independent mosaic single nucleotide variant detection with DeepMosaic. Nat Biotechnol. 2023;41(6):870-7. Epub 20230102. doi: 10.1038/s41587-022-01559-w. PubMed PMID: 36593400; PMCID: PMC10314968.  

<a id="5">[5]</a> McKenna A, Hanna M, Banks E, Sivachenko A, Cibulskis K, Kernytsky A, Garimella K, Altshuler D, Gabriel S, Daly M, DePristo MA. The Genome Analysis Toolkit: a MapReduce framework for analyzing next-generation DNA sequencing data. Genome Res. 2010;20(9):1297-303. Epub 20100719. doi: 10.1101/gr.107524.110. PubMed PMID: 20644199; PMCID: PMC2928508.  

<a id="6">[6]</a> Poplin R, Chang PC, Alexander D, Schwartz S, Colthurst T, Ku A, Newburger D, Dijamco J, Nguyen N, Afshar PT, Gross SS, Dorfman L, McLean CY, DePristo MA. A universal SNP and small-indel variant caller using deep neural networks. Nat Biotechnol. 2018;36(10):983-7. Epub 20180924. doi: 10.1038/nbt.4235. PubMed PMID: 30247488.
