## Supplemental File S3 for "MosaicSim: A Novel Mosaic Variant Simulator Reveals Diminishing Returns of Ultra-High Coverage for Mosaic Variant Detection": 2b_re-genotyping_main.sh

```
#!/bin/bash

# Input variables are: VARIANT, VAF, VCF_1, VCF_2, MOD_BAM, OUTPUT_DIR, READ_LENGTH
# VARIANT can be either SNV or SV
# READ_LENGTH can be either SHORT or LONG for SVs

# Example command:
# For SNV:
# ./2b_re-genotyping_main.sh SNV 0.1 /path/to/vcf1.vcf /path/to/vcf2.vcf /path/to/mod.bam /path/to/output_dir
# or 
# For SV:
# ./2b_re-genotyping_main.sh SV 0.1 /path/to/vcf1.vcf /path/to/vcf2.vcf /path/to/mod.bam /path/to/output_dir SHORT
# ./2b_re-genotyping_main.sh SV 0.1 /path/to/vcf1.vcf /path/to/vcf2.vcf /path/to/mod.bam /path/to/output_dir LONG

VARIANT=$1
VAF=$2
VCF_1=$3
VCF_2=$4
MOD_BAM=$5
OUTPUT_DIR=$6
READ_LENGTH=$7
MERGED_VCF_BCFTOOLS="${OUTPUT_DIR}/merged_bcftools.vcf.gz"
MERGED_VCF="${OUTPUT_DIR}/merged.vcf"

OUTPUT_VCF="${OUTPUT_DIR}/output_regenotyped.vcf"
OUTPUT_VCF_FILTERED="${OUTPUT_DIR}/output_genotypes_filtered.vcf"
REFERENCE=${8}

mkdir -p $OUTPUT_DIR

if [ "$VARIANT" = "SNV" ]; then
    echo "Input is 'SNV'"
    ./2b_SNV.sh $READ_LENGTH $VAF $MERGED_VCF $MOD_BAM $OUTPUT_VCF $REFERENCE
elif [ "$VARIANT" = "SV" ]; then
    echo "Input is 'SV'"
    ./2b_SV.sh $READ_LENGTH $VAF $VCF_1 $VCF_2 $MOD_BAM $OUTPUT_DIR $REFERENCE "Spike_out"
else
    echo "Error: Invalid input"
fi
```
