## Supplemental File S3 for "MosaicSim: A Novel Mosaic Variant Simulator Reveals Diminishing Returns of Ultra-High Coverage for Mosaic Variant Detection": 2b_SNV.sh

```
#!/bin/bash

# Input vairables are: READ_LENGTH, VAF, MERGED_VCF, MOD_BAM and OUTPUT_DIR
# READ_LENGTH can be either SHORT or LONG

READ_LENGTH=$1
VAF=$2
MERGED_VCF=$3
MOD_BAM=$4
OUTPUT_VCF=$5
REFERENCE=$6
# activate conda env which uses python 3.6 called mosaicSim
conda activate mosaicSim
# call samtools mpileup to genotypes the variants
#samtools mpileup $MOD_BAM | bcftools call -mv -Oz -o $OUTPUT_VCF
bcftools  mpileup  -Ou -f $REFERENCE $MOD_BAM  |  bcftools call -mv -Oz -o $OUTPUT_VCF
```
