## Supplemental File S3 for "MosaicSim: A Novel Mosaic Variant Simulator Reveals Diminishing Returns of Ultra-High Coverage for Mosaic Variant Detection": 2b_SV.sh

```
#!/bin/bash

# Input vairables are: READ_LENGTH, VAF, MERGED_VCF, MOD_BAM and OUTPUT_DIR
# Optional input variables for short_paragraph mode:
# 1. REFERENCE : reference for MOD_BAM in fasta format
# 2. OUTPUT_PFX : output prefix for paragraph
# READ_LENGTH can be either SHORT or LONG or short_paragraph

READ_LENGTH=$1
VAF=$2
VCF_1=$3
VCF_2=$4
#MERGED_VCF=$3
MOD_BAM=$5
OUTPUT_DIR=$6
REFERENCE=$7
OUTPUT_PFX=$8

echo "VCF_1: $VCF_1"
echo "VCF_2: $VCF_2"
echo "MOD_BAM: $MOD_BAM"

if [ "$READ_LENGTH" = "short" ]; then
    
    # Run SV regenotyper for short reads
    ./2b_sv_short.sh $VCF_1 $MOD_BAM $OUTPUT_DIR $REFERENCE
    ./2b_sv_short.sh $VCF_2 $MOD_BAM $OUTPUT_DIR $REFERENCE
elif [ "$READ_LENGTH" = "long" ]; then
    echo "Sniffles 1"
    ./2b_sv_long.sh $VCF_1 $MOD_BAM "$OUTPUT_DIR/sample1.regenotyped.vcf.gz"
    ./2b_vaf_filter.sh "$OUTPUT_DIR/sample1.regenotyped.vcf.gz" $VAF "$OUTPUT_DIR/sample1.regenotyped.filtered.vcf"
    
    echo "Sniffles 2"
    ./2b_sv_long.sh $VCF_2 $MOD_BAM "$OUTPUT_DIR/sample2.regenotyped.vcf.gz"
    ./2b_vaf_filter.sh "$OUTPUT_DIR/sample2.regenotyped.vcf.gz" $VAF "$OUTPUT_DIR/sample2.regenotyped.filtered.vcf"

    # Do merging of the two filtered VCF files
    ./2b_vcf_merge.sh "$OUTPUT_DIR/sample1.regenotyped.filtered.vcf.gz" "$OUTPUT_DIR/sample2.regenotyped.filtered.vcf.gz" "$OUTPUT_DIR/output_regenotyped.vcf"
    #./2b_vcf_merge.sh "$OUTPUT_DIR/sample1.regenotyped.vcf.gz" "$OUTPUT_DIR/sample2.regenotyped.vcf.gz" "$OUTPUT_DIR/output_regenotyped.vcf"
else
    echo "Error: Invalid input"
fi
```
