## Supplemental File S3 for "MosaicSim: A Novel Mosaic Variant Simulator Reveals Diminishing Returns of Ultra-High Coverage for Mosaic Variant Detection": 2b_sv_long.sh

```
#!/bin/bash

VCF_FILE=$1
BAM_FILE=$2
OUTPUT_FILE=$3

# Run sniffles
echo "====== Running sniffles ======"
echo "VCF_FILE: $VCF_FILE"
echo "BAM_FILE: $BAM_FILE"
echo "OUTPUT_FILE: $OUTPUT_FILE"

sniffles --input $BAM_FILE --genotype-vcf $VCF_FILE --sample-id mosaicSim --vcf $OUTPUT_FILE
```
