## Supplemental File S3 for "MosaicSim: A Novel Mosaic Variant Simulator Reveals Diminishing Returns of Ultra-High Coverage for Mosaic Variant Detection": 2b_vaf_filter.sh

```
#!/bin/bash
OUTPUT_VCF=$1
VAF=$2
OUTPUT_VCF_FILTERED=$3

echo "===== Filtering ======"
echo "OUTPUT_VCF: $OUTPUT_VCF"
echo "VAF: $VAF"
echo "OUTPUT_VCF_FILTERED: $OUTPUT_VCF_FILTERED"

python vaf_seperator.py $OUTPUT_VCF $VAF $OUTPUT_VCF_FILTERED
bgzip -f $OUTPUT_VCF_FILTERED
tabix -f $OUTPUT_VCF_FILTERED.gz
```
