## Supplemental File S3 for "MosaicSim: A Novel Mosaic Variant Simulator Reveals Diminishing Returns of Ultra-High Coverage for Mosaic Variant Detection": 2b_vcf_merge.sh

```
#!/bin/bash

VCF_1=$1
VCF_2=$2
OUTPUT_VCF=$3

echo "===== Merging ======"
echo "VCF_1: $VCF_1"
echo "VCF_2: $VCF_2"
echo "OUTPUT_VCF: $OUTPUT_VCF.tmp.vcf"
echo "End OUTPUT_VCF: $OUTPUT_VCF"

# Merge the two VCF files with bcftools
echo "Merging: bcftools merge -m none --force-samples -o $OUTPUT_VCF.tmp.vcf $VCF_1 $VCF_2"
bcftools merge -m none --force-samples -o $OUTPUT_VCF.tmp.vcf $VCF_1 $VCF_2

bgzip -f "$OUTPUT_VCF.tmp.vcf"
tabix -f "$OUTPUT_VCF.tmp.vcf.gz"

# Run truvari collapse to get ride of all variants which match the thresholds

truvari collapse -i "$OUTPUT_VCF.tmp.vcf.gz" -o "$OUTPUT_VCF"
bgzip -f "$OUTPUT_VCF"
tabix -f "$OUTPUT_VCF.gz"

# Clean up
rm "$OUTPUT_VCF.tmp.vcf.gz"
rm "$OUTPUT_VCF.tmp.vcf.gz.tbi"
```
