## Supplemental File S3 for "MosaicSim: A Novel Mosaic Variant Simulator Reveals Diminishing Returns of Ultra-High Coverage for Mosaic Variant Detection": calculate_ratio.py

```
#!/usr/bin/env python3

import sys

from pathlib import Path

spikein_ratio = sys.argv[1]
baseline_sample_cov = sys.argv[2]
spikein_sample_cov = sys.argv[3]

baseline_sample_ratio = 1.00 - float(spikein_ratio)
spikein_sample_ratio = (float(baseline_sample_cov) * float(spikein_ratio)) / float(spikein_sample_cov)

print(baseline_sample_ratio, spikein_sample_ratio)
```
