## Supplemental File S3 for "MosaicSim: A Novel Mosaic Variant Simulator Reveals Diminishing Returns of Ultra-High Coverage for Mosaic Variant Detection": README.md

# SpikeVar: Simulation of Mosaic Variants by Combination of Sequencing Data From Two Individuals

**Hackathon team: Lead: Fritz Sedlazeck - Developers: Erik Stricker, Xinchang Zheng, Michal Izydorczyk, Chi-Lam Poon, Philippe Sanio, Farhang Jaryani, Joyjit Daw, Divya Kalra, Adam Alexander - Writers: Erik Stricker, Sontosh Deb**

*We provide two simulation workflows which output sequencing read files with artificial mosaic variants and a ground truth mosaic variant annotation file for the validation of mosaic variant callers.*


## Table of Contents
|1. [Background](#background)<br>2. [Installation](#installation)<br>3. [Dependencies](#dependencies)<br>4. [Tests](#tests)<br>5. [How to Use It](#how-to-use-it)<br>6. [Example Implementation](#example-implementation)<br>7. [Method Description](#method-description)<br>8. [Contributers](#contributers)<br>9. [References](#references)<br><img width=1000/>|<img src="https://github.com/erikstricker/MosaicSim/blob/93ae22dd82122271b36fc1b585e283c59a3f4795/images/SpikeVar_workflow_simple.png"  style="width: auto; display: block; margin: auto;">|
|:------|-:|


## Background

<img src="https://github.com/erikstricker/MosaicSim/blob/93ae22dd82122271b36fc1b585e283c59a3f4795/images/BackgroundMV.png"  style="width: auto; display: block; margin: auto;">

In the context of individual genome comparison, mutations that appear within a small fraction of the population are considered rare variants[<sup>1</sup>](#1). When assessing a population of cells from a tissue of the same individual in turn, rare variants only present in a small fraction of the cells are defined as mosaic variants (MVs)[<sup>2</sup>](#2). Recent studies have shown that there is potential disease associations of for certain MVs[<sup>2</sup>](#2). However, MVs are challenging to detect because they are mixed in with data from the non-mutated cells and present in the same sequencing file. Therefore, several pipelines have been developed or adjusted to extract mosaic single nucleotide, structural or indel variants from whole genome sequencing data such as Sniffles[<sup>3</sup>](#3), DeepMosaic[<sup>4</sup>](#4), Mutect2[<sup>5</sup>](#5), DeepVariant[<sup>6</sup>](#6). To benchmark and validate the efficiency and accuracy of these methods, sequencing files with known MVs are necessary. We developed two simulation workflows called SpikeVar (*Spike* in *Var*iants from a second individual) and TweakVar (*Tweak* *Var*iants within existing reads of one individual), which output sequencing read files with artificial MVs and a ground truth annotation file for the MVs. SpikeVar accomplishes this by spiking in real reads from a sample at user-defined ratio into the sequencing file from a second sample. In contrast, TweakVar creates a list of random mutations and modifies a fraction of existing reads to match the user-defined MV frequency.


## Installation
These instructions are valid for **Linux x86**. For other platforms (e.g. MacOS), instructions will need to be adapted.

Load conda, e.g.
```
module load miniconda/3
```
or
```
module load anaconda3 ##sometimes python has to be loaded before starting an environment)
module load python
```

Installing two conda environments with python 3.10
```
conda create -n MosaicSim python=3.10
conda init
conda activate MosaicSim
```

Unload python if previously loaded so that python 3.10 from conda environment will be used
```
module unload python
```

Obtain the MosaicSim from github (replace $HOME with your preferred installation director)
```
cd  $HOME
git clone https://github.com/erikstricker/MosaicSim.git
```

To install the relevant python dependencies, run
```
REPO_ROOT="$HOME/MosaicSim/"

pip install -r $REPO_ROOT/requirements.txt
```
Ensure to also load mosdepth>0.3.2 (for SpikeVar), samtools >1.15.1, and bcftools>1.19

Once the requirements are installed, please install or load the following additional packages

_Installation_
```
conda install -c bioconda samtools bcftools mosdepth
```
_Loading (e.g.)_
```
export PATH=/path/to/software/mosdepth/mosdepth-0.3.2/bin:$PATH
export PATH=/path/to/software/samtools/samtools-1.21/bin:$PATH
export PATH=/path/to/software/bcftools/bcftools-1.19/bin:$PATH
```
or
```
module load mosdepth-0.3.2
module load samtools-1.21
module load bcftools-1.19
```

## Dependencies

### SpikeVar
- mosdepth 0.3.2
- samtools 1.15.1
- bcftools 1.19
- Python 3.6.8
- bcftools
  
```

## How to Use It

### SpikeVar

The spiked-in dataset simulates a sample with potential mosiac variants at a user-specified ratio. The re-genotyped VCFs of the samples and the VCF of the spiked-in dataset can be compared to evaluate AF < user-specified value.

#### 1) SpikeVarDatabaseCreator - Generate Spiked-in Dataset

<img src="https://github.com/erikstricker/MosaicSim/blob/93ae22dd82122271b36fc1b585e283c59a3f4795/images/SpikeVarDatabaseCreator.png"  height="150" align="right">  

In this step, x% of mutations are strategically introduced from sample A to sample B. Both datasets are down-sampled and then merged to create a mixed dataset that represents a sequence read dataset with mosaic variants, including structural variations (SVs), single nucleotide variations (SNVs), and insertions/deletions (indels). 

```
./spike-in.sh <path to sampleA.bam> <path to sampleB.bam> <spike-in ratio x/100> <path to samtools binary> <path to mosdepth binary> <output dirpath> <path to script calculate_ratio.py>
```
#### 2) SpikeVarReporter - Filter Reads After Variant Allele Frequency Recalculation

<img src="https://github.com/erikstricker/MosaicSim/blob/93ae22dd82122271b36fc1b585e283c59a3f4795/images/SpikeVarReporter.png"  height="250" align="right">  

After creating the modified BAM file we have to re-calculate the variant allele frequency (VAF) for all variants.
First, all variants stored from both VCF files must be merged and the VAF must be recalculated. 
Depending on the variants we either start a SNV or SV caller, which can recalculate the VAF of each variant. 
For SNVs, we are using bcftools mpileup. For SVs and short read data we are using Paragraph from Illumina and for long read data Sniffles2 is used.

Last the re-genotyped VCF is filtered according to the VAF with a small Python script, which calculates the minor allele frequency (MAF) for each variant and lets a variant pass to the final output if the MAF is equal or greater than the use specified VAF.
```
./2b_re-genotyping_main.sh <VARIANT> <VAF> <sampleA.vcf> <sampleB.vcf> <sampleAandB.bam> output/path <ref.fa> <short|long>
```
#### 3) Run Your Favorite Mosaic Variant Caller and Compare Results

## Example Implementation

### SpikeVar

Here, we use the SpikeVar workflow to automatically spike in sample HG002 at a 5% concentration into sample HG0733, to result in a 5% mosaic variant allele frequency (VAF). A downside is that the generated mixed .bam file will include 4 haplotype structures which cannot be corrected for. Furthermore, certain variants (e.g. HG002 variants) will not be presented at the targeted VAF. For example, heterozygous variants will not be represented by 5% VAF but rather at ~2.5% VAF. To account for this we re-genotype variants and report only variants that should be identifiable at the user-defined threshold or higher VAF.   

#### 1) Fetch Data
In order to spike in sample B into sample A, the pipeline first needs an initial set of aligned reads. We used HG002 and HG00733 datasets.

Reads - `ftp://ftp-trace.ncbi.nlm.nih.gov/ReferenceSamples/giab/data/AshkenazimTrio/HG002_NA24385_son/Ultralong_OxfordNanopore/guppy-V3.2.4_2020-01-22/HG002_hs37d5_ONT-UL_GIAB_20200122.phased.bam` and `ftp://ftp-trace.ncbi.nlm.nih.gov/ReferenceSamples/giab/data/AshkenazimTrio/HG002_NA24385_son/Ultralong_OxfordNanopore/guppy-V3.2.4_2020-01-22/HG002_hs37d5_ONT-UL_GIAB_20200122.phased.bam.bai`


#### 2) SpikeVarDatabaseCreator - Generate Spike-in Dataset
We spiked 5% reads from HG0733 to HG002 for the next part of the workflow.
```
./spike-in.sh HG002_hs37d5_ONT-UL_GIAB_20200122.phased.bam HG007733.bam 0.05 /software/bin/samtools /software/bin/mosdepth /output `pwd`/calculate_ratio.py
```
#### 3) SpikeVarReporter - Filter Reads with >5% Variant Allele Frequency After Recalculation
After creating the new BAM file from e.g. HG002 and HG00733, we have to re-calculate the variant allele frequency (VAF) for all variants.
First we merge both VCF files from e.g. HG002 and HG00733 with bcftools. Depending on the variants we either start a SNV or SV caller, which can recalculate the VAF of each variant. 
For SNVs we are using bcftools mpileup. For SVs and short read data we are using Paragraph from Illumina and for long read data Sniffles2 is used.

```
./2b_re-genotyping_main.sh SV 0.05 HG002_SV.Tier1.vcf HG00733_SV.Tier1.vcf /output/SPIKED.BAM ./ LONG hs37d5.fa
```

#### 4) Run Your Favorite Mosaic Variant Caller

Run your choice of mosaic variant caller on the modified `HG002_ONT_hg37_chr5_HG00733_ONT_hg37_chr5_merged.sorted.bam` file and compare the results with the truth set `output_genotypes_filtered.vcf` file.

#### 5) Results

Your spiked in reads are now visible in the IGV genome browser. 
 
<img src="https://github.com/erikstricker/MosaicSim/blob/93ae22dd82122271b36fc1b585e283c59a3f4795/images/Spike_screenshot_sv.png"  align="center"> 
<p align="center">
<b>Example f a spiked in deletion.</b>
</p>
<br />

<img src="https://github.com/erikstricker/MosaicSim/blob/93ae22dd82122271b36fc1b585e283c59a3f4795/images/Spike_screenshot_sv2_ins.png"  align="center">
<p align="center">
<b>Example of a spiked in inserton.</b>
</p>  

## Method Description 

### SpikeVar - Generation of Sequencing Data With a Low Frequencing of Reads From Another Sample
<img src="https://github.com/erikstricker/MosaicSim/blob/93ae22dd82122271b36fc1b585e283c59a3f4795/images/SpikeVarflowchart_updated.png" width="500"/>
<p align="justify">
<b>SpikeVar workflow, with major steps to assess the sensitivity and accuracy of the mosaic variant callers. (A, B: individual samples, A/B: merged samples, .bam and .vcf: input and output file formats in different steps, Black header boxes: tool or file names, Green header boxes: simulated files or final files used for validation comparisons)</b>
</p>  

<br />
The SpikeVar workflow outputs a mixed sequencing read dataset in .bam format containing reads from one dominant sample and reads from another sample spiked in at a user-defined ratio corresponding to the simulated mosaic variant allele frequency (VAF) together with a .vcf file annotating the confirmed mosaic variant locations within the mixed dataset. The SpikeVarDatasetCreator takes aligned sequencing reads from sample A and sample B as the initial input. In this step, a spike-in methodology is applied to strategically introduce x% of mutations from one sample to another using <insert tool>. Accordingly, sample A is first down-sampled to retain 100-x% of its original reads, then sample B is down-sampled to x% considering the coverage differences between the samples. Using <insert tool>, both down-sampled datasets are then merged to create a mixed dataset that represents a sequence read dataset with mosaic variants, including structural variations (SVs), single nucleotide variations (SNVs), and insertions/deletions (indels).  

The SpikeVarReporter then determines VAFs for each variant in the mixed dataset using <insert tool> based on the mixed variant locations derived by merging the .vcf files from sample A and sample B using <insert tool>. Variants with VAFs exceeding or equal to the introduced mutations (i.e., x%) are then selected to create a truth set for benchmarking using <insert tool>.  
 
To assess a mosaic variant caller’s sensitivity and accuracy, the same mixed dataset is used to call mosaic variants. The output mosaic variant locations and VAFs are then compared to the truth set for validation.  

## Contributers


|<img src="https://github.com/erikstricker/MosaicSim/blob/93ae22dd82122271b36fc1b585e283c59a3f4795/images/Erik Stricker.jpg" width="150"/><br>Erik Stricker|<img src="https://github.com/erikstricker/MosaicSim/blob/93ae22dd82122271b36fc1b585e283c59a3f4795/images/Chi-Lam Poon.jpg" width="150"/><br>Chi-Lam Poon|<img src="https://github.com/erikstricker/MosaicSim/blob/93ae22dd82122271b36fc1b585e283c59a3f4795/images/Philippe Sanio.jpg" width="150"/><br>Philippe Sanio|<img src="https://github.com/erikstricker/MosaicSim/blob/93ae22dd82122271b36fc1b585e283c59a3f4795/images/Xinchang Zheng.jpg" width="150"/><br>Xinchang Zheng|<img src="https://github.com/erikstricker/MosaicSim/blob/93ae22dd82122271b36fc1b585e283c59a3f4795/images/Farhang Jaryani.png" width="150"/><br>Farhang Jaryani|
|:-:|:-:|:-:|:-:|:-:|

|<img src="https://github.com/erikstricker/MosaicSim/blob/93ae22dd82122271b36fc1b585e283c59a3f4795/images/Joyjit Daw.png" width="150"/><br>Joyjit Daw |<img src="https://github.com/erikstricker/MosaicSim/blob/93ae22dd82122271b36fc1b585e283c59a3f4795/images/Michal Bogumil Izydorczyk.png" width="150"/><br>Michal Izydorczyk|<img src="https://github.com/erikstricker/MosaicSim/blob/93ae22dd82122271b36fc1b585e283c59a3f4795/images/Sontosh K Deb.jpg" width="150"/><br>Sontosh Deb|<img src="https://github.com/erikstricker/MosaicSim/blob/93ae22dd82122271b36fc1b585e283c59a3f4795/images/Fritz Sedlazeck.jpg" width="150"/><br>Fritz Sedlazeck |<img src="https://github.com/erikstricker/MosaicSim/blob/93ae22dd82122271b36fc1b585e283c59a3f4795/images/Alexander Adam.jpg" width="150"/><br>Adam Alexander|
|:-:|:-:|:-:|:-:|:-:|

|<img src="https://github.com/erikstricker/MosaicSim/blob/93ae22dd82122271b36fc1b585e283c59a3f4795/images/Divya Kalrai_placeholder.jpg" width="150"/><br>Divya Kalra|
|:-:|


## References

<a id="1">[1]</a> Sariya S, Lee JH, Mayeux R, Vardarajan BN, Reyes-Dumeyer D, Manly JJ, Brickman AM, Lantigua R, Medrano M, Jimenez-Velazquez IZ, Tosto G. Rare Variants Imputation in Admixed Populations: Comparison Across Reference Panels and Bioinformatics Tools. Front Genet. 2019;10:239. Epub 20190403. doi: 10.3389/fgene.2019.00239. PubMed PMID: 31001313; PMCID: PMC6456789.  

<a id="2">[2]</a> Miller CR, Lee K, Pfau RB, Reshmi SC, Corsmeier DJ, Hashimoto S, Dave-Wala A, Jayaraman V, Koboldt D, Matthews T, Mouhlas D, Stein M, McKinney A, Grossman T, Kelly BJ, White P, Magrini V, Wilson RK, Mardis ER, Cottrell CE. Disease-associated mosaic variation in clinical exome sequencing: a two-year pediatric tertiary care experience. Cold Spring Harb Mol Case Stud. 2020;6(3). Epub 20200612. doi: 10.1101/mcs.a005231. PubMed PMID: 32371413; PMCID: PMC7304353.  

<a id="3">[3]</a> Sedlazeck FJ, Rescheneder P, Smolka M, Fang H, Nattestad M, von Haeseler A, Schatz MC. Accurate detection of complex structural variations using single-molecule sequencing. Nat Methods. 2018;15(6):461-8. Epub 20180430. doi: 10.1038/s41592-018-0001-7. PubMed PMID: 29713083; PMCID: PMC5990442.  

<a id="4">[4]</a> Yang X, Xu X, Breuss MW, Antaki D, Ball LL, Chung C, Shen J, Li C, George RD, Wang Y, Bae T, Cheng Y, Abyzov A, Wei L, Alexandrov LB, Sebat JL, Network NBSM, Gleeson JG. Control-independent mosaic single nucleotide variant detection with DeepMosaic. Nat Biotechnol. 2023;41(6):870-7. Epub 20230102. doi: 10.1038/s41587-022-01559-w. PubMed PMID: 36593400; PMCID: PMC10314968.  

<a id="5">[5]</a> McKenna A, Hanna M, Banks E, Sivachenko A, Cibulskis K, Kernytsky A, Garimella K, Altshuler D, Gabriel S, Daly M, DePristo MA. The Genome Analysis Toolkit: a MapReduce framework for analyzing next-generation DNA sequencing data. Genome Res. 2010;20(9):1297-303. Epub 20100719. doi: 10.1101/gr.107524.110. PubMed PMID: 20644199; PMCID: PMC2928508.  

<a id="6">[6]</a> Poplin R, Chang PC, Alexander D, Schwartz S, Colthurst T, Ku A, Newburger D, Dijamco J, Nguyen N, Afshar PT, Gross SS, Dorfman L, McLean CY, DePristo MA. A universal SNP and small-indel variant caller using deep neural networks. Nat Biotechnol. 2018;36(10):983-7. Epub 20180924. doi: 10.1038/nbt.4235. PubMed PMID: 30247488.
