## Supplemental File S3 for "MosaicSim: A Novel Mosaic Variant Simulator Reveals Diminishing Returns of Ultra-High Coverage for Mosaic Variant Detection": run_calculate_ratio.sh

```
#!/bin/bash

# params
baseline_prefix=$1
spikein_prefix=$2
output_dirpath=$3
ratio=$4
calcratio_script_path=$5

baseline_cov=$(grep ^total ${output_dirpath}/${baseline_prefix}.mosdepth.summary.txt | awk '{print $4}')
spikein_cov=$(grep ^total ${output_dirpath}/${spikein_prefix}.mosdepth.summary.txt | awk '{print $4}')

python3 ${calcratio_script_path} ${ratio} ${baseline_cov} ${spikein_cov} | while read baseline spikein
do
    echo ${baseline} > ${output_dirpath}/${baseline_prefix}.ratio.txt
    echo ${spikein} > ${output_dirpath}/${spikein_prefix}.ratio.txt
done
```
