## Supplemental File S3 for "MosaicSim: A Novel Mosaic Variant Simulator Reveals Diminishing Returns of Ultra-High Coverage for Mosaic Variant Detection": run_merge.sh

```
#!/bin/bash

# params
baseline_prefix=$1
spikein_prefix=$2
samtools_path=$3
output_dirpath=$4

threads=10

baseline_ratio=$(cat ${output_dirpath}/${baseline_prefix}.ratio.txt)
spikein_ratio=$(cat ${output_dirpath}/${spikein_prefix}.ratio.txt)

baseline_subsample_bampath=${output_dirpath}/${baseline_prefix}"_"${baseline_ratio}.bam
spikein_subsample_bampath=${output_dirpath}/${spikein_prefix}"_"${spikein_ratio}.bam
            
${samtools_path} merge -r -o ${output_dirpath}/${baseline_prefix}"_"${spikein_prefix}"_merged.bam" -@ ${threads} ${baseline_subsample_bampath} ${spikein_subsample_bampath}
${samtools_path} sort -o ${output_dirpath}/${baseline_prefix}"_"${spikein_prefix}"_merged.sorted.bam" -@ ${threads} ${output_dirpath}/${baseline_prefix}"_"${spikein_prefix}"_merged.bam"
${samtools_path} index ${output_dirpath}/${baseline_prefix}"_"${spikein_prefix}"_merged.sorted.bam"
```
