## Supplemental File S3 for "MosaicSim: A Novel Mosaic Variant Simulator Reveals Diminishing Returns of Ultra-High Coverage for Mosaic Variant Detection": run_subsample.sh

```
#!/bin/bash

# params
bampath=$1
samtools_path=$2
output_dirpath=$3

threads=10

prefix=`basename ${bampath} .bam`

ratio=$(cat ${output_dirpath}/${prefix}.ratio.txt)
${samtools_path} view -h -@ ${threads} -s ${ratio} -b -o ${output_dirpath}/${prefix}"_"${ratio}.bam ${bampath}
${samtools_path} index ${output_dirpath}/${prefix}"_"${ratio}.bam
```
