## Supplemental File S3 for "MosaicSim: A Novel Mosaic Variant Simulator Reveals Diminishing Returns of Ultra-High Coverage for Mosaic Variant Detection": spike-in.sh

```
#!/bin/sh

# downsample data based on user-defined ratio
# spike-in into another sample

# params
baseline_bampath=$1   # main sample to spike data into
spikein_bampath=$2   # sample to extract data from
ratio=$3   # spike-in ratio e.g 0.05 (5%)
samtools_path=$4   # path to binary
mosdepth_path=$5   # path to binary
output_dirpath=$6
calcratio_script_path=$7  # path to script

threads=10
baseline_prefix=`basename ${baseline_bampath} .bam`
spikein_prefix=`basename ${spikein_bampath} .bam`

# check coverage of input data
baseline_mosdepth_submission=$(sbatch --chdir ${output_dirpath} --mail-type FAIL \
                                      --error baseline-mosdepth.err --output baseline-mosdepth.out \
                                      --job-name mosdepth --tasks-per-node 8 --mem 16gb --time 3:00:00 \
                                      --partition medium --account proj-fs0002 \
                                      run_mosdepth.sh ${baseline_bampath} \
                                      ${baseline_prefix} ${mosdepth_path}
                            )
spikein_mosdepth_submission=$(sbatch --chdir ${output_dirpath} --mail-type FAIL \
                                     --error spikein-mosdepth.err --output spikein-mosdepth.out \
                                     --job-name mosdepth --tasks-per-node 8 --mem 16gb	--time 3:00:00 \
                                     --partition medium --account proj-fs0002 \
                                     run_mosdepth.sh ${spikein_bampath} \
                                      ${spikein_prefix} ${mosdepth_path}
                           )
echo `date` "- INFO: submitting mosdepth for baseline and spikein samples"

# calculate downsample ratio
if [ $? -eq 0 ]
then
    baseline_mosdepth_jobid=`echo ${baseline_mosdepth_submission} | awk '{print $4}'`
    spikein_mosdepth_jobid=`echo ${spikein_mosdepth_submission} | awk '{print $4}'`
    calc_ratio_submission=$(sbatch --chdir ${output_dirpath} --mail-type FAIL \
                                   --error calculate-ratio.err --output calculate-ratio.out \
                                   --job-name calculate-ratio --tasks-per-node 1 --mem 2gb \
                                   --time 2:00:00 --partition compute --account proj-fs0002 \
                                   --dependency afterok:${baseline_mosdepth_jobid}:${spikein_mosdepth_jobid} \
                                   run_calculate_ratio.sh ${baseline_prefix} ${spikein_prefix} \
                                   ${output_dirpath} ${ratio} ${calcratio_script_path}
                         )
    echo `date` "- INFO: calculating ratios for subsampling"
    # run downsampling
    if [ $? -eq 0 ]
    then
        calcratio_jobid=`echo ${calc_ratio_submission} | awk '{print $4}'`
        baseline_subsample_submission=$(sbatch --chdir ${output_dirpath} --mail-type FAIL \
                                               --error baseline_subsample.err --output baseline_subsample.out \
                                               --job-name subsample --tasks-per-node 10 --mem 30gb \
                                               --time 3:00:00 --partition medium --account proj-fs0002 \
                                               --dependency afterok:${calcratio_jobid} \
                                               run_subsample.sh ${baseline_bampath} ${samtools_path} ${output_dirpath}
                                     )
        spikein_subsample_submission=$(sbatch --chdir ${output_dirpath} --mail-type FAIL \
                                              --error spikein_subsample.err --output spikein_subsample.out \
                                              --job-name subsample --tasks-per-node 10 --mem 30gb \
                                              --time 3:00:00 --partition medium --account proj-fs0002 \
                                              --dependency afterok:${calcratio_jobid} \
                                              run_subsample.sh ${spikein_bampath} ${samtools_path} ${output_dirpath}
                                    )
        echo `date` "- INFO: running subsampling"
        if [ $? -eq 0 ]
        then
            baseline_subsample_jobid=`echo ${baseline_subsample_submission} | awk '{print $4}'`
            spikein_subsample_jobid=`echo ${spikein_subsample_submission} | awk '{print $4}'`

            merge_submission=$(sbatch --chdir ${output_dirpath} --mail-type FAIL \
                                      --error merge.err --output merge.out \
                                      --job-name merge --tasks-per-node 10 --mem 16gb \
                                      --time 3:00:00 --partition medium --account proj-fs0002 \
                                      --dependency afterok:${baseline_subsample_jobid}:${spikein_subsample_jobid} \
                                      run_merge.sh ${baseline_prefix} ${spikein_prefix} \
                                      ${samtools_path} ${output_dirpath}
                            )
            echo `date` "- INFO: merge subsample data"
            if [ $? -eq 0 ]
            then
                merge_jobid=`echo ${merge_submission} | awk '{print $4}'`
                merge_bampath=${output_dirpath}/${baseline_prefix}"_"${spikein_prefix}"_merged.sorted.bam"
                merge_prefix=`basename ${merge_bampath} .bam`
                
                subsample_mosdepth_submission=$(sbatch --chdir ${output_dirpath} --mail-type FAIL \
                                                       --error merge-mosdepth.err --output merge-mosdepth.out \
                                                       --job-name mosdepth --tasks-per-node 8 --mem 16gb \
                                                       --time 3:00:00 --partition medium --account proj-fs0002 \
                                                       --dependency afterok:${merge_jobid} \
                                                       run_mosdepth.sh ${merge_bampath} \
                                                       ${merge_prefix} ${mosdepth_path}
                                             )
                echo `date` "- INFO: mosdepth on merged command - queued up"
            else
                echo `date` "- ERROR: mosdepth on merged command - not queued up"
            fi
        else
            echo `date` "- ERROR: merge command - not queued up"
        fi
    else
        echo `date` "- ERROR: subsample commands - not queued up"
    fi
fi
```
