## Supplemental File S3 for "MosaicSim: A Novel Mosaic Variant Simulator Reveals Diminishing Returns of Ultra-High Coverage for Mosaic Variant Detection": vaf_seperator.py

```
import pandas as pd
import gzip
import sys

def main(input_file:str,VAF:float, output_file:str):
    # Input vcf file
    vcf_file = rf'{input_file}'
    # check if vcf file exists 
    try:
        open(vcf_file, 'r')
    except FileNotFoundError:
        print(f"Error: {vcf_file} not found")
        sys.exit(1)

    # Determine if the vcf file is compressed
    compression=None
    if vcf_file.endswith('.gz'):
        compression='gzip'
    df = pd.read_csv(vcf_file, sep='\t', comment='#', header=None, compression=compression)


    df_af = df.iloc[:, 8:]
    df_af.columns = ['def','val']
    # get index from def column where DR and DV is
    df_af['DR_idx'] = df_af['def'].apply(lambda x: x.split(':').index('DR'))
    df_af['DV_idx'] = df_af['def'].apply(lambda x: x.split(':').index('DV'))
    # get the DR and DV value according to the index DR_idx from val column
    df_af['DR'] = df_af.apply(lambda x: int(x['val'].split(':')[x['DR_idx']]), axis=1)
    df_af['DV'] = df_af.apply(lambda x: int(x['val'].split(':')[x['DV_idx']]), axis=1)


    # get the AF value if DR and DV is not 0
    df_af['AF'] = df_af.apply(lambda x: x['DV']/x['DR'] if x['DR'] != 0 else 0, axis=1)


    # calculating the minor allele frequency (MAF)
    df_af['MAF'] = df_af.apply(lambda x: 1-x['AF'] if x['AF'] > 0.5 else x['AF'], axis=1)
    # get all indices where MAF is greater than 0.01 add true to the collumn qualified
    df_af['qualified'] = df_af['MAF'] >= VAF


    df['qualified'] = df_af['qualified']


    # load comment lines from vcf file
    comments = []
    vcf_file_handler = open(vcf_file, "r") if "gz" not in vcf_file \
        else gzip.open(vcf_file, "rt")
    for line in vcf_file_handler:
        if line.startswith("#"):
            comments.append(line)
        else:
            break
    vcf_file_handler.close()

    # convert dataframe to a list of string which are tab separated for each column where the qualified column is true
    df_qualified = df[df['qualified'] == True]
    # drop the qualified column
    df_qualified = df_qualified.drop(columns=['qualified'])


    # for each row in the dataframe convert the row to a string, where each column is tab seperated, and add a newline
    # character
    df_qualified_mod = df_qualified.apply(lambda x: '\t'.join(x.astype(str)) + '\n', axis=1)
    qualified = df_qualified_mod.tolist()


    output_file = rf'{output_file}'
    # write the qualified lines to a new vcf file
    with open(output_file, 'w') as f:
        f.writelines(comments)
        f.writelines(qualified)

if __name__ == '__main__':
    main(sys.argv[1], float(sys.argv[2]) ,sys.argv[3])
```
