## Supplemental File S3 for "MosaicSim: A Novel Mosaic Variant Simulator Reveals Diminishing Returns of Ultra-High Coverage for Mosaic Variant Detection": extract_read_bam_out.py

```
import pysam
import sys
from Bio import SeqIO
import argparse
import gzip

DEBUG = False

def write_fastx_record(fh, id, read, qual):
    if not qual:
        qual = [60] * len(read)
    qual_str = "".join([chr(v + 33) for v in qual])
    fh.write(f"@{id}\n")
    fh.write(f"{read}\n")
    fh.write("+\n")
    fh.write(f"{qual_str}\n")

def write_bam_record(bam_out, read_name, new_seq, new_qual, ref_name, ref_start, cigartuples):
    """Writes an edited read to a BAM file."""
    a = pysam.AlignedSegment()
    a.query_name = read_name
    a.query_sequence = new_seq
    a.flag = 0  # Adjust the flag as needed
    a.reference_id = bam_out.get_tid(ref_name)
    a.reference_start = ref_start
    a.mapping_quality = 255  # Placeholder quality
    a.cigar = cigartuples  # Keep the original or adjust if needed
    a.query_qualities = pysam.qualitystring_to_array("".join(chr(q + 33) for q in new_qual)) if new_qual else None
    bam_out.write(a)


def edit_read(ref_seq, ref_start, read, quals, cigartuples, variant_pos, variant_length, variant_op, insert_seq):
    if DEBUG:
        print(f"----ref start {ref_start} read len {len(read)}")
        print(f"----variant pos {variant_pos} var length {variant_length} var op {variant_op} var seq {insert_seq}")
        print(f"----cigar tuples {cigartuples}")
    ref_pos = ref_start # ref pos, variant pos are 1 indexed
    last_ref_pos = ref_pos
    read_var_pos = -1
    new_seq = ""
    new_qual = ""
    read_pos = 0 # read pos is 0 indexed
    last_read_pos = 0
    for (c, l) in cigartuples:
        if c in (0, 7, 8):
            if DEBUG:
                new_seq += read[read_pos:read_pos + l]
            read_pos += l
            ref_pos += l
        elif c == 1:
            if DEBUG:
                new_seq += read[read_pos: read_pos + l]
            read_pos += l
        elif c == 2:
            ref_pos += l
        elif c == 3:
            ref_pos += l
        elif c == 4:
            if DEBUG:
                new_seq += read[read_pos: read_pos + l]
            read_pos += l
        elif c == 5 or c == 6:
            pass
        else:
            raise RuntimeError("Unknown cigr ops found " + c)
        if ref_pos > variant_pos and read_var_pos == -1:
            if DEBUG:
                print(f"Current ref pos {ref_pos} last ref pos {last_ref_pos} variant pos {variant_pos}")
            if c == 2 or c == 3:
                # if there's an del variant here, then we remove it from the following seq
                # if there's an insertion or snv do nothing
                if variant_op == "DEL":
                    if variant_length > (ref_pos - variant_pos):
                        variant_length -= (ref_pos - variant_pos)
                        variant_pos = ref_pos
            else:
                read_var_pos = last_read_pos + (variant_pos - last_ref_pos) - 1
        last_ref_pos = ref_pos
        last_read_pos = read_pos
    if DEBUG and new_seq != read:
        print(f"Something went wrong! Constructed read {new_seq} doesn't match query {read}")
    if read_var_pos != -1:
        if variant_op == "INS":
            new_seq = read[:read_var_pos + 1] + insert_seq + read[read_var_pos + 1:]
            if quals:
                qual_seq = [60] * len(insert_seq)
                new_qual = list(quals[:read_var_pos + 1]) + qual_seq + list(quals[read_var_pos + 1:])
        elif variant_op == "SNV":
            new_seq = read[:read_var_pos] + insert_seq + read[read_var_pos+1:]
            if quals:
                new_qual = list(quals[:read_var_pos]) + [60] + list(quals[read_var_pos+1:])
        else:
            new_seq = read[:read_var_pos] + ( read[read_var_pos + variant_length:] if (read_var_pos + variant_length) < len(read) else "")
            if quals:
                new_qual = list(quals[:read_var_pos]) + ( list(quals[read_var_pos + variant_length:]) if (read_var_pos + variant_length) < len(read) else [])
        if DEBUG:
            print(f"Insert variant at {read_var_pos}")
            print(f"Bef - {read}")
            print(f"Aft - {new_seq}")
            print(f"Len diff {len(new_seq) - len(read)}")
        return (new_seq, new_qual)
    else:
        if DEBUG:
            print("No variant location found. Does the last cigar entry contain insertions to the ref or soft clips?")
    return (None, None)

def read_ref(fq_file):
    refs = {}
    for seq_record in SeqIO.parse(fq_file, "fasta"):
        refs[seq_record.id] = repr(seq_record.seq)
    return refs

def load_bam_queries(bam_file, refs, outfile):
    import random
    bam = pysam.AlignmentFile(bam_file, "r")
    with open(outfile, 'w') as fh:
        for i, row in enumerate(bam):
            if i > -1:
                if not row.query_sequence:
                    print("No query sequence for ", row.query_name)
                    continue
                if (row.reference_name and row.query_sequence):
                    variant_pos = row.reference_start + random.randint(0, len(row.query_sequence) - 1)
                    ops = ["INS", "DEL", "SNV"]
                    variant_op = ops[random.randint(0, len(ops) - 1)]
                    variant_length = 1 if variant_op == "SNV" else random.randint(1, 10000)
                    bases = ['A', 'T', 'C', 'G']
                    var_seq = "".join([bases[random.randint(0, 3)] for _ in range(variant_length)])
                    new_seq, new_qual = edit_read(refs[row.reference_name], row.reference_start, row.query_sequence, row.query_qualities, row.cigartuples, variant_pos, variant_length, variant_op, var_seq)
                    if new_seq:
                        write_bam_record(bam_out, f"{row.query_name}|after|{variant_op}|{variant_length}|{variant_pos}", new_seq, new_qual, row.reference_name, row.reference_start, row.cigartuples)

if __name__ == "__main__":
    parser = argparse.ArgumentParser(
                    prog='Read editor',
                    description='edit reeads based on a variant and a BAM file')
    parser.add_argument('--bam', help='bam file')
    parser.add_argument('--ref', help='ref file')
    parser.add_argument('--out', help='output fastx name')
    args = parser.parse_args()
    refs = read_ref(args.ref)
    bam1_queries = load_bam_queries(args.bam, refs, args.out)
```
