## Supplemental File S3 for "MosaicSim: A Novel Mosaic Variant Simulator Reveals Diminishing Returns of Ultra-High Coverage for Mosaic Variant Detection": README.md

# TweakVar: Simulation of Mosaic Variants in Empirical Sequencing Data

**Hackathon team: Lead: Fritz Sedlazeck - Developers: Erik Stricker, Xinchang Zheng, Michal Izydorczyk, Chi-Lam Poon, Philippe Sanio, Farhang Jaryani, Joyjit Daw, Divya Kalra, Adam Alexander - Writers: Erik Stricker, Sontosh Deb**

*We provide two simulation workflows which output sequencing read files with artificial mosaic variants and a ground truth mosaic variant annotation file for the validation of mosaic variant callers.*


## Table of Contents
|1. [Background](#background)<br>2. [Installation](#installation)<br>3. [Dependencies](#dependencies)<br>4. [Tests](#tests)<br>5. [How to Use It](#how-to-use-it)<br>6. [Example Implementation](#example-implementation)<br>7. [Method Description](#method-description)<br>8. [Contributers](#contributers)<br>9. [References](#references)<br><img width=1000/>|<img src="https://github.com/erikstricker/MosaicSim/blob/93ae22dd82122271b36fc1b585e283c59a3f4795/images/TweakVar_workflow_simple.png"  style="width: auto; display: block; margin: auto;">|
|:------|-:|


## Background

<img src="https://github.com/erikstricker/MosaicSim/blob/93ae22dd82122271b36fc1b585e283c59a3f4795/images/BackgroundMV.png"  style="width: auto; display: block; margin: auto;">

In the context of individual genome comparison, mutations that appear within a small fraction of the population are considered rare variants[<sup>1</sup>](#1). When assessing a population of cells from a tissue of the same individual in turn, rare variants only present in a small fraction of the cells are defined as mosaic variants (MVs)[<sup>2</sup>](#2). Recent studies have shown that there is potential disease associations of for certain MVs[<sup>2</sup>](#2). However, MVs are challenging to detect because they are mixed in with data from the non-mutated cells and present in the same sequencing file. Therefore, several pipelines have been developed or adjusted to extract mosaic single nucleotide, structural or indel variants from whole genome sequencing data such as Sniffles[<sup>3</sup>](#3), DeepMosaic[<sup>4</sup>](#4), Mutect2[<sup>5</sup>](#5), DeepVariant[<sup>6</sup>](#6). To benchmark and validate the efficiency and accuracy of these methods, sequencing files with known MVs are necessary. We developed a simulation workflow TweakVar (*Tweak* *Var*iants within existing reads of one individual), which outputs sequencing read files with artificial MVs and a ground truth annotation file for the MVs. TweakVar accomplishes this by creating a list of random mutations and modifying a fraction of existing reads to match the user-defined MV frequency.


## Installation
These instructions are valid for **Linux x86**. For other platforms (e.g. MacOS), instructions will need to be adapted.

Load conda, e.g.
```
module load miniconda/3
```
or
```
module load anaconda3 ##sometimes python has to be loaded before starting an environment)
module load python
```

Installing two conda environments with python 3.10
```
conda create -n MosaicSim python=3.10
conda init
conda activate MosaicSim
```

Unload python if previously loaded so that python 3.10 from conda environment will be used
```
module unload python
```

Obtain the MosaicSim from github (replace $HOME with your preferred installation director)
```
cd  $HOME
git clone https://github.com/erikstricker/MosaicSim.git
```

To install the relevant python dependencies, run
```
REPO_ROOT="$HOME/MosaicSim/"

pip install -r $REPO_ROOT/requirements.txt
```
Ensure to also load mosdepth>0.3.2 (for SpikeVar), samtools >1.15.1, and bcftools>1.19

Once the requirements are installed, please install or load the following additional packages

_Installation_
```
conda install -c bioconda samtools bcftools mosdepth
```
_Loading (e.g.)_
```
export PATH=/path/to/software/mosdepth/mosdepth-0.3.2/bin:$PATH
export PATH=/path/to/software/samtools/samtools-1.21/bin:$PATH
export PATH=/path/to/software/bcftools/bcftools-1.19/bin:$PATH
```
or
```
module load mosdepth-0.3.2
module load samtools-1.21
module load bcftools-1.19
```

## Dependencies
  
### TweakVar
- pysam (0.21.0) 
- numpy (1.25.2)
- biopython (1.81)
- samtools 1.15.1
- bcftools 1.19

## How to Use It

### TweakVar

Once the Python dependencies are installed, the scripts can be run directly from the `scripts` subfolder.

#### 0) TweakVar Prep

Before getting started ensure that:
* The input file is in bam format with corresponding index (.bai) file
* The corresponding fasta reference file is downloaded with appropriate index (.fai) file

#### 1) TweakVarSimulator - Generate Simulated VCF

<img src="https://github.com/erikstricker/MosaicSim/blob/93ae22dd82122271b36fc1b585e283c59a3f4795/images/TweakVarSimulator.png" height="130" align="right">

The **TweakVarSimulator** generates a random set of mosaic variants, including **single nucleotide variants (SNVs)** and **structural variants (SVs)**. These variants can be customized by adjusting parameters such as **variant allele frequency (VAF)**, **the number of variants to simulate**, and **variant size**.

The output is a **VCF file in Sniffles format**, which serves as:  
- **Input** for the read editor step (described below) to modify sequencing reads by inserting the simulated variants.  
- **Ground truth** for evaluating mosaic variant callers.  

##### Usage
```bash
python tweakvarsimulator.py -i <path_to_bam> -T <path_to_ref> -o <output_path_prefix> [optional arguments]
```

##### Required Parameters  
| Parameter | Description |
|-----------|-------------|
| `-i, --input` | Path to the **input BAM file** containing sequencing reads. |
| `-T, --reference` | Path to the **reference genome FASTA file**. |
| `-o, --output` | Prefix for the **output files** (e.g., VCF files for simulated SNVs and SVs). |

##### Optional Parameters  
| Parameter | Description | Default Value |
|-----------|-------------|---------------|
| `-s, --seed` | **Random seed** for reproducibility. If not set, the results will vary between runs. The same seed with different input but identical parameters will lead to the same list of loci. | `0` |
| `-minAFsv, --minimum_allele_frequency_sv` | **Minimum allele frequency** for simulated **structural variants (SVs)**. | `0.01` |
| `-maxAFsv, --maximum_allele_frequency_sv` | **Maximum allele frequency** for **SVs**. | `0.05` |
| `-minAFsnv, --minimum_allele_frequency_snv` | **Minimum allele frequency** for **single nucleotide variants (SNVs)**. | `0.01` |
| `-maxAFsnv, --maximum_allele_frequency_snv` | **Maximum allele frequency** for **SNVs**. | `0.05` |
| `-numsv, --number_of_svs` | Number of **structural variants (SVs)** to simulate. | `50` |
| `-numsnv, --number_of_snvs` | Number of **single nucleotide variants (SNVs)** to simulate. | `200` |
| `-minsvl, --minimum_sv_length` | **Minimum length** of simulated **SVs** (in base pairs). | `50` |
| `-maxsvl, --maximum_sv_length` | **Maximum length** of simulated **SVs** (in base pairs). | `10,000` |
| `-sub, --substitution_rate` | **Probability of generating a SNP** versus an indel (in SNVs). A value of **1.0** means only SNPs will be generated. | `1.0` |
| `-insdelsnv, --insdel_snv_rate` | **Probability of generating an insertion vs. a deletion** in SNVs. A value of **0.5** means equal chances of insertion and deletion. | `0.5` |
| `-insdel, --insdel_sv_rate` | **Probability of generating an insertion vs. a deletion** in SVs. A value of **0.7** means insertions are more likely than deletions. | `0.7` |
| `--ref_chroms` | List of **chromosomes to include** in the simulation (e.g., `chr1 chr2 ... chrY`). Only these chromosomes will be used for variant generation. | *All chromosomes in the BAM header* |
| `-snv, --SNV_truth_file` | SNV truth file of which variant locations, allele frequencies, and alternate alleles are used as the foundation to create a new SNV VCF file with updated coverage and read count information. | Will be ignored if not present. |
| `-sv, --SV_truth_file` | SV truth file of which variant locations, allele frequencies, and alternate alleles are used as the foundation to create a new SV VCF file with updated coverage and read count information. | Will be ignored if not present. |
| `-bps, --bp_shift` | Allows the shift of all bp positions up- or downstream by the indicated number of bps. | `0` |
| `-imc, --ignore_minimum_cov` | Ignores minimum coverage requirement for the selection of variant loci. This is especially useful when selecting identical regions in down-sampled files. | `FALSE` |

#### 2) TweakVarEditor - Generate Edited Reads Based on Simulated VCF

<img src="https://github.com/erikstricker/MosaicSim/blob/93ae22dd82122271b36fc1b585e283c59a3f4795/images/TweakVarEditor.png"  height="200" align="right">

This command above takes in the VCF which determines which variants to introduce into the reads.
The BAM file is used to find the reads which overlap with variant locations. Only a subset of the reads
corresponding to a particular variant location are edited. This is determined by the allele frequency.
The output BAM file retains the alignment info and now contains the edited reads. Also, the query name of each read is kept the same.

##### Usage
```bash
python tweakvareditor.py -v <simulated_vcf> -b <path_to_bam> -T <path_to_ref> -o <output_bam>
```
##### Required Parameters  
| Parameter               | Description |
|-------------------------|-------------|
| `-v, --vcf_file`         | VCF file from TweakVarSimulator  (SNV or SV). |
| `-b, --bam_file`         | Input BAM file. |
| `-T, --ref_file`         | Reference genome file for BAM. |
| `-o, --out_file`         | Output BAM file name and directory. |

##### Optional Parameters  
| Parameter | Description | Default Value |
|-----------|-------------|---------------|
| `-of, --output_format` | **Output format** can be either sorted `bam` file or unmapped `fastq` file. Fastq format has to converted into a mapped bam file with an aligner, while mapped reads in the bam file will directly replace existing reads at their original position. | `"bam"` |

#### 3) TweakVarMerger - Re-Align Modified Reads and Merge Them

Replace the reads in the original dataset with the modified reads. This step is time intensive so make sure to run it in a cluster environment for large files.

##### Usage
```bash
python tweakvarmerger.py -b <input bam> -m <bam file with modified reads> -o <out_dir and file name>
```

##### Required Parameters  

| Parameter        | Description                                   |
|-----------------|-----------------------------------------------|
| `-b`, `--input_bam`  | Unmodified input BAM file                 |
| `-m`, `--modified_bam` | BAM file with modified reads only        |
| `-o`, `--out_file`  | Output file name (default: `tweakvar_modified.bam` in the current directory) |


<img src="https://github.com/erikstricker/MosaicSim/blob/93ae22dd82122271b36fc1b585e283c59a3f4795/images/TweakVarMerger.png"  height="200" align="right">

#### 4) Run Your Favorite Mosaic Variant Caller and Compare Results
For SV calling, we tried Sniffles:
```
sniffles --input <MOD_MERGED_BAM> --vcf sniffles_out.vcf --mosaic --threads 8
```
* MOD_MERGED_BAM is the output bam file from `tweakvarmerger.py`

for SNV calling, we ran longshot:
```
longshot --bam <MOD_MERGED_BAM> --ref <REF> --out longshot_out.vcf --min_cov 3
```


## Example Implementation

### TweakVar

Here, we use the TweakVar workflow to modify reads of HG002 directly at their reference position by including artificial mutations. To demonstrate the wide application of this tool, we generate a random distribution of allele frequencies between 1% and 40%. In contrast to the above approach, we do not introduce new haplotypes with this. However, more complex mutations (e.g. rearrangements, duplication, or very long structural variants) will not be able to be introduced to the data itself, since the size of the reads is limited.


#### 0) Fetch Data
In order to simulate and edit reads, the pipeline first needs an initial set of aligned reads and a reference. For our demonstration, we will use the GIAB datasets.

Reads (bam file)
```
mkdir $HOME/data ##or your data folder
wget -P $HOME/data https://ftp-trace.ncbi.nlm.nih.gov/ReferenceSamples/giab/data/AshkenazimTrio/HG002_NA24385_son/NIST_Illumina_2x250bps/novoalign_bams/HG002.GRCh38.2x250.bam
wget -P $HOME/data https://ftp-trace.ncbi.nlm.nih.gov/ReferenceSamples/giab/data/AshkenazimTrio/HG002_NA24385_son/NIST_Illumina_2x250bps/novoalign_bams/HG002.GRCh38.2x250.bam.bai
```

Reference
```
mkdir $HOME/data/ref 
wget -P $HOME/data/refhttps://ftp-trace.ncbi.nlm.nih.gov/ReferenceSamples/giab/release/references/GRCh38/GRCh38_masked_v2_decoy_gene.fasta.gz 
```


Then, we decompress the FASTA file.
```
gunzip $HOME/data/ref/GRCh38_masked_v2_decoy_gene.fasta.gz 
samtools faidx $HOME/data/ref/GRCh38_masked_v2_decoy_gene.fasta
```

We run the demonstration on chr22 only, so the dataset is filtered using
```
samtools view -b $HOME/data/HG002.GRCh38.2x250.bam chr22 \
  | samtools sort -o $HOME/data/chr22.HG002.GRCh38.2x250.bam
samtools index $HOME/data/chr22.HG002.GRCh38.2x250.bam
```


#### 1) TweakVarSimulator - Generate Variants and Modified Reads

Then we simulate variants
```
## Example Usage
```bash
module load anaconda3/2024.02
module load python
conda activate MosaicSim
module unload python
cd $HOME/MosaicSim
python scripts/tweakvarsimulator.py \
 -i $HOME/data/chr22.HG002.GRCh38.2x250.bam \
 -T $HOME/data/ref/GRCh38_masked_v2_decoy_gene.fasta \
 -o $HOME/test_output_dir/chr22/chr22.HG002.GRCh38.2x250_MAF0.01-0.05 \
 -s 0 -numsv 5 -numsnv 100
```

This command:  
- Uses `chr22.HG002.GRCh38.2x250.bam` as input and `GRCh38_masked_v2_decoy_gene.fasta` as the reference genome.  
- Outputs simulated VCFs to `test_output_dir/chr22`.  
- Sets a **random seed of 0** for reproducibility.  

#### 3) TweakVarEditor - Add Modified Reads Back In

Generate a set of modified reads with inserted variants.
```
module load anaconda3/2024.02
module load python
conda activate MosaicSim
module unload python
cd $HOME/MosaicSim
python  scripts/tweakvareditor.py \
 -v $HOME/test_output_dir/chr22/chr22.HG002.GRCh38.2x250_MAF0.01-0.05_SNV.vcf \
 -b $HOME/data/chr22.HG002.GRCh38.2x250.bam \
 -T $HOME/data/ref/GRCh38_masked_v2_decoy_gene.fasta  \
 -o $HOME/test_output_dir/chr22/output_SNV_chr22.HG002.GRCh38.2x250_MAF0.01-0.05_SNV.bam \
 -of "bam"
```

The modified reads can also be generated as an unmapped FASTQ file to simulate potential mapping error that could arise:
```
module load anaconda3/2024.02
module load python
conda activate MosaicSim
module unload python
cd $HOME/MosaicSim
python  scripts/tweakvareditor.py \
 -v $HOME/test_output_dir/chr22/chr22.HG002.GRCh38.2x250_MAF0.01-0.05_SNV.vcf \
 -b $HOME/data/chr22.HG002.GRCh38.2x250.bam \
 -T $HOME/data/ref/GRCh38_masked_v2_decoy_gene.fasta  \
 -o $HOME/test_output_dir/chr22/output_SNV_chr22.HG002.GRCh38.2x250_MAF0.01-0.05_SNV.fastq \
 -of "fastq"
```

#### 4) TweakVarMerger - Re-Align Modified Reads and Merge Them

Once the new reads are generated in BAM format, we then remove the old alignments with the same IDs of modified reads in the original BAM file, then insert the new alignments of modified reads back into this BAM file.
```
module load anaconda3/2024.02
module load python
conda activate MosaicSim
module unload python
cd $HOME/MosaicSim
python  scripts/tweakvarmerger.py \
 -b $HOME/data/chr22.HG002.GRCh38.2x250.bam \
 -m $HOME/test_output_dir/chr22/output_SNV_chr22.HG002.GRCh38.2x250_MAF0.01-0.05_SNV.bam \
 -o $HOME/test_output_dir/chr22/merged.modified_chr22.HG002.GRCh38.2x250.bam
```

If the reads were generated in FASTQ format, they need to be re-aligned using a tool like minimap2 and then re-inserted back into the dataset by replacing the original reads.

We can use `minimap2` for both short-read and long-read alignment. In the example, we tested on chromosome 22.
```
input_fastq="$HOME/test_output_dir/chr22/output_SNV_chr22.HG002.GRCh38.2x250_MAF0.01-0.05_SNV.fastq"
output_bam="$HOME/test_output_dir/chr22/output_SNV_chr22.HG002.GRCh38.2x250_MAF0.01-0.05_SNV.fastq.bam"
reference="$HOME/data/ref/GRCh38_masked_v2_decoy_gene.fasta"
```
Choose the alignment setting that best fits your needs: 
***short read - standard***
```
minimap2 -ax sr ${reference}  -t 14 --secondary=no $input_fastq | samtools view -bS --no-PG - | \
samtools sort -o ${output_bam}
samtools index -b ${output_bam}
```
***long read - standard***
```
minimap2 -ax map-ont ${reference}  -t 14 --secondary=no $input_fastq | samtools view -bS --no-PG - | \
samtools sort -o ${output_bam}
samtools index -b ${output_bam}
```
***short read - error permissable***
```
minimap2 -ax sr \
    -A 2 -B 3 -O 5,56 -E 4,1 --score-N 0 -e 0 \
    --max-chain-skip 25 \
    ${reference} \
    -t 14 --secondary=no \
    $input_fastq | \
samtools view -bS --no-PG - | \
samtools sort -o ${output_bam}
samtools index -b ${output_bam}
```
***long read - error permissable***
```
minimap2 -ax map-ont \
    -k 19 -w 5 \
    -A 2 -B 4 -O 2,20 -E 2,1 \
    --score-N 0 --max-chain-skip 50 --max-chain-gap 10000 \
    -e 0 \
    $HOME/data/ref/hs37d5.fa \
    -t 14 $input_fastq | \
samtools view -bS --no-PG - | \
samtools sort -o ${output_bam}
samtools index -b ${output_bam}
```

Now that we have the alignments for modified reads, we then remove the old alignments with the same IDs of modified reads in the original bam, then insert the new alignments of modified reads back into this bam.
```
module load anaconda3/2024.02
module load python
conda activate MosaicSim
module unload python
cd $HOME/MosaicSim
python  scripts/tweakvarmerger.py \
 -b $HOME/data/chr22.HG002.GRCh38.2x250.bam \
 -m $HOME/test_output_dir/chr22/output_SNV_chr22.HG002.GRCh38.2x250_MAF0.01-0.05_SNV.fastq.bam \
 -o $HOME/test_output_dir/chr22/merged.modified_chr22.HG002.GRCh38.2x250.bam
```

#### 5) Run Your Favorite Mosaic Variant Caller

Run you choice of mosaic variant caller on the modified `merged.modified_chr22.HG002_hs37d5_ONT-UL_GIAB_20200122.phased.bam` file and compare the results with the simulated `chr22_HG002_hs37d5_ONT-UL_GIAB_20200122.phased_MAF0.01-0.05_SNV.vcf` file.

#### 6) Results

The `chr22.HG002.GRCh38.2x250.bam` and `merged.modified_chr22.HG002.GRCh38.2x250.bam` were both visualized on IGV to get a subjective view of whether the modified reads led to mosaic variants being introduced and detected.
Below are 2 of several variants that overlapped between the ground truth and caller VCF and were confirmed in the underlying data.

<img src="https://github.com/erikstricker/MosaicSim/blob/93ae22dd82122271b36fc1b585e283c59a3f4795/images/TweakVarMosaicSNV1.png" align="center"/>
<p align="center"><b>A mosaic SNV at position chr22:10961073</b></p>
<br />

<img src="https://github.com/erikstricker/MosaicSim/blob/93ae22dd82122271b36fc1b585e283c59a3f4795/images/TweakVarMosaicSNV2.png" align="center"/>
<p align="center"><b>A mosaic SNV at position chr22:30088854</b></p>


## Method Description 

### TweakVar - Creation of Sequencing Data With a Subset of Modified Reads
<img src="https://github.com/erikstricker/MosaicSim/blob/93ae22dd82122271b36fc1b585e283c59a3f4795/images/TweakVar_flowchart_updated.png" width="500"/>
<p align="justify">
<b>TweakVar workflow, with major steps to assess the sensitivity and accuracy of the mosaic variant callers. (A, B: individual samples, A/B: merged samples, .bam and .vcf: input and output file formats in different steps, Black header boxes: tool or file names, Green header boxes: simulated files or final files used for validation comparisons)</b>
</p>  
<br />

The TweakVar workflow produces a modified aligned sequence file in .bam format. This file contains modified reads simulating randomly positioned mosaic variants with user-defined VAF in random locations and is accompanied by a .vcf file containing the locations of the simulated mosaic variants with user-defined VAF. 

The TweakVar workflow can be broadly split into 3 parts: 
1) The TweakVarSimulator takes an aligned BAM, a reference, and several parameters such as range of VAF, variant sizes, etc. to generate a set of simulated mosaic SV and SNVs. It does so by choosing a random location and VAF from the given range and then evaluating whether that location has sufficient coverage for the desired VAF. If that condition is met, that variant is added to the output VCF. 
2) The TweakVarEditor is responsible for inserting the simulated variants into the query sequences from the original dataset to generate modified reads with the mosaic variants built-in. The TweakVarEditor accepts a BAM, reference and the simulated VCF file as an input. Then for each variant, it fetches the overlapping reads from the BAM file, subsamples the reads to get the coverage that satisfies the desired VAF, and traverses the cigar string, query and reference sequences for each alignment to find the exact location to insert the variant. Once a modified read is created, it is written out into a FASTQ file. Note that for all new bases (SNVs or inserts), a q-score of 60 is chosen. The parsing and traversing of VCF, BAM and reference files are performed using APIs from pysam, biopython.SeqIO and numpy.
3) The TweakVarMerger re-introduces the modified reads into the original dataset. It does so by first removing the modified read ids from the input BAM to create a filtered BAM. Then the modified reads are aligned against the reference, and merged with the filtered BAM. The end result is a BAM with the same set of read ids as the original dataset, except with some reads modified to contain the mosaic variants. 
The output of this pipeline is thus a modified BAM and a VCF file which provides the truth set for the mosaic variants.

## Contributers


|<img src="https://github.com/erikstricker/MosaicSim/blob/93ae22dd82122271b36fc1b585e283c59a3f4795/images/Erik Stricker.jpg" width="150"/><br>Erik Stricker|<img src="https://github.com/erikstricker/MosaicSim/blob/93ae22dd82122271b36fc1b585e283c59a3f4795/images/Chi-Lam Poon.jpg" width="150"/><br>Chi-Lam Poon|<img src="https://github.com/erikstricker/MosaicSim/blob/93ae22dd82122271b36fc1b585e283c59a3f4795/images/Philippe Sanio.jpg" width="150"/><br>Philippe Sanio|<img src="https://github.com/erikstricker/MosaicSim/blob/93ae22dd82122271b36fc1b585e283c59a3f4795/images/Xinchang Zheng.jpg" width="150"/><br>Xinchang Zheng|<img src="https://github.com/erikstricker/MosaicSim/blob/93ae22dd82122271b36fc1b585e283c59a3f4795/images/Farhang Jaryani.png" width="150"/><br>Farhang Jaryani|
|:-:|:-:|:-:|:-:|:-:|

|<img src="https://github.com/erikstricker/MosaicSim/blob/93ae22dd82122271b36fc1b585e283c59a3f4795/images/Joyjit Daw.png" width="150"/><br>Joyjit Daw |<img src="https://github.com/erikstricker/MosaicSim/blob/93ae22dd82122271b36fc1b585e283c59a3f4795/images/Michal Bogumil Izydorczyk.png" width="150"/><br>Michal Izydorczyk|<img src="https://github.com/erikstricker/MosaicSim/blob/93ae22dd82122271b36fc1b585e283c59a3f4795/images/Sontosh K Deb.jpg" width="150"/><br>Sontosh Deb|<img src="https://github.com/erikstricker/MosaicSim/blob/93ae22dd82122271b36fc1b585e283c59a3f4795/images/Fritz Sedlazeck.jpg" width="150"/><br>Fritz Sedlazeck |<img src="https://github.com/erikstricker/MosaicSim/blob/93ae22dd82122271b36fc1b585e283c59a3f4795/images/Alexander Adam.jpg" width="150"/><br>Adam Alexander|
|:-:|:-:|:-:|:-:|:-:|

|<img src="https://github.com/erikstricker/MosaicSim/blob/93ae22dd82122271b36fc1b585e283c59a3f4795/images/Divya Kalrai_placeholder.jpg" width="150"/><br>Divya Kalra|
|:-:|


## References

<a id="1">[1]</a> Sariya S, Lee JH, Mayeux R, Vardarajan BN, Reyes-Dumeyer D, Manly JJ, Brickman AM, Lantigua R, Medrano M, Jimenez-Velazquez IZ, Tosto G. Rare Variants Imputation in Admixed Populations: Comparison Across Reference Panels and Bioinformatics Tools. Front Genet. 2019;10:239. Epub 20190403. doi: 10.3389/fgene.2019.00239. PubMed PMID: 31001313; PMCID: PMC6456789.  

<a id="2">[2]</a> Miller CR, Lee K, Pfau RB, Reshmi SC, Corsmeier DJ, Hashimoto S, Dave-Wala A, Jayaraman V, Koboldt D, Matthews T, Mouhlas D, Stein M, McKinney A, Grossman T, Kelly BJ, White P, Magrini V, Wilson RK, Mardis ER, Cottrell CE. Disease-associated mosaic variation in clinical exome sequencing: a two-year pediatric tertiary care experience. Cold Spring Harb Mol Case Stud. 2020;6(3). Epub 20200612. doi: 10.1101/mcs.a005231. PubMed PMID: 32371413; PMCID: PMC7304353.  

<a id="3">[3]</a> Sedlazeck FJ, Rescheneder P, Smolka M, Fang H, Nattestad M, von Haeseler A, Schatz MC. Accurate detection of complex structural variations using single-molecule sequencing. Nat Methods. 2018;15(6):461-8. Epub 20180430. doi: 10.1038/s41592-018-0001-7. PubMed PMID: 29713083; PMCID: PMC5990442.  

<a id="4">[4]</a> Yang X, Xu X, Breuss MW, Antaki D, Ball LL, Chung C, Shen J, Li C, George RD, Wang Y, Bae T, Cheng Y, Abyzov A, Wei L, Alexandrov LB, Sebat JL, Network NBSM, Gleeson JG. Control-independent mosaic single nucleotide variant detection with DeepMosaic. Nat Biotechnol. 2023;41(6):870-7. Epub 20230102. doi: 10.1038/s41587-022-01559-w. PubMed PMID: 36593400; PMCID: PMC10314968.  

<a id="5">[5]</a> McKenna A, Hanna M, Banks E, Sivachenko A, Cibulskis K, Kernytsky A, Garimella K, Altshuler D, Gabriel S, Daly M, DePristo MA. The Genome Analysis Toolkit: a MapReduce framework for analyzing next-generation DNA sequencing data. Genome Res. 2010;20(9):1297-303. Epub 20100719. doi: 10.1101/gr.107524.110. PubMed PMID: 20644199; PMCID: PMC2928508.  

<a id="6">[6]</a> Poplin R, Chang PC, Alexander D, Schwartz S, Colthurst T, Ku A, Newburger D, Dijamco J, Nguyen N, Afshar PT, Gross SS, Dorfman L, McLean CY, DePristo MA. A universal SNP and small-indel variant caller using deep neural networks. Nat Biotechnol. 2018;36(10):983-7. Epub 20180924. doi: 10.1038/nbt.4235. PubMed PMID: 30247488.
