## Supplemental File S3 for "MosaicSim: A Novel Mosaic Variant Simulator Reveals Diminishing Returns of Ultra-High Coverage for Mosaic Variant Detection": tweakvareditor.py

```
#!/usr/bin/env python
# -*- coding: utf-8 -*-
import pysam
import argparse
from vcf_line_parser import VCFLineSV
import math
import random
import os

from extract_read_bam_out import edit_read, read_ref, write_fastx_record, write_bam_record

def argument_parser():
    parser = argparse.ArgumentParser(
        description="TweakVarEditor",
        prog="TweakVarEditor",
        formatter_class=argparse.RawTextHelpFormatter,
    )
    parser.add_argument(
        "-v",
        "--vcf_file",
        dest="vcf_file",
        type=str,
        help="vcf file from TweakVarSimulator (SNV or SV)",
        required=True,
    )
    parser.add_argument(
        "-b",
        "--bam_file",
        dest="bam_file",
        type=str,
        help="bam file",
    )
    parser.add_argument(
        "-T",
        "--ref_file",
        type=str,
        help="Reference genome file for BAM",
    )
    parser.add_argument(
        "-o",
        "--out_file",
        type=str,
        help="Output file name and dir",
    )
    parser.add_argument(
        "-of",
        "--output_format",
        type=str,
        help="Output format can be either bam or fastq",
    )

    return parser.parse_args()

def vcf_number_variants_bam_out(input_vcf_file, input_bam_file, refs, outfile):
    processed_read_ids = set()
    """Return the number of each variant separately and write output in BAM format."""
    bamfile = pysam.AlignmentFile(input_bam_file, 'rb')
    vcffile = pysam.VariantFile(input_vcf_file, 'r')  # SV or SNV VCF
    bam_out = pysam.AlignmentFile(outfile.replace(".bam", "_pre.bam"), 'wb', header=bamfile.header)  # Output BAM file

    for v in vcffile:
        var_type = v.info.get('SVTYPE') if 'SVTYPE' in v.info else 'SNV'
        chromosome = v.chrom
        start_position = v.start
        variant_af = v.info.get('AF')

        if var_type == 'DEL':
            end_position = start_position + abs(v.info.get('SVLEN'))
            variant_len = abs(v.info.get('SVLEN'))
            variant_seq = ""
        elif var_type == 'INS':
            variant_seq = v.alts[0]
            variant_len = len(variant_seq)
            end_position = start_position + variant_len
        elif var_type == 'SNV':
            variant_seq = v.alts[0]
            variant_len = 1
            end_position = start_position + variant_len

        print(f"{chromosome}:{start_position}-{end_position} LEN={variant_len} {variant_seq} AF={variant_af}")

        variant_reads = []
        for read in bamfile.fetch(chromosome, start_position, end_position):
            if read.query_sequence is not None and read.flag in {0, 16, 99, 147, 83, 163} and read.query_name not in processed_read_ids:
                variant_reads.append(read)

        no_reads = max(1, math.ceil(float(len(variant_reads)) * float(v.info.get('AF')[0])))
        print(no_reads, len(variant_reads))

        if no_reads > len(variant_reads):
            print(f"Coverage {len(variant_reads)} is below required minimum reads for variant {chromosome}:{start_position} AF={variant_af}")
            continue

        random.shuffle(variant_reads)
        ref_seq = refs[read.reference_name]
        num_reads_edited = 0
        mos_counter = 0
        norm_counter = 0

        for read in variant_reads:
            read_edited = False
            if num_reads_edited < no_reads:
                mos_counter += 1
                num_reads_edited += 1
                if var_type == 'SNV':
                    new_seq, new_qual = edit_read(ref_seq, read.reference_start, read.query_sequence, read.query_qualities, 
                                                  read.cigartuples, end_position, variant_len, var_type, variant_seq)
                else:
                    new_seq, new_qual = edit_read(ref_seq, read.reference_start, read.query_sequence, read.query_qualities, 
                                                  read.cigartuples, start_position, variant_len, var_type, variant_seq)               

                if new_seq:
                    write_bam_record(bam_out, read.query_name, new_seq, new_qual, read.reference_name, read.reference_start, read.cigartuples)
                    read_edited = True
                    processed_read_ids.add(read.query_name)

            if not read_edited:
                norm_counter += 1
                write_bam_record(bam_out, read.query_name, read.query_sequence, read.query_qualities, read.reference_name, read.reference_start, read.cigartuples)

        print("mos+&norm", mos_counter, norm_counter)
        if num_reads_edited != no_reads:
            print(f"Could not fulfill required minimum edited reads for variant at {chromosome}:{start_position}:{end_position} type {var_type}")

    bamfile.close()
    vcffile.close()
    bam_out.close()  # Close output BAM file

    # **Sort and Index the BAM file**
    pysam.sort("-o", outfile, outfile.replace(".bam", "_pre.bam"))
    pysam.index(outfile)
	
    os.remove(outfile.replace(".bam", "_pre.bam"))

    print(f"Sorted BAM file saved as: {outfile}")
    print(f"Index file created: {outfile}.bai")

    return

def vcf_number_variants_fastq_out(input_vcf_file, input_bam_file, refs, outfile):
    processed_read_ids = set()
    """return the number of each variant seprately"""
    bamfile = pysam.AlignmentFile(input_bam_file, 'rb')
    vcffile = pysam.VariantFile(input_vcf_file, 'r') # SV or SNV vcf
    
    with open(outfile, 'w') as out_fh:
        for v in vcffile:
            var_type = v.info.get('SVTYPE') if 'SVTYPE' in v.info else 'SNV'
            ## does pysam return errors? maybe try & except ##
            # if obj.ERROR:
            #     continue
            chromosome = v.chrom
            start_position = v.start
            variant_af = v.info.get('AF')
            if var_type =='DEL':
                end_position = start_position + abs(v.info.get('SVLEN'))
                variant_len=abs(v.info.get('SVLEN'))
                variant_seq=""
            elif var_type =='INS':
                variant_seq = v.alts[0]
                variant_len = len(variant_seq)
                end_position = start_position + variant_len
            elif var_type =='SNV':
                variant_seq = v.alts[0]
                variant_len = 1
                end_position = start_position + variant_len
            #svtype = "INS"
            #variant_len = 100
            #variant_seq = 'T' * variant_len
            print(f"{chromosome}:{start_position}-{end_position} LEN={variant_len} {variant_seq} AF={variant_af}")
            variant_reads = []
            if var_type =='SNV':
                for read in bamfile.fetch(chromosome, start_position, end_position):
                    ##Test
                    ##reads = bamfile.fetch(chromosome, start_position, end_position); read = next(reads); print(read)
                    if read.query_sequence is not None and read.flag in {0, 16, 99, 147, 83, 163} and read.query_name not in processed_read_ids:
                        variant_reads.append(read)
            else:
                for read in bamfile.fetch(chromosome, start_position, end_position):
                    ##Test
                    ##reads = bamfile.fetch(chromosome, start_position, end_position); read = next(reads); print(read)
                    if read.query_sequence is not None and read.flag == 0 and read.query_name not in processed_read_ids:
                        variant_reads.append(read)

            no_reads = max(1, math.ceil(float(len(variant_reads)) * float(v.info.get('AF')[0])))
            print(no_reads, len(variant_reads))
            if no_reads > len(variant_reads):
                print(f"Coverage {len(variant_reads)} is below required minimum reads for variant {chromosome}:{start_position} AF={variant_af}")
                continue
            random.shuffle(variant_reads)
            #print(f"DR={obj.DR},AF={obj.AF},SVTYPE={obj.SVTYPE},CHROMOE={obj.CHROM},POS={obj.POS},SV_LENGTH={obj.SVLEN}")
            #print(f"{obj.DR}AF={obj.AF},SVTYPE={obj.SVTYPE},CHROMOE={obj.CHROM},POS={obj.POS},SV_LENGTH={obj.SVLEN},SV={obj.ALT}")
            ref_seq = refs[read.reference_name]
            num_reads_edited = 0
            mos_counter=0
            norm_counter=0
            for read in variant_reads:
                read_edited = False
                if num_reads_edited < no_reads:
                    mos_counter+=1
                    num_reads_edited += 1
                    if var_type =='SNV':
                        new_seq, new_qual = edit_read(ref_seq, read.reference_start, read.query_sequence, read.query_qualities, 
                                                  read.cigartuples, end_position, variant_len, var_type, variant_seq)
                    else:
                        new_seq, new_qual = edit_read(ref_seq, read.reference_start, read.query_sequence, read.query_qualities, 
                                                  read.cigartuples, start_position, variant_len, var_type, variant_seq)               
                    if new_seq:
                        query_name = read.query_name
                        write_fastx_record(out_fh, query_name, new_seq, new_qual)
                        
                        read_edited = True
                        processed_read_ids.add(read.query_name)
                if not read_edited:
                    norm_counter+=1
                    write_fastx_record(out_fh, read.query_name, read.query_sequence, read.query_qualities)
            print("mos+&norm",mos_counter,norm_counter)
            if num_reads_edited != no_reads:
                print("Could not fulfill required minimum edited reads for variant at {chromosome}:{start_position}:{end_position} type {svtype}")

        bamfile.close()
        vcffile.close()
        return

        print(f"FastQ file saved as: {out_fh}")


def main():
    args=argument_parser()
    refs = read_ref(args.ref_file)
    output_format = args.output_format
    if args.output_format is None:
        output_format = "bam"
    else:
        output_format = args.output_format

    if output_format == "bam":
        vcf_number_variants_bam_out(args.vcf_file, args.bam_file, refs, args.out_file)
    else:
        vcf_number_variants_fastq_out(args.vcf_file, args.bam_file, refs, args.out_file)
    out_file = args.out_file
if __name__ == "__main__":
	main()
```
