## Supplemental File S3 for "MosaicSim: A Novel Mosaic Variant Simulator Reveals Diminishing Returns of Ultra-High Coverage for Mosaic Variant Detection": tweakvarmerger.py

```
#!/usr/bin/env python

import pysam
import os
from argparse import ArgumentParser
from argparse import ArgumentDefaultsHelpFormatter


def filter_and_merge_bam(input_bam, modified_bam, out_dir, out_file):
    print('start filtering...')
    modified_reads = fetch_modified_reads(modified_bam, out_dir)

    bam = pysam.AlignmentFile(input_bam, 'rb')
    filtered_bam = pysam.AlignmentFile(f'{out_dir}/filtered.bam', 'wb', template=bam)
    for aln in bam:
        if throw_away_aln(aln):
            continue
        if aln.query_name not in modified_reads:
            filtered_bam.write(aln)
    bam.close()
    filtered_bam.close()

    print('Merge and sort...')
    bams_to_merge = [f'{out_dir}/filtered.bam', f'{out_dir}/modified_primary.bam']
    pysam.merge("-f", f'{out_dir}/merged.bam', *bams_to_merge)

    pysam.sort("-o", out_file, f'{out_dir}/merged.bam', "-@", "6")
    pysam.index(out_file)

    os.remove(f'{out_dir}/merged.bam')
    os.remove(f'{out_dir}/filtered.bam')
    os.remove(f'{out_dir}/modified_primary.bam')

def fetch_modified_reads(modified_bam, out_dir):
    read_ids = set()
    tmp_out = f'{out_dir}/modified_primary.bam'
    with pysam.AlignmentFile(modified_bam, 'rb') as bam:
        with pysam.AlignmentFile(tmp_out, 'wb', template=bam) as out_bam:
            for aln in bam:
                read_ids.add(aln.query_name)
                if not throw_away_aln(aln):
                    out_bam.write(aln)
    return read_ids

def throw_away_aln(aln):
    # can add more filters
    if aln.is_secondary or aln.is_supplementary:
        return True
    return False

def main(args):
    out_dir=os.path.dirname(args.out_file)
    os.makedirs(out_dir, exist_ok=True)
    filter_and_merge_bam(args.input_bam, 
                         args.modified_bam, 
                         out_dir,
                         args.out_file)

if __name__ == '__main__':
    parser = ArgumentParser(
        formatter_class=ArgumentDefaultsHelpFormatter,
        add_help=False
    )
    parser.add_argument("-b", "--input_bam", help="unmodified input bam file")
    parser.add_argument("-m", "--modified_bam", help="bam file with modified reads only")
    parser.add_argument("-o", "--out_file", help="output file name", default=os.getcwd()+"tweakvar_modified.bam")
    args = parser.parse_args()
    main(args)
```
