## Supplemental File S3 for "MosaicSim: A Novel Mosaic Variant Simulator Reveals Diminishing Returns of Ultra-High Coverage for Mosaic Variant Detection": tweakvarsimulator.py

```
import sys
import pysam
from pysam import depth
import argparse
import os
import numpy as np
from numpy.random import choice, uniform, seed as npseed
from numpy import random as nran
from math import ceil
import random

# Initialize parser
parser = argparse.ArgumentParser(description="TweakVarSimulator: A tool for simulating variants")

# Required arguments
parser.add_argument("-i", "--input", required=True, help="Path to input BAM file")
parser.add_argument("-T", "--reference", required=True, help="Path to reference genome")
parser.add_argument("-o", "--output", required=True, help="Output file prefix")

# Optional arguments
parser.add_argument("-s", "--seed", type=int, required=False, help="Seed for random number generation")
parser.add_argument("-minAFsv", "--minimum_allele_frequency_sv", type=float, required=False)
parser.add_argument("-maxAFsv", "--maximum_allele_frequency_sv", type=float, required=False)
parser.add_argument("-minAFsnv", "--minimum_allele_frequency_snv", type=float, required=False)
parser.add_argument("-maxAFsnv", "--maximum_allele_frequency_snv", type=float, required=False)
parser.add_argument("-numsv", "--number_of_svs", type=int, required=False)
parser.add_argument("-numsnv", "--number_of_snvs", type=int, required=False)
parser.add_argument("-maxsnvl", "--maximum_snv_length", type=int, required=False)
parser.add_argument("-minsvl", "--minimum_sv_length", type=int, required=False)
parser.add_argument("-maxsvl", "--maximum_sv_length", type=int, required=False)
parser.add_argument("-sub", "--substitution_rate", type=float, required=False)
parser.add_argument("-insdelsnv", "--insdel_snv_rate", type=float, required=False)
parser.add_argument("-insdel", "--insdel_sv_rate", type=float, required=False)
parser.add_argument("--ref_chroms", nargs='+', help="Optional list of reference chromosomes")
parser.add_argument("-snv", "--SNV_truth_file", required=False, help="Optional SNV truth VCF file")
parser.add_argument("-sv", "--SV_truth_file", required=False, help="Optional SV truth VCF file")
parser.add_argument("-bps", "--bp_shift", type=int,  required=False, help="Optionally allows you to shift all bp positions up or downstream by the indicated number of bps")
parser.add_argument("-imc", "--ignore_minimum_cov", type=bool, required=False, help="If True, loci will not be evaluated for minimum coverage")

args = parser.parse_args()

bam_path = args.input
ref_path = args.reference
output_prefix = args.output
snv_truth_file = args.SNV_truth_file
sv_truth_file = args.SV_truth_file

bp_shift = args.bp_shift
ignore_minimum_cov = args.ignore_minimum_cov

if ignore_minimum_cov is None:
    ignore_minimum_cov = False

if bp_shift is None:
    bp_shift = 0
else:
    bp_shift = int(bp_shift)

seed = args.seed if args.seed is not None else 0
npseed(seed)
random.seed(seed)

minAFsv = args.minimum_allele_frequency_sv if args.minimum_allele_frequency_sv is not None else 0.01
maxAFsv = args.maximum_allele_frequency_sv if args.maximum_allele_frequency_sv is not None else 0.05
numsv = args.number_of_svs if args.number_of_svs is not None else 50
minsvl = args.minimum_sv_length if args.minimum_sv_length is not None else 50
maxsvl = args.maximum_sv_length if args.maximum_sv_length is not None else 10000
insdel = args.insdel_sv_rate if args.insdel_sv_rate is not None else 0.7

minAFsnv = args.minimum_allele_frequency_snv if args.minimum_allele_frequency_snv is not None else 0.01
maxAFsnv = args.maximum_allele_frequency_snv if args.maximum_allele_frequency_snv is not None else 0.05
numsnv = args.number_of_snvs if args.number_of_snvs is not None else 200
maxsnvl = args.maximum_snv_length if args.maximum_snv_length is not None else 100
sub = args.substitution_rate if args.substitution_rate is not None else 1
insdelsnv = args.insdel_snv_rate if args.insdel_snv_rate is not None else 0.5

SVvcf = f"{output_prefix}_SV.vcf"
SNVvcf = f"{output_prefix}_SNV.vcf"

def count_decimals(x):
    s = str(x)
    if '.' in s:
        return len(s.split('.')[-1].rstrip('0'))
    else:
        return 0

def parse_truth_vcf(vcf_path):
    records = []
    with open(vcf_path, 'r') as f:
        for line in f:
            if line.startswith('#'):
                continue
            parts = line.strip().split('\t')
            chrom, pos, _, ref, alt, _, _, info, *_ = parts
            af_val = 0.05
            for field in info.split(';'):
                if field.startswith('AF='):
                    try:
                        af_val = float(field.split('=')[1])
                    except:
                        pass
            records.append((chrom, pos, ref, alt, af_val))
    return records

def get_chrom_lengths(bam_path, ref_chroms=None):
    with pysam.AlignmentFile(bam_path) as bam:
        chromol = dict(zip(bam.references, bam.lengths))

    if ref_chroms is not None:
        # Filter only if ref_chroms is specified
        chromol = {c: l for c, l in chromol.items() if c in ref_chroms}

    return chromol, tuple(chromol.keys())

def genlocSNV(num,bam_path,mincov=20):
    npseed(seed)
    ref_chroms = args.ref_chroms
    chromol, chrom = get_chrom_lengths(bam_path, ref_chroms=ref_chroms)
    locations=[]
    for i in range(num):
        print(f"Creating SNV {i+1}/{num}...")
        while True:
            ranchrom=nran.choice(chrom)
            loc=str(nran.randint(0,chromol[ranchrom])+bp_shift)
            try:
                cover=int(depth(bam_path,'-r',ranchrom+":"+loc+"-"+loc).rstrip("\n").split("\t")[-1])
            except ValueError:
                continue
            ##Test for minimum coverage requirement
            if ignore_minimum_cov:
                break
            else:
                if cover>=mincov:
                    break
        locations.append(tuple([ranchrom,loc,cover]))
    return tuple(locations)

def genlocSV(num,bam_path,mincov=20):
    npseed(seed)
    ref_chroms = args.ref_chroms
    chromol, chrom = get_chrom_lengths(bam_path, ref_chroms=ref_chroms)
    locations=[]
    for i in range(num):
        print(f"Creating SV {i+1}/{num}...")
        while True:
            ranchrom=nran.choice(chrom)
            loc=str(nran.randint(0,chromol[ranchrom])+bp_shift)
            for i in locations:
                if i[0]==ranchrom and abs(int(i[1]))-int(loc)<50000:
                    continue
            try:
                cover=int(depth(bam_path,'-r',ranchrom+":"+loc+"-"+loc).rstrip("\n").split("\t")[-1])
            except ValueError:
                continue
            ##Test for minimum coverage requirement
            if ignore_minimum_cov:
                break
            else:
                if cover>=mincov:
                    break
        locations.append(tuple([ranchrom,loc,cover]))
    return tuple(locations)

def genseq(minl,maxl):
    npseed(seed)
    nucl=tuple(["A","T","C","G"])
    res=''
    for i in range(choice(range(minl,maxl+1))):
        res+=choice(nucl)
    return res

def gensnps(maxsnvl,sub,insdelsnv=insdelsnv, snplist=[]):
    npseed(seed)
    result=[]
    nucl=["A","T","C","G"]
    for i in range(len(snplist)):
        draw = choice(tuple(['snp','indel']), 1, p= [sub,1-sub])    
        if draw=='snp':
            nucl=["A","T","C","G"]
            nucl.remove(snplist[i][3][0].upper())
            result.append(tuple(list(snplist[i][0:3])+[snplist[i][3][0].upper()]+[choice(nucl,1)[0]]))
        else:
            draw = choice(tuple(['in','del'],), 1, p= [insdelsnv,1-insdelsnv])    
            if draw=='in':
                result.append(tuple(list(snplist[i][0:3])+[snplist[i][3][0].upper()]+[snplist[i][3][0].upper()+genseq(1,maxsnvl-1).upper()]))
            else:
                rmlen=choice(range(maxsnvl+1)[1:])
                result.append(tuple(list(snplist[i][0:3])+[snplist[i][3].upper()[:rmlen]]+[snplist[i][3].upper()[0]]))

    return tuple(result)
def getrefsnp(reffile,snplist=1):
    with open(reffile,'r') as f:
        data=[]
        cta=[]
        for i in snplist:
            cta.append(i[0])
        cta=set(cta)
        start=1
        for line in f:
            if line.startswith(">"):
                if start==1:
                    start=0
                else:
                    if analyse==1:
                        data.append(tuple([chromosome,seq]))
                if "chr"==snplist[0][0][0:3]:
                    chromosome=line.lstrip(">").split(" ")[0].rstrip("\n")
                else:
                    chromosome=line.lstrip(">").split(" ")[0].lstrip("chr").rstrip("\n")
                seq=''
                analyse=0
                if chromosome in cta:
                    analyse=1
            else:
                if analyse==0:
                    continue
                seq+=line.rstrip("\n") 
        if analyse==1:
            data.append(tuple([chromosome,seq]))
        data=tuple(data)

    f.close()
    snplistmod=[]
    for i in snplist:
        for j in data:
            if i[0]==j[0]:
                snplistmod.append(tuple(list(i)+[j[1][int(i[1])-1:int(i[1])+maxsnvl]]))
                break
    return tuple(snplistmod)
    


def genseq(minl,maxl):
    npseed(seed)
    nucl=tuple(["A","T","C","G"])
    res=''
    for i in range(choice(range(minl,maxl+1))):
        res+=choice(nucl)
    return res

def main():
    chromol, chrom = get_chrom_lengths(bam_path)

    # SNV processing
    if snv_truth_file is not None:
        raw_snvloc = parse_truth_vcf(snv_truth_file)

        vcfsnv = [
            '##fileformat=VCFv4.2',
            '##FILTER=<ID=PASS,Description="All filters passed">',
            '##FORMAT=<ID=GT,Number=1,Type=String,Description="Genotype">',
            '##FORMAT=<ID=AD,Number=R,Type=Integer,Description="Read depth for each allele">',
            '##FORMAT=<ID=DV,Number=1,Type=Integer,Description="Number of variant reads">',
            '##INFO=<ID=AF,Number=A,Type=Float,Description="Allele Frequency">'
        ]
        vcfsnv.extend([f'##contig=<ID={c},length={l}>' for c, l in chromol.items()])

        vcfsnv.append('\t'.join(['#CHROM', 'POS', 'ID', 'REF', 'ALT', 'QUAL', 'FILTER', 'INFO', 'FORMAT', 'SAMPLE']))
        snvloc = []
        for i in range(len(raw_snvloc)):
            print(f"Retrieving SNV {i+1}/{len(raw_snvloc)} from truth vcf...")
            chrom, pos, ref, alt, af_val = raw_snvloc[i]
            ##update the position
            pos = str(int(pos)+bp_shift)
            cover = int(depth(bam_path, '-r', chrom + ":" + pos + "-" + pos).rstrip("\n").split("\t")[-1])
            pos = int(pos)
            readnum = ceil(af_val * cover)
            vcfsnv.append(f"{chrom}\t{pos}\t.\t{ref}\t{alt}\t1500\tPASS\tAF={af_val}\tGT:AD:DV\t0/0:{cover-readnum}:{readnum}")

        with open(SNVvcf, "w") as f:
            f.write('\n'.join(vcfsnv))
        f.close()

        print(f"New SNV truth vcf written to {SNVvcf}.")
    else:
        snvloc=genlocSNV(numsnv,bam_path,ceil(1/maxAFsnv))
        random.seed(seed)

        print("Writing the SNV output file...")

        if numsnv > 0:
            vcfsnv=[
                '##fileformat=VCFv4.2',
                '##FILTER=<ID=PASS,Description="All filters passed">',
                '##FORMAT=<ID=GT,Number=1,Type=String,Description="Genotype">',
                '##FORMAT=<ID=AD,Number=R,Type=Integer,Description="Read depth for each allele">',
                '##FORMAT=<ID=DV,Number=1,Type=Integer,Description="Number of variant reads">',
                '##INFO=<ID=AF,Number=A,Type=Float,Description="Allele Frequency">'
            ]

            # Dynamically add contig lines
            vcfsnv.extend([f'##contig=<ID={c},length={l}>' for c, l in chromol.items()])

            # Add VCF column headers
            vcfsnv.append('\t'.join(['#CHROM', 'POS', 'ID', 'REF', 'ALT', 'QUAL', 'FILTER', 'INFO', 'FORMAT', 'SAMPLE']))

            snps=getrefsnp(ref_path,snvloc)
            
            
            for i in gensnps(maxsnvl=maxsnvl,sub=sub, snplist=snps):
                AFnum=round(random.uniform(minAFsnv,maxAFsnv),count_decimals(minAFsnv))
                readnum=ceil(AFnum*i[2])
                vcfsnv.append(str(i[0])+'\t'+str(i[1])+'\t.\t'+str(i[3])+'\t'+str(i[4])+"\t1500\tPASS\tAF="+str(AFnum)+"\tGT:AD:DV\t0/0:"+str(i[2]-readnum)+":"+str(readnum))
            with open(SNVvcf,"w") as f:
                for i in tuple(vcfsnv)[:-1]:
                    f.write(i+'\n')
                f.write(tuple(vcfsnv)[-1])
            f.close()

            print(f"New SNV truth vcf written to {SNVvcf}.")
            

    # SV processing
    if sv_truth_file is not None:

        print(f"Retrieving SVs from truth vcf...")
        svloc = parse_truth_vcf(sv_truth_file)
        vcfsv = [
            '##fileformat=VCFv4.2',
            '##ALT=<ID=INS,Description="Insertion">',
            '##ALT=<ID=DEL,Description="Deletion">',
            '##FORMAT=<ID=GT,Number=1,Type=String,Description="Genotype">',
            '##FORMAT=<ID=GQ,Number=1,Type=Integer,Description="Genotype quality">',
            '##FILTER=<ID=PASS,Description="All filters passed">',
            '##INFO=<ID=PRECISE,Number=0,Type=Flag,Description="Structural variation with precise breakpoints">',
            '##INFO=<ID=SVTYPE,Number=1,Type=String,Description="Type of structural variation">',
            '##INFO=<ID=SVLEN,Number=1,Type=Integer,Description="Length of structural variation">',
            '##INFO=<ID=END,Number=1,Type=Integer,Description="End position of structural variation">',
            '##INFO=<ID=AF,Number=A,Type=Float,Description="Allele Frequency">'
        ]
        vcfsv.extend([f'##contig=<ID={c},length={l}>' for c, l in chromol.items()])
        vcfsv.append('\t'.join(['#CHROM', 'POS', 'ID', 'REF', 'ALT', 'QUAL', 'FILTER', 'INFO', 'FORMAT', 'SAMPLE']))

        for idx, record in enumerate(svloc):
            if sv_truth_file:
                chrom, pos, ref, alt, af = record
            else:
                chrom, pos = record
                ref, alt = 'N', genseq(minsvl, maxsvl) if choice([True, False], p=[insdel, 1-insdel]) else '<DEL>'
                af = round(random.uniform(minAFsv, maxAFsv), count_decimals(minAFsv))
            pos = str(int(pos) + bp_shift)
            svtype = 'INS' if len(alt) > len(ref) else 'DEL'
            svlen = len(alt) - len(ref) if svtype == 'INS' else -(len(ref) - len(alt))
            end = int(pos) + abs(svlen)
            ID = f"SV{idx+1}"
            vcfsv.append(f"{chrom}\t{pos}\t{ID}\t{ref}\t{alt}\t60\tPASS\tPRECISE;SVTYPE={svtype};SVLEN={svlen};END={end};AF={af}\tGT:GQ\t0/0:60")

        with open(SVvcf, "w") as f:
            f.write('\n'.join(vcfsv))

        print(f"New SV truth vcf written to {SVvcf}.")
        

    else:
        svloc=genlocSV(numsv,bam_path,ceil(1/minAFsv))
        chromol, chrom = get_chrom_lengths(bam_path)
        random.seed(seed)

        print("Writing the SV output file...")
        if numsv > 0:
            vcfsv=[
                '##fileformat=VCFv4.2',
                '##ALT=<ID=INS,Description="Insertion">',
                '##ALT=<ID=DEL,Description="Deletion">',
                '##FORMAT=<ID=GT,Number=1,Type=String,Description="Genotype">',
                '##FORMAT=<ID=GQ,Number=1,Type=Integer,Description="Genotype quality">',
                '##FILTER=<ID=PASS,Description="All filters passed">',
                '##INFO=<ID=PRECISE,Number=0,Type=Flag,Description="Structural variation with precise breakpoints">',
                '##INFO=<ID=SVTYPE,Number=1,Type=String,Description="Type of structural variation">',
                '##INFO=<ID=SVLEN,Number=1,Type=Integer,Description="Length of structural variation">',
                '##INFO=<ID=END,Number=1,Type=Integer,Description="End position of structural variation">',
                '##INFO=<ID=AF,Number=A,Type=Float,Description="Allele Frequency">'
            ]

            # Dynamically add contig lines
            vcfsv.extend([f'##contig=<ID={c},length={l}>' for c, l in chromol.items()])

            # Add VCF column headers
            vcfsv.append('\t'.join(['#CHROM', 'POS', 'ID', 'REF', 'ALT', 'QUAL', 'FILTER', 'INFO', 'FORMAT', 'SAMPLE']))

            insertnum=1
            delnum=1
            for i in svloc:
                draw = choice(tuple(['in','del']), 1, p=[insdel,1-insdel])    
                if draw=='in':
                    seq=genseq(minsvl,maxsvl)
                    vcfsv.append(str(i[0])+"\t"+str(i[1])+"\tHackIns"+str(insertnum)+"\tN\t"+seq+"\t60\tPASS\tPRECISE;SVTYPE=INS;SVLEN="+str(len(seq))+";END="+str(int(i[1])+1)+";AF="+str(round(random.uniform(minAFsv,maxAFsv),count_decimals(minAFsv)))+"\tGT:GQ\t0/0:60")
                    #PRECISE;SVTYPE=INS;SVLEN=333;END=748218 AF \t GT:GQ:DR:DV \t   0/0:28:28:5
                    insertnum+=1
                else:
                    dellen=choice(range(minsvl,maxsvl))
                    vcfsv.append(str(i[0])+"\t"+str(i[1])+"\tHackDel"+str(delnum)+"\tN\t<DEL>\t60\tPASS\tPRECISE;SVTYPE=DEL;SVLEN=-"+str(dellen)+";END="+str(int(i[1])+dellen)+";AF="+str(round(random.uniform(minAFsv,maxAFsv),count_decimals(minAFsv)))+"\tGT:GQ\t0/0:60")
                    delnum+=1

            # Ensure parent directories exist
            os.makedirs(os.path.dirname(output_prefix), exist_ok=True)

            with open(SVvcf,"w") as f:
                for i in tuple(vcfsv)[:-1]:
                    f.write(i+'\n')
                f.write(tuple(vcfsv)[-1])
            f.close()
            
            print(f"New SV truth vcf written to {SVvcf}.")
        

if __name__ == "__main__":
    main()
```
