## Supplemental File S3 for "MosaicSim: A Novel Mosaic Variant Simulator Reveals Diminishing Returns of Ultra-High Coverage for Mosaic Variant Detection": vcf_line_parser.py

```
# -*- coding: utf-8 -*-
"""
Created on Wed May 17 10:37:37 2023

@author: HGSC
"""
import numpy as np
class VCFHeader(object):
    # init
    def __init__(self, input_line=None):
        self.SAMPLES = ""
        if input_line is not None:
            # save the original
            self.vcf_header = input_line
            # vcf files have 9 fields + the samples/genomes data
            # split and analysis
            VCF_MANDATORY_FIELDS = 9
            tab_sep_fields = input_line.split("\t")
            self.ERROR = len(tab_sep_fields) < VCF_MANDATORY_FIELDS   # 9 VCF fields + genotypes
            if not self.ERROR:
                # extract mandatory fields
                self.SAMPLES = tab_sep_fields[VCF_MANDATORY_FIELDS:]
        else:
            self.ERROR = True

class VCFLineSV(object):
    # init
    def __init__(self, input_line):
        # save the original
        self.sv_line = input_line
        # vcf files have 9 fields + the samples/genomes data
        # split and analysis
        VCF_MANDATORY_FIELDS = 9
        tab_sep_fields = input_line.split("\t")
        self.ERROR = len(tab_sep_fields) < VCF_MANDATORY_FIELDS   # 9 VCF fields + genotypes
        if not self.ERROR:
            # extract mandatory fields
            # not used REF, as _
            [self.CHROM, POS, self.ID, _, ALT, self.QUAL, self.FILTER, INFO,
             FORMAT] = tab_sep_fields[:VCF_MANDATORY_FIELDS]
            self.POS = int(POS)
            self.ALT=ALT
            # for INFO -> get_parsed_info()
            self.BREAKPOIN = ""
            self.SVTYPE = ""
            self.SVLEN = 0
            self.END = -1
            self.SUPPORT = 0
            self.COVERAGE = []
            self.STRAND = ""
            self.STDEV_LEN = -1
            self.STDEV_POS = -1
            self.RNAMES = []
            self.SUPPORT_LONG = 0
            self.AF = "NA"
            # for FORMAT -> get_genotype()
            self.GENOTYPE = ""
            self.GQ = ""
            self.DR = 0
            self.DV = 0
            # parse data
            self.get_genotype(tab_sep_fields[VCF_MANDATORY_FIELDS], FORMAT)
            self.get_parsed_info(INFO)
            # for Translocation (BND/TRA)
            #self.AF = self.DV/(self.DV+self.DR) if (self.AF == "NA" and self.DV+self.DR != 0) else self.AF
            self.TRA = "" if self.SVTYPE != "BND" else ALT
            self.END = self.END if self.SVTYPE != "BND" else self.POS + 1
            self.SVLEN = int(self.SVLEN) if self.SVTYPE != "BND" else self.SVLEN
    def get_parsed_info(self, info_string):
        # INFO field extraction
        extract_info = ["SVTYPE", "SVLEN", "END", "SUPPORT", "COVERAGE", "STRAND", "STDEV_LEN", 
                        "STDEV_POS", "RNAMES", "SUPPORT_LONG", "AF"]
        for each_info in info_string.split(";"):
            if "=" not in each_info:
                self.BREAKPOIN = each_info
            else:
                [info_key, info_val] = each_info.split("=")
                if info_key in extract_info:
                    self.SVTYPE = info_val if info_key == "SVTYPE" else self.SVTYPE
                    self.SVLEN = info_val if info_key == "SVLEN" else self.SVLEN
                    self.END = int(info_val) if info_key == "END" else self.END
                    self.SUPPORT = int(info_val) if info_key == "SUPPORT" else self.SUPPORT
                    #self.COVERAGE = [int(x) for x in info_val.split(",")] if info_key == "COVERAGE" else self.COVERAGE
                    self.STRAND = info_val if info_key == "STRAND" else self.STRAND
                    self.AF = float(info_val) if info_key == "AF" else self.AF
                    self.STDEV_LEN = float(info_val) if info_key == "STDEV_LEN" else self.STDEV_LEN
                    self.STDEV_POS = float(info_val) if info_key == "STDEV_POS" else self.STDEV_POS
                    self.RNAMES = info_val.split(",") if info_key == "RNAMES" else self.RNAMES
                    self.SUPPORT_LONG = int(info_val) if info_key == "SUPPORT_LONG" else self.SUPPORT_LONG

    # extract information from the "FORMAT" field and each sample
    def get_genotype(self, _sample, _format):
        # FORMAT field and genotype extraction
        extract_genotype = ["GT", "GQ", "DR", "DV"]
        # single individual in the vcf file
        split_format = _format.split(":")
        split_gt = _sample.split(":")
        for gt_format, gt_value in zip(split_format, split_gt):
            if gt_format in extract_genotype:
                self.GENOTYPE = gt_value if gt_format == "GT" else self.GENOTYPE
                self.GQ = int(gt_value) if gt_format == "GQ" else self.GQ
                self.DR = int(gt_value) if gt_format == "DR" else self.DR
                self.DV = int(gt_value) if gt_format == "DV" else self.DV
class VCFSampleGenotype(object):
    def __init__(self):
        self.gt = ""
        self.dr = 0
        self.dv = 0
        self.af = 0
        self.id = ""
        self.name = ""
        self.is_mosaic = False

    def set_gt(self, this_gt):
        self.gt = this_gt

    def set_dr(self, this_dr):
        self.dr = int(this_dr)

    def set_dv(self, this_dv):
        self.dv = int(this_dv)

    def set_af(self, this_af):
        self.af = float(this_af)

    def set_id(self, this_id):
        self.id = this_id

    def set_name(self, this_name):
        self.name = this_name

    def set_mosaic(self, this_mosaic_status):
        self.is_mosaic = this_mosaic_status

class VCFSampleGenotype(object):
    def __init__(self):
        self.gt = ""
        self.dr = 0
        self.dv = 0
        self.af = 0
        self.id = ""
        self.name = ""
        self.is_mosaic = False

    def set_gt(self, this_gt):
        self.gt = this_gt

    def set_dr(self, this_dr):
        self.dr = int(this_dr)

    def set_dv(self, this_dv):
        self.dv = int(this_dv)

    def set_af(self, this_af):
        self.af = float(this_af)

    def set_id(self, this_id):
        self.id = this_id

    def set_name(self, this_name):
        self.name = this_name

    def set_mosaic(self, this_mosaic_status):
        self.is_mosaic = this_mosaic_status


# vcf line class for multi individual/population file
class VCFLineSVPopulation(object):
    # init
    def __init__(self, input_line, sample_header_names=None):
        # save the original
        self.svline = input_line
        # vcf files have 9 fields + the sample(s)/genome(s) data
        self.VCF_MANDATORY_FIELDS = 9
        tab_sep_fields = input_line.split("\t")
        self.ERROR = len(tab_sep_fields) < self.VCF_MANDATORY_FIELDS   # 9 VCF fields + genotypes
        # FILTERS for sniffles2
        self.af_min_mosaic = 0.05
        self.af_max_mosaic = 0.20
        if not self.ERROR:
            # extract mandatory fields
            # not used (_): REF, QUAL
            [self.CHROM, POS, self.ID, _, ALT, _, self.FILTER, 
             INFO, FORMAT] = tab_sep_fields[:self.VCF_MANDATORY_FIELDS]
            self.SAMPLES = tab_sep_fields[self.VCF_MANDATORY_FIELDS:]
            self.POS = int(POS)
            # for INFO -> get_parsed_info()
            self.BREAKPOINT = ""
            self.SVTYPE = ""
            self.SVLEN = 0
            self.END = -1
            self.SUPPORT = 0
            self.COVERAGE = []
            self.STRAND = ""
            self.STDEV_LEN = -1
            self.STDEV_POS = -1
            self.RNAMES = []
            self.SUPPORT_LONG = 0
            self.AF = "NA"
            self.SUPP_VEC = ""
            self.N_SUPP_VEC = 0
            self.SUPP_VEC_BOOL_LIST = []
            # for FORMAT -> get_genotype()
            # self.GENOTYPE = ""
            self.samples_GT = []
            self.samples_DR = []
            self.samples_DV = []
            self.samples_AF = []
            self.samples_SV_ID = []
            self.sample_mosaic_status = []
            self.samples_obj = []
            # extract INFO and FORMAT
            self.get_parsed_info(INFO)
            self.get_genotype(FORMAT, sample_header_names)
            # post-procesign
            self.TRA = "" if self.SVTYPE != "BND" else ALT
            self.SVLEN = int(self.SVLEN) if self.SVTYPE != "BND" else 1
            self.END = int(self.END) if self.SVTYPE != "BND" else self.POS+1

    def get_parsed_info(self, info_string):
        extract_info = ["SVTYPE", "SVLEN", "END", "SUPPORT", "COVERAGE", "STRAND", "STDEV_LEN", 
                        "STDEV_POS", "RNAMES", "SUPPORT_LONG", "SUPP_VEC"]
        for each_info in info_string.split(";"):
            if "=" not in each_info:
                self.BREAKPOIN = each_info
            else:
                [info_key, info_val] = each_info.split("=")
                if info_key in extract_info:
                    if info_key == "SVTYPE":
                        self.SVTYPE = info_val
                    elif info_key == "SVLEN":
                        self.SVLEN = info_val
                    elif info_key == "END":
                        self.END = int(info_val)
                    elif info_key == "SUPPORT":
                        self.SUPPORT = int(info_val)
                    # elif info_key == "COVERAGE":
                        # self.COVERAGE = [int(x) if x is not None else 0 for x in info_val.split(",")]
                    elif info_key == "STRAND":
                        self.STRAND = info_val
                    elif info_key == "STDEV_LEN":
                        self.STDEV_LEN = float(info_val)
                    elif info_key == "STDEV_POS":
                        self.STDEV_POS = float(info_val)
                    elif info_key == "RNAMES":
                        self.RNAMES = info_val.split(",")
                    elif info_key == "SUPPORT_LONG":
                        self.SUPPORT_LONG = int(info_val)
                    elif info_key == "SUPP_VEC":
                        self.SUPP_VEC = info_val
                        self.SUPP_VEC_BOOL_LIST = [supp == "1" for supp in list(info_val)]
                        self.N_SUPP_VEC = sum([int(supp) for supp in list(info_val)])
                    else:
                        pass

    def get_genotype(self, _format, _sample_names=None):
        split_format = _format.split(":")
        # each individual in the population vcf file
        for each_gt in self.SAMPLES:
            sample_gt = VCFSampleGenotype()
            split_gt = each_gt.split(":")
            for gt_format, gt_value in zip(split_format, split_gt):
                if gt_format == "GT":
                    self.samples_GT.append(gt_value)
                    sample_gt.set_gt(gt_value)
                elif gt_format == "DR":
                    self.samples_DR.append(gt_value)
                    sample_gt.set_dr(gt_value)
                elif gt_format == "DV":
                    self.samples_DV.append(gt_value)
                    sample_gt.set_dv(gt_value)
                elif gt_format == "ID":
                    self.samples_SV_ID.append(gt_value)
                    sample_gt.set_id(gt_value)
                else:
                    pass
            #AF
            if sample_gt.dv+sample_gt.dr > 0:
                sample_gt.set_af(sample_gt.dv/(sample_gt.dv+sample_gt.dr))
            else:
                sample_gt.set_af(np.nan)
            self.samples_AF.append(sample_gt.af)
            # Mosaic
            sample_gt.set_mosaic(self.af_min_mosaic <= sample_gt.af <= self.af_max_mosaic)
            self.sample_mosaic_status.append(sample_gt.is_mosaic)
            # Name 
            if _sample_names is not None:
                sample_gt.set_name(_sample_names[self.SAMPLES.index(each_gt)])
            # Obj
            self.samples_obj.append(sample_gt)


# vcf line class for single individuals SV
class VCFLineSurvivor(object):
    def __init__(self, input_line):
        pass
        
```
