## Supplementary figures and images for "MosaicSim: A Novel Mosaic Variant Simulator Reveals Diminishing Returns of Ultra-High Coverage for Mosaic Variant Detection"

### Alexander Adam.jpg

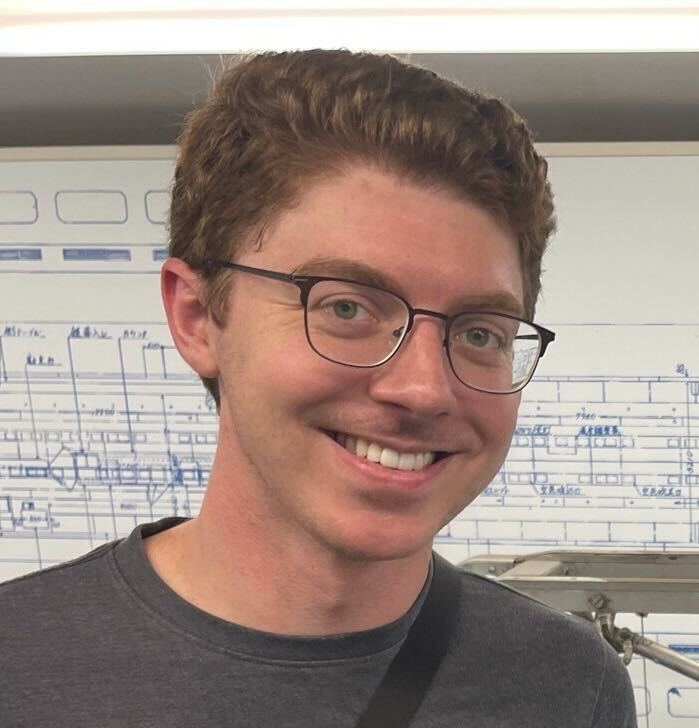

### BackgroundMV.png

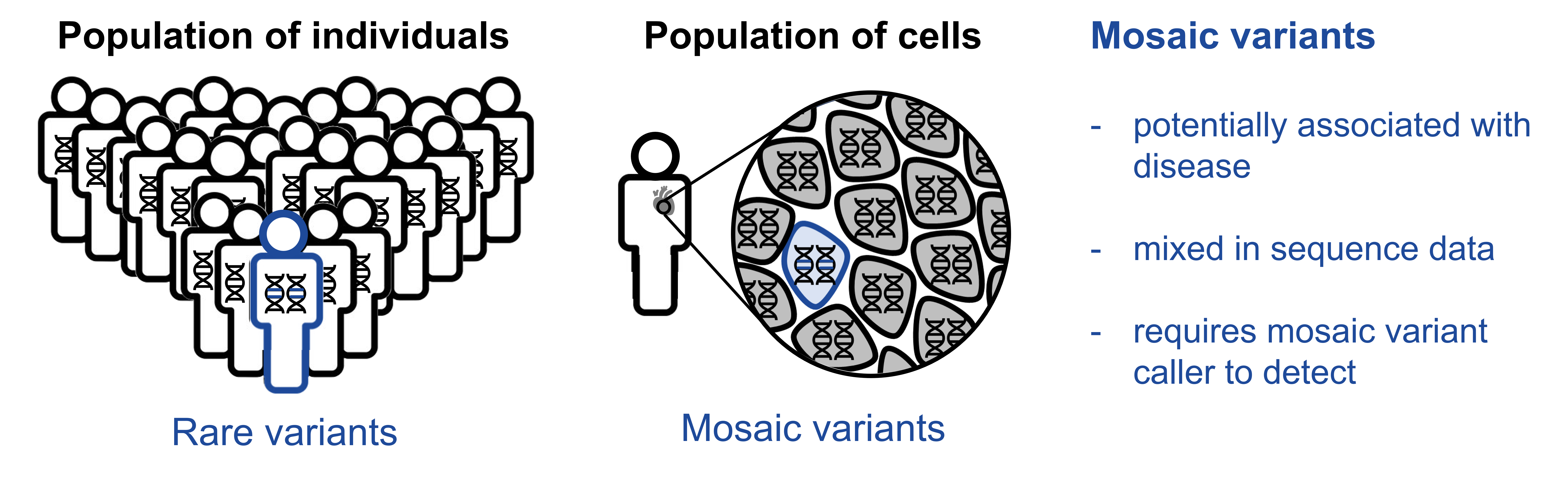

### Chi-Lam Poon.jpg

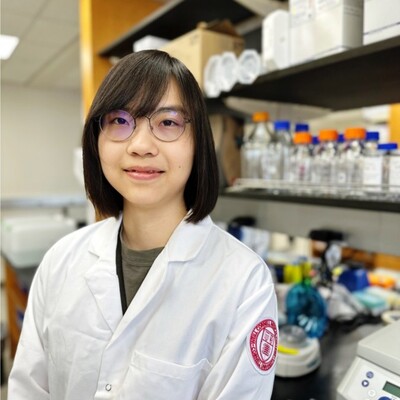

### Divya Kalrai_placeholder.jpg

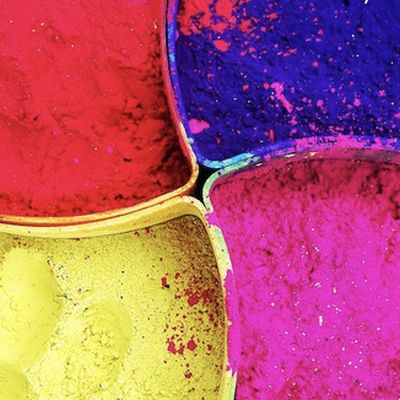

### Erik Stricker.jpg

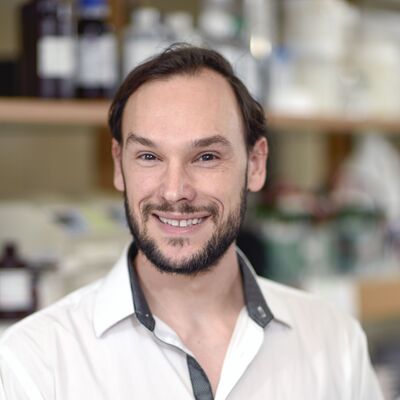

### Farhang Jaryani.png

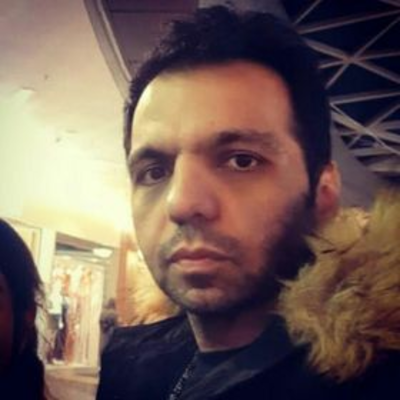

### Fritz Sedlazeck.jpg

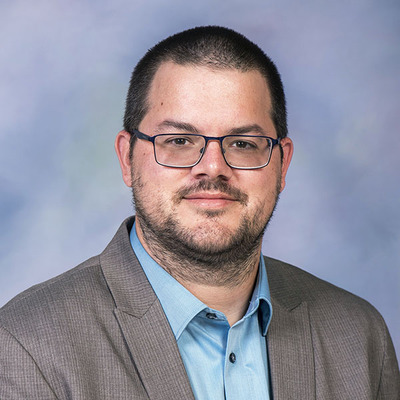

### Joyjit Daw.png

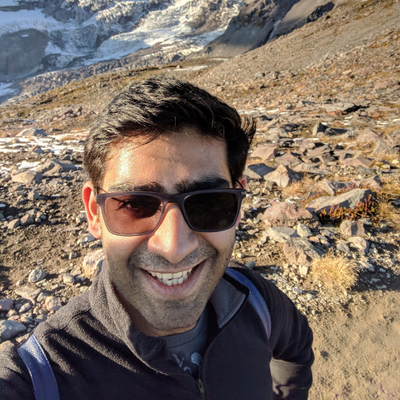

### Michal Bogumil Izydorczyk.png

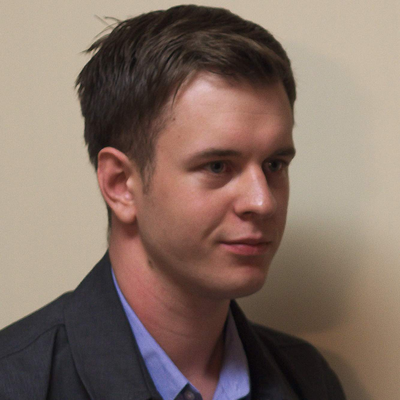

### MosaicSim_workflows.png

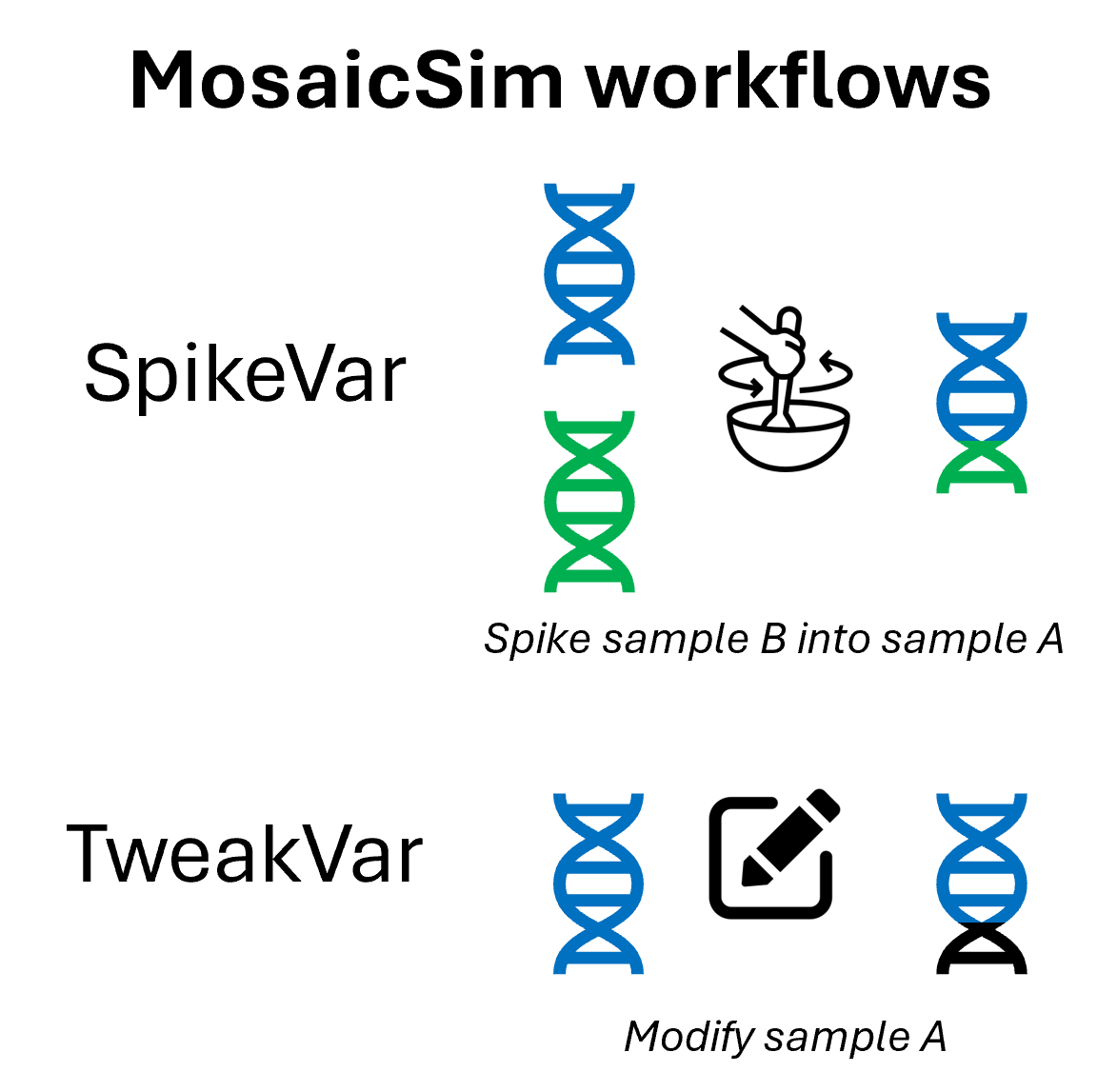

### Philippe Sanio.jpg

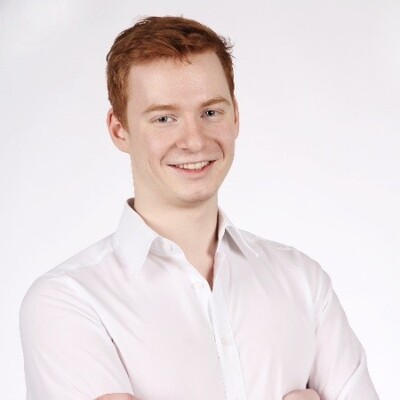

### Sontosh K Deb.jpg

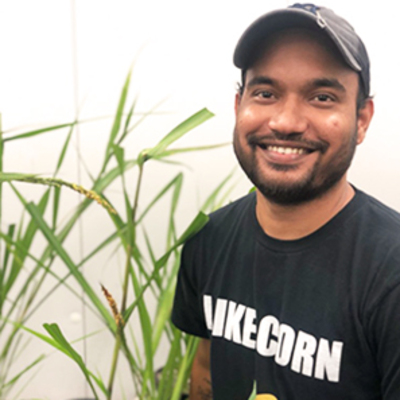

### Spike and Tyke image4.jpg

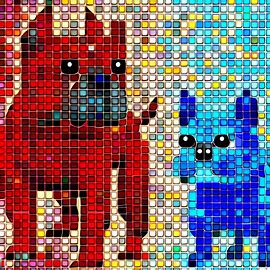

### Spike_screenshot_sv2_ins.png

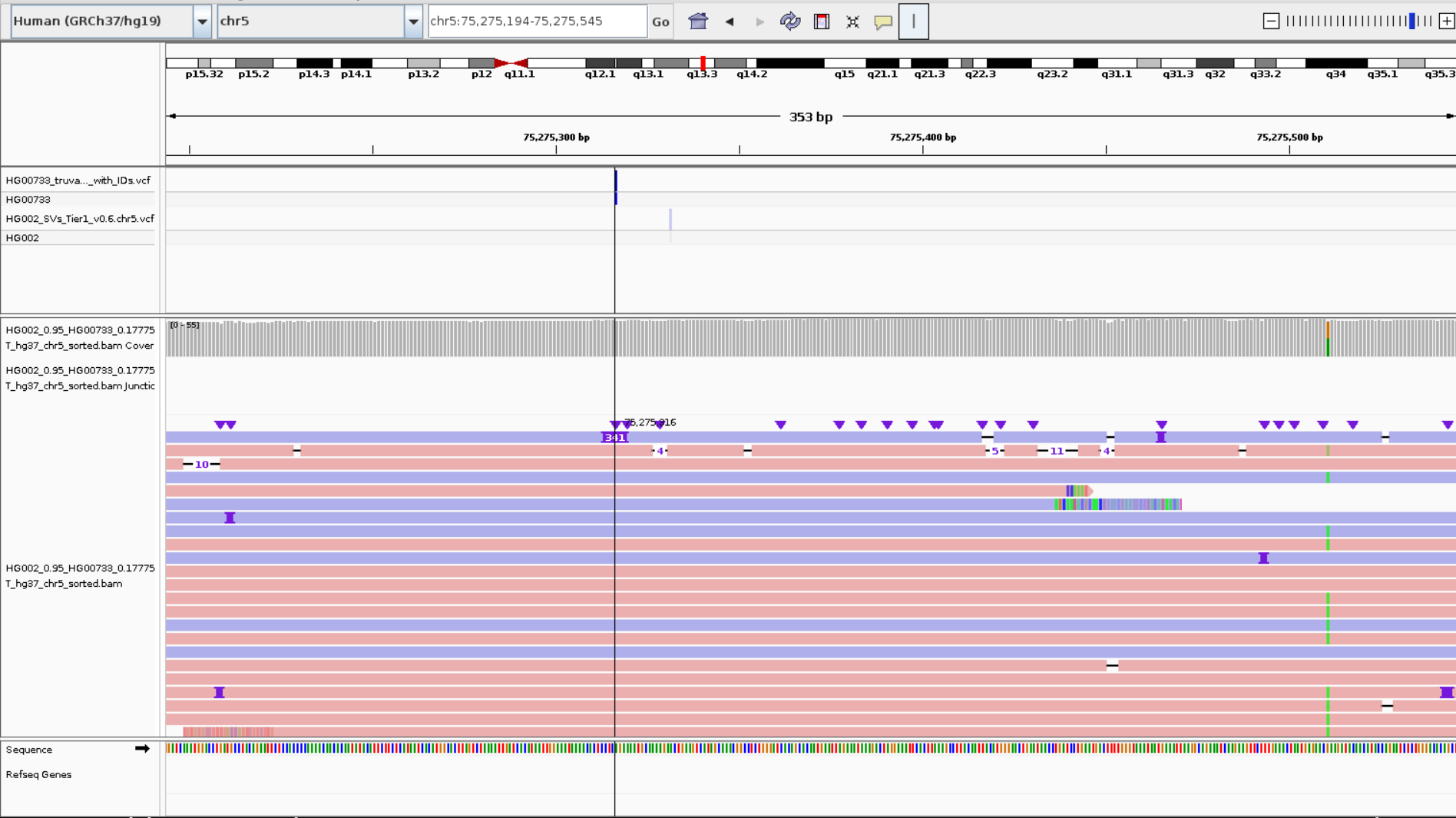

### Spike_screenshot_sv.png

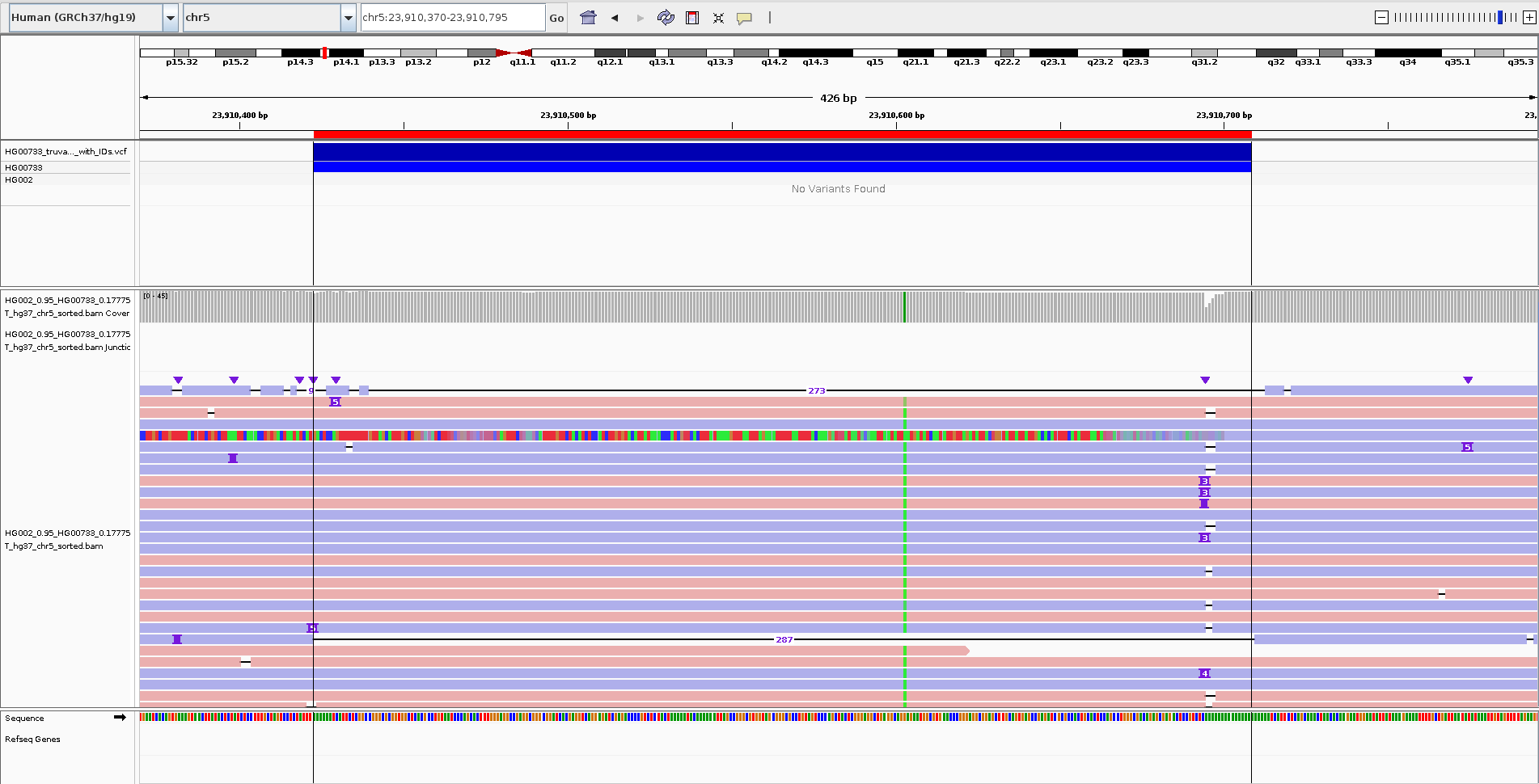

### SpikeVarDatabaseCreator.png

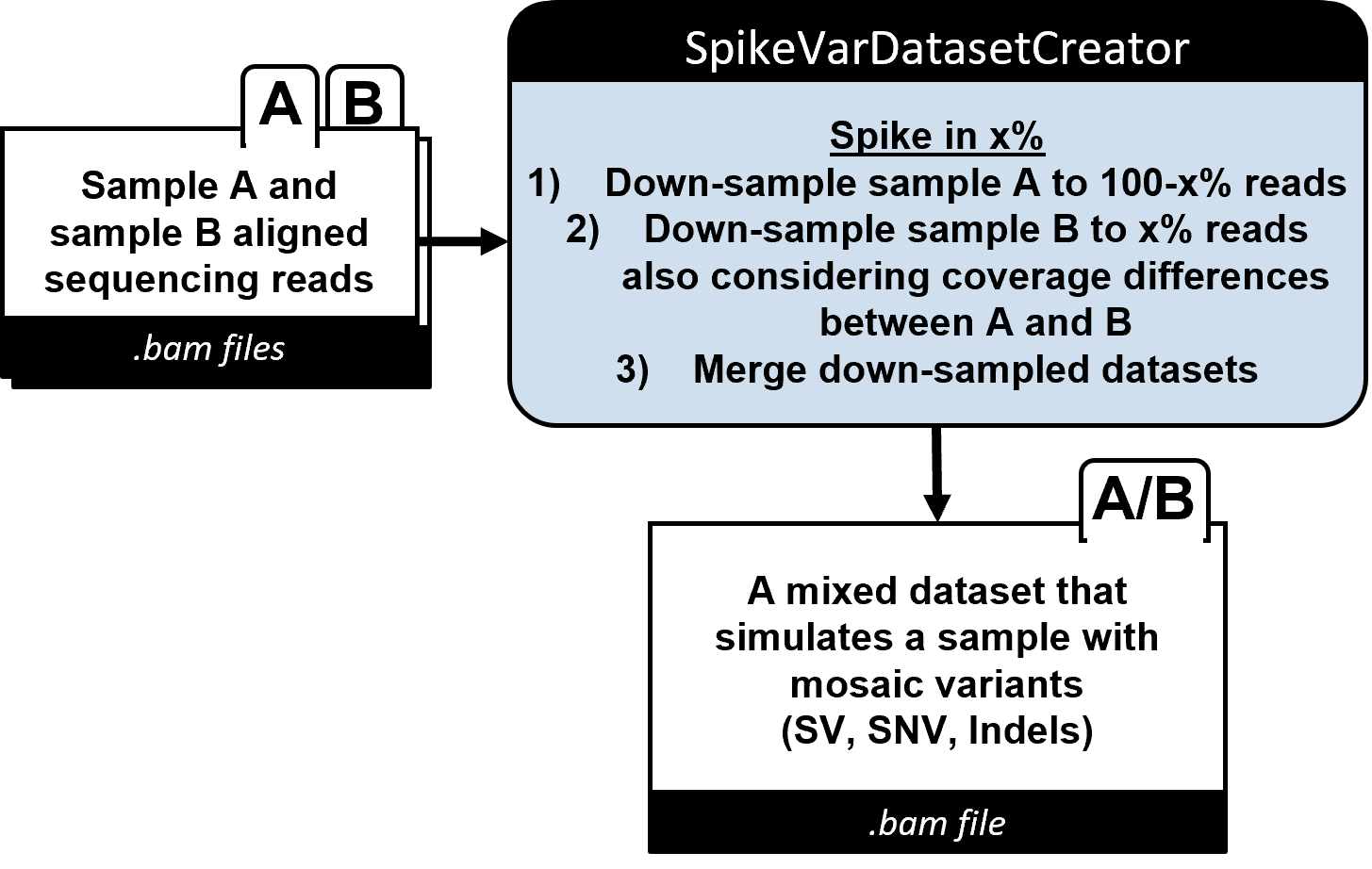

### SpikeVarflowchart_updated.png

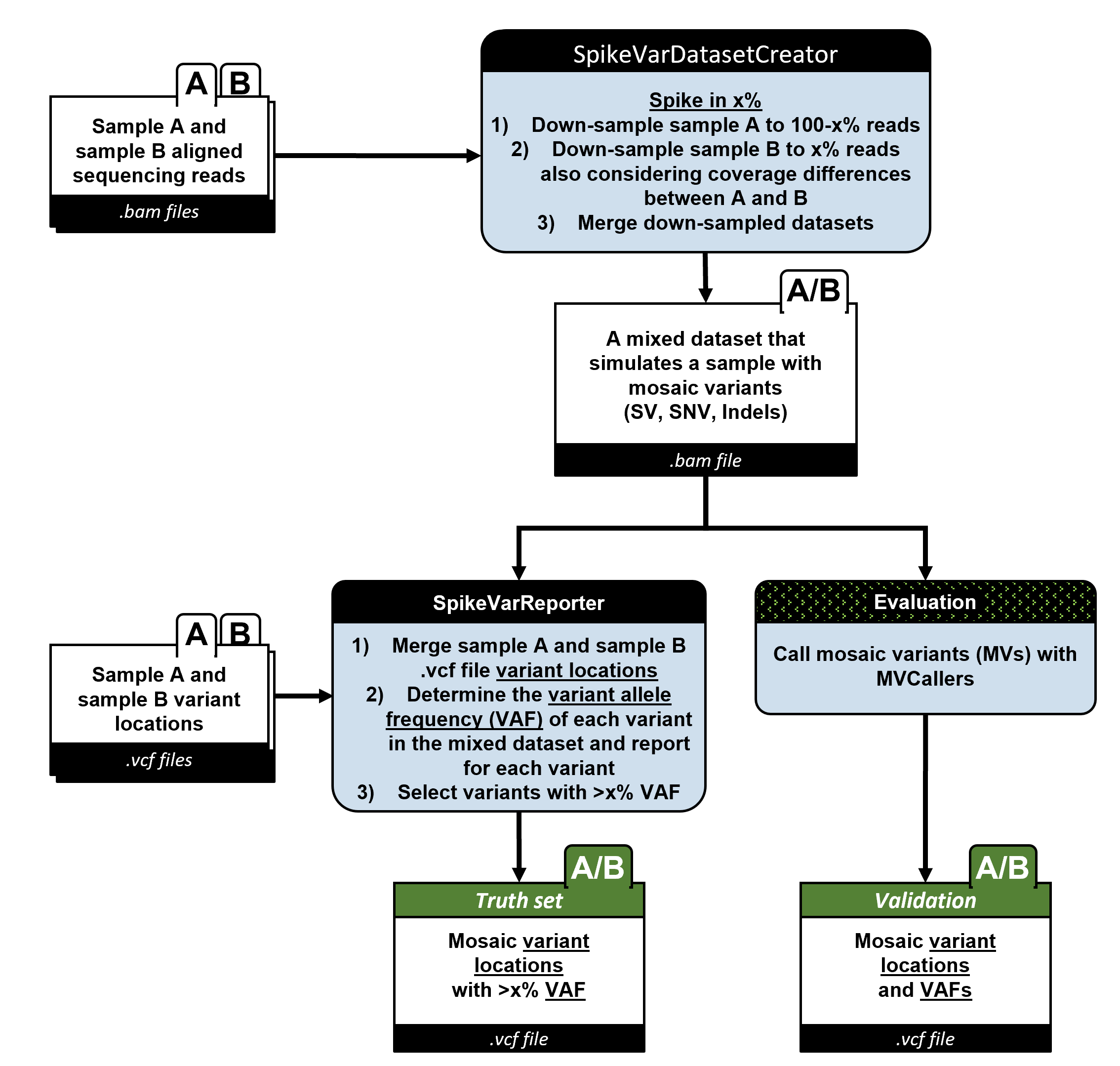

### SpikeVarReporter.png

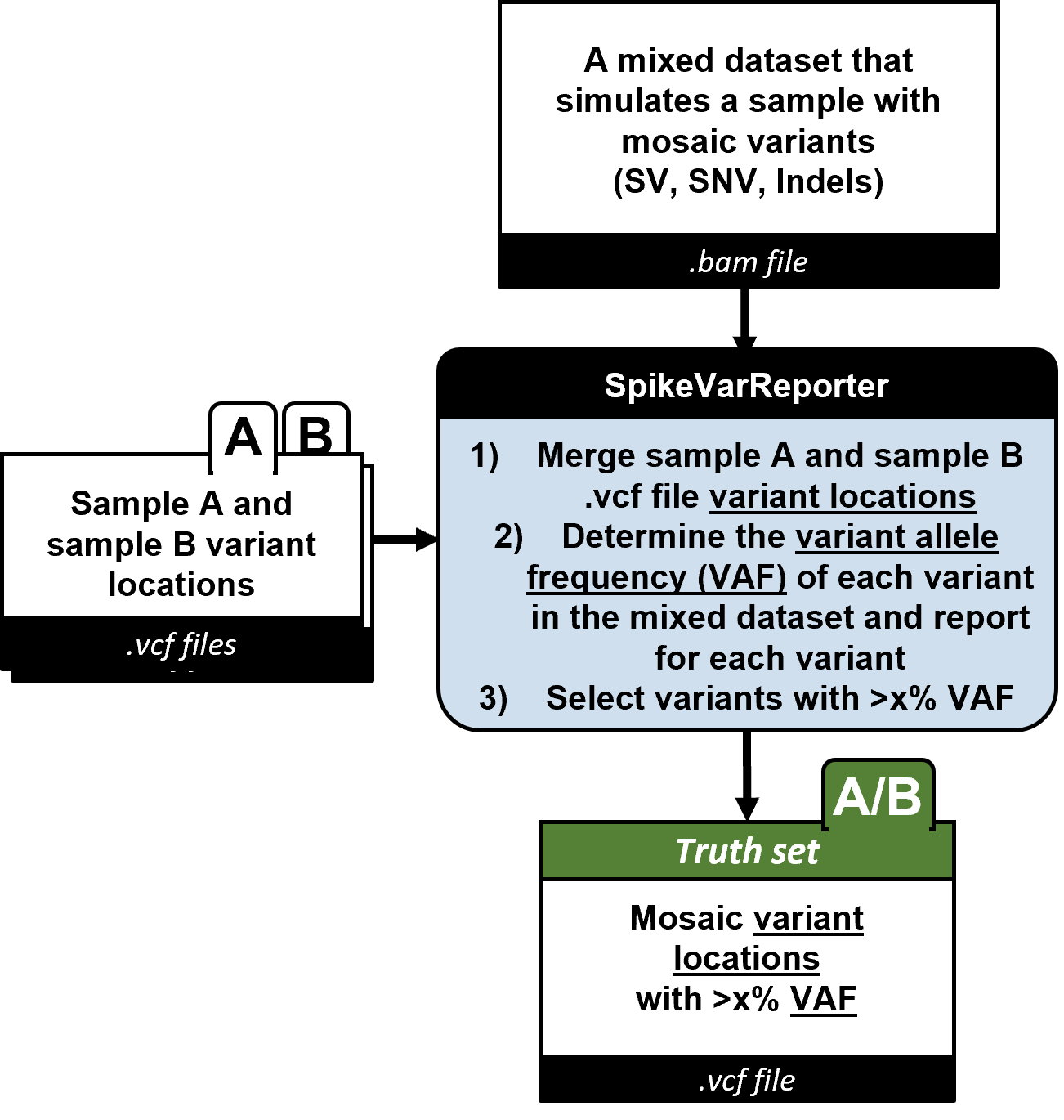

### TweakVar_flowchart_updated.png

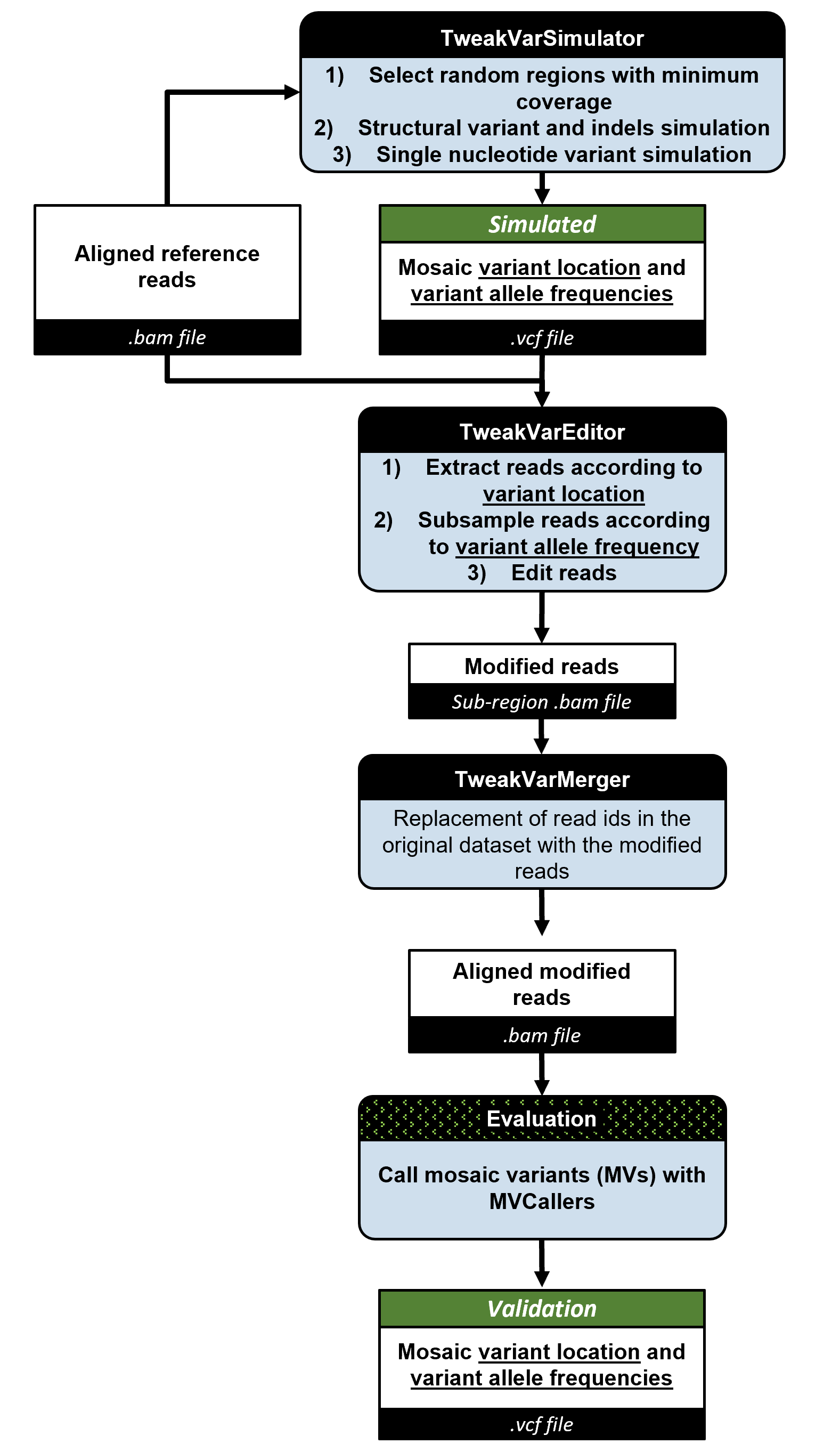

### TweakVar_workflow_simple.png

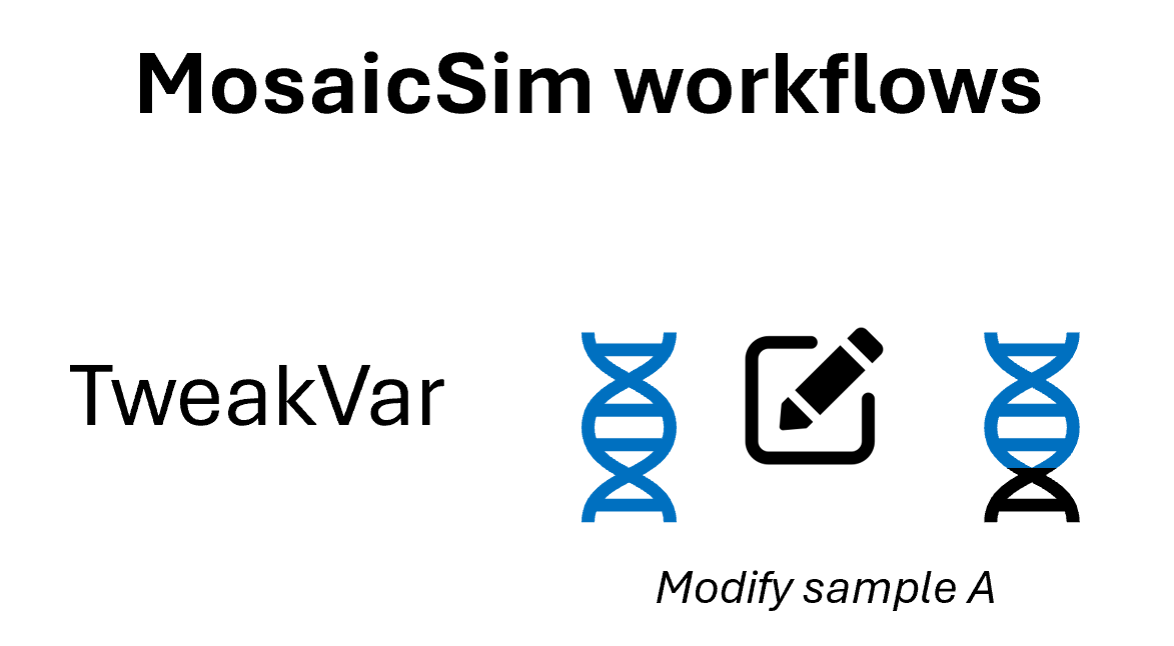

### TweakVarEditor.png

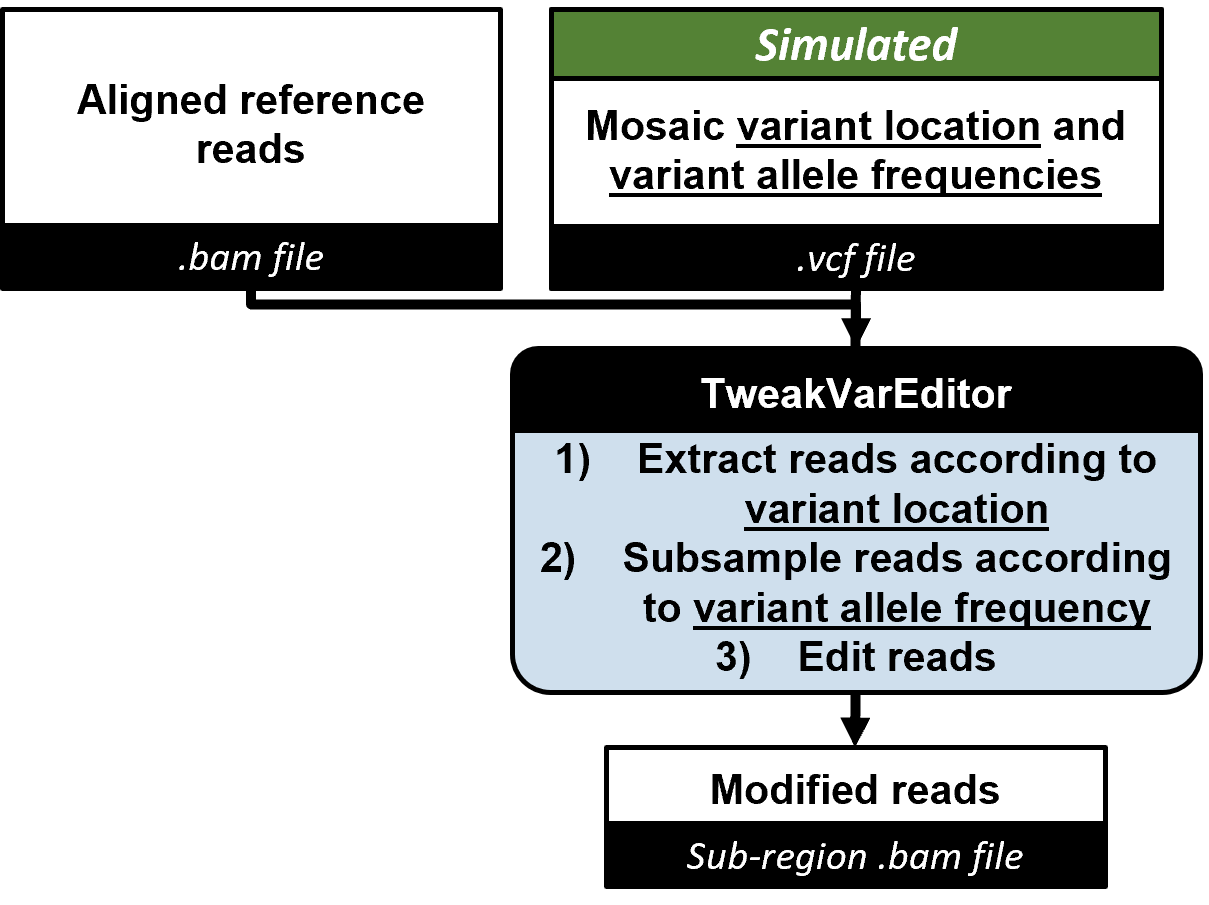

### TweakVarMerger.png

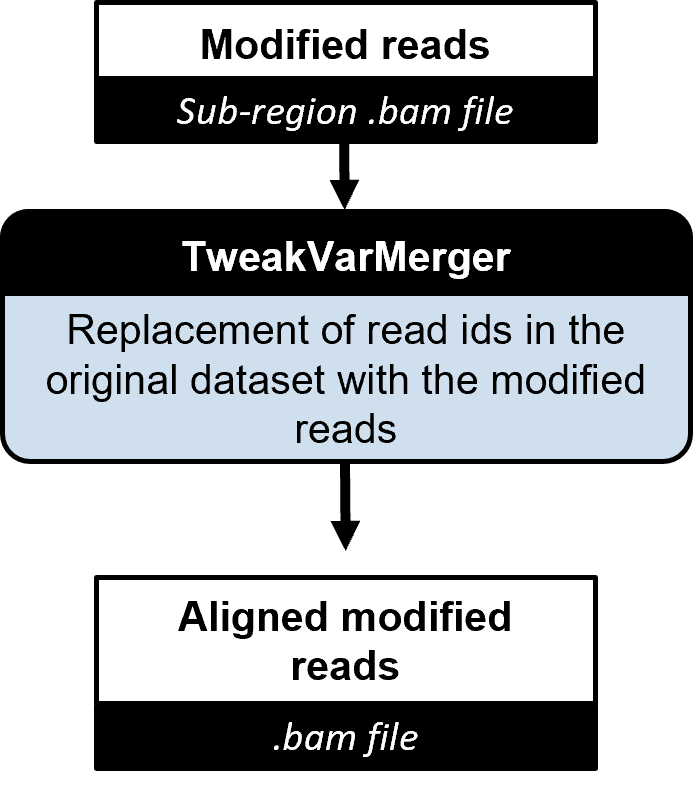

### TweakVarMosaicDel.png

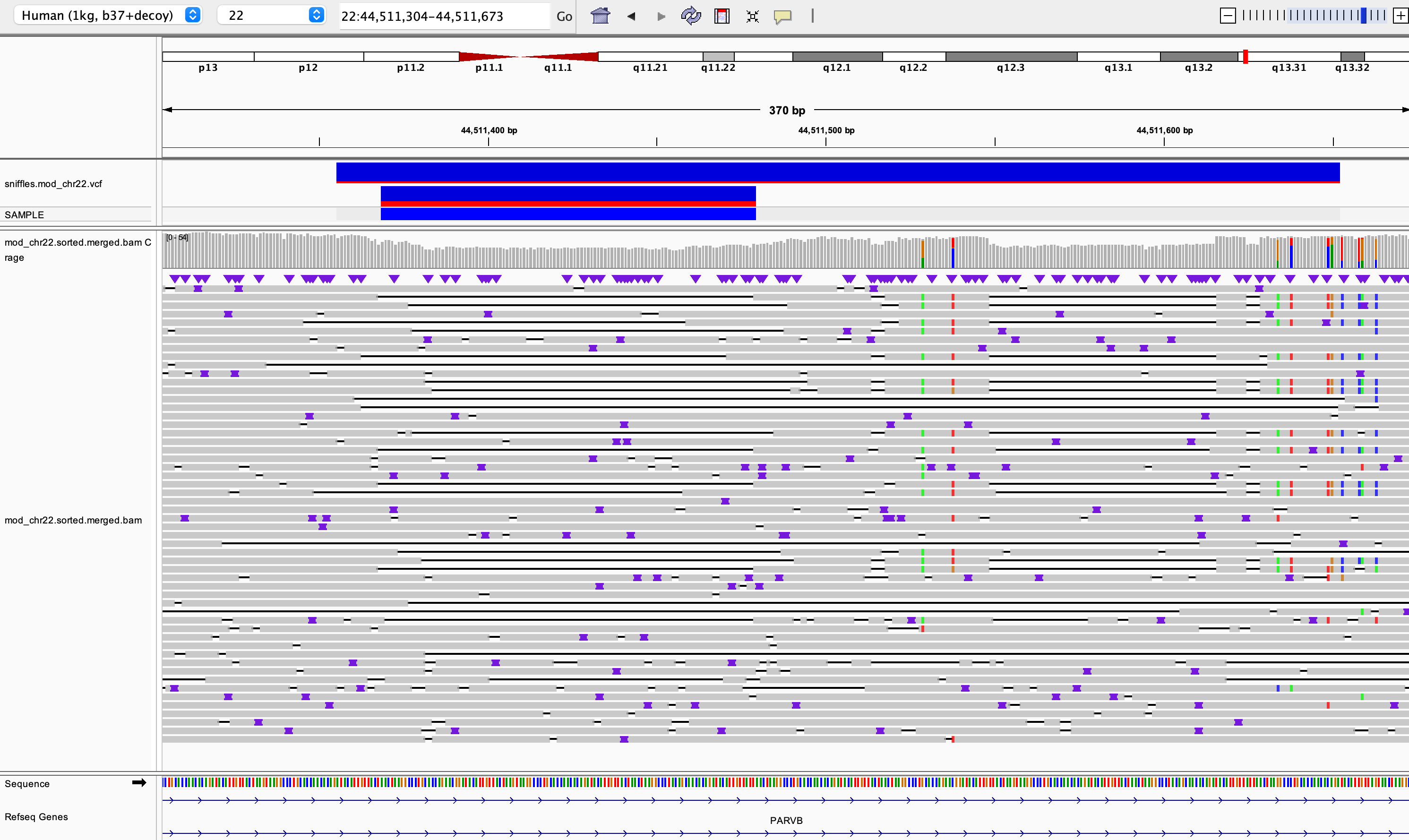

### TweakVarMosaicInsert.png

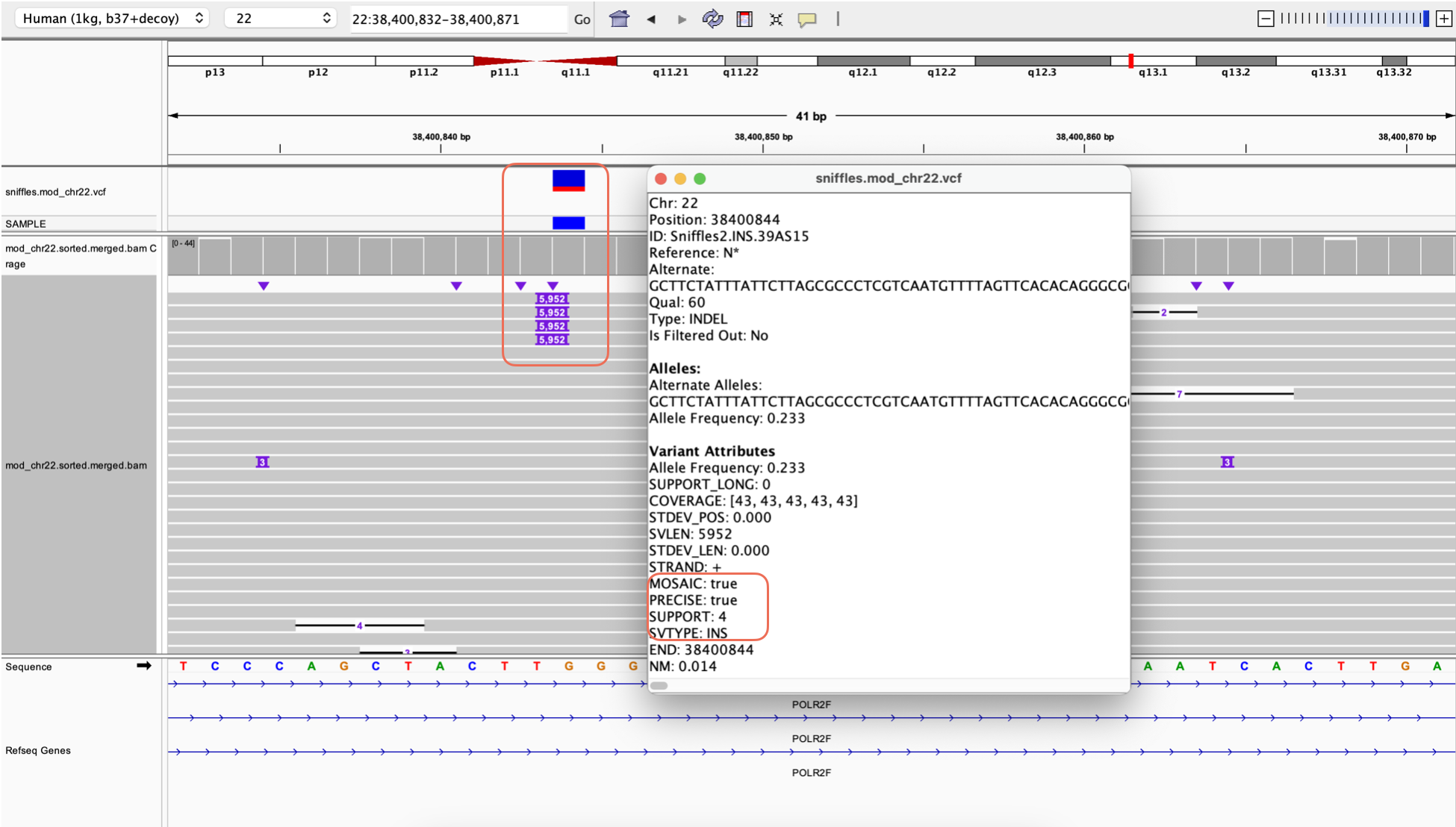

### TweakVarMosaicSNP.png

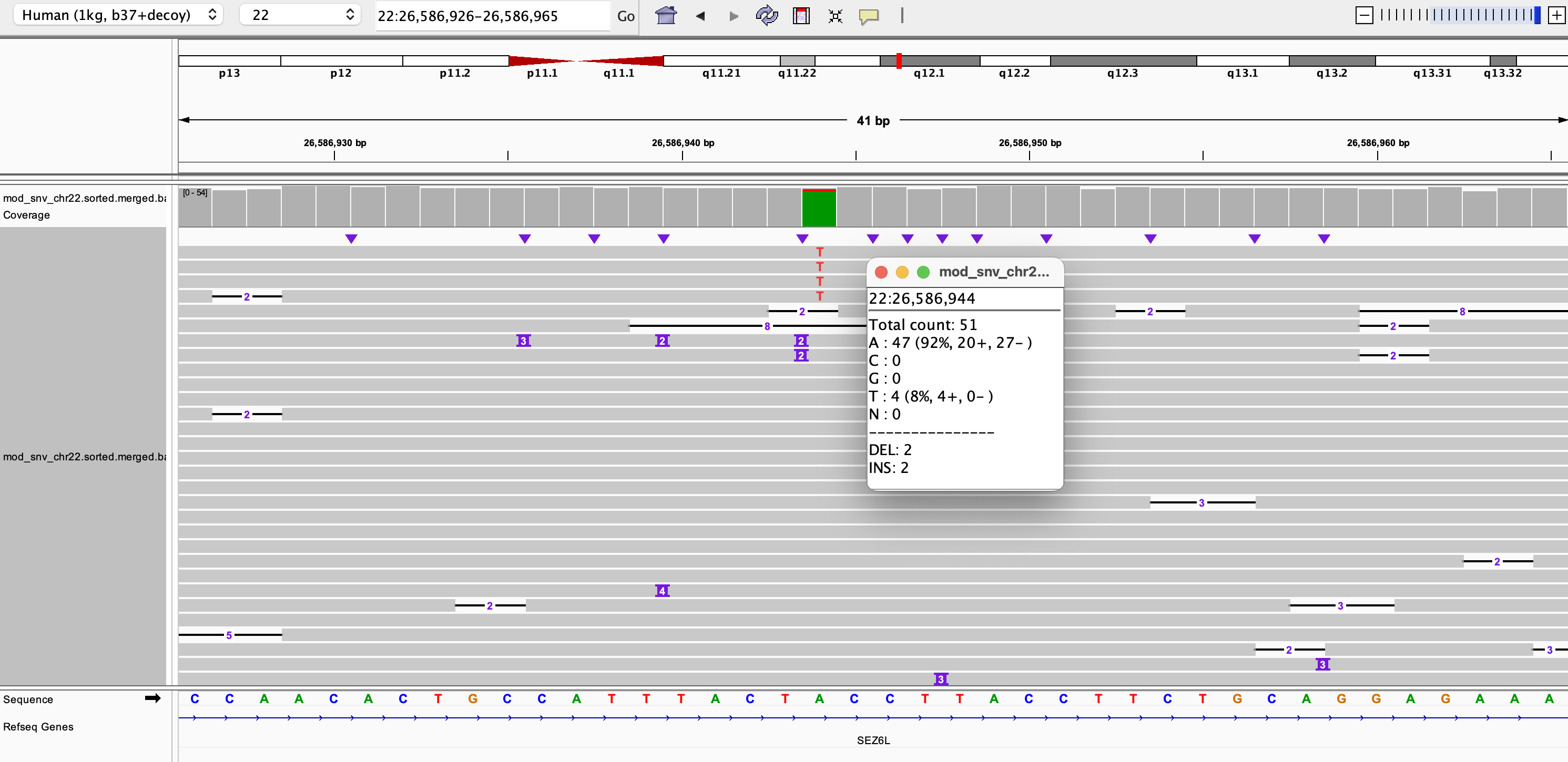

### TweakVarMosaicSNV1.png

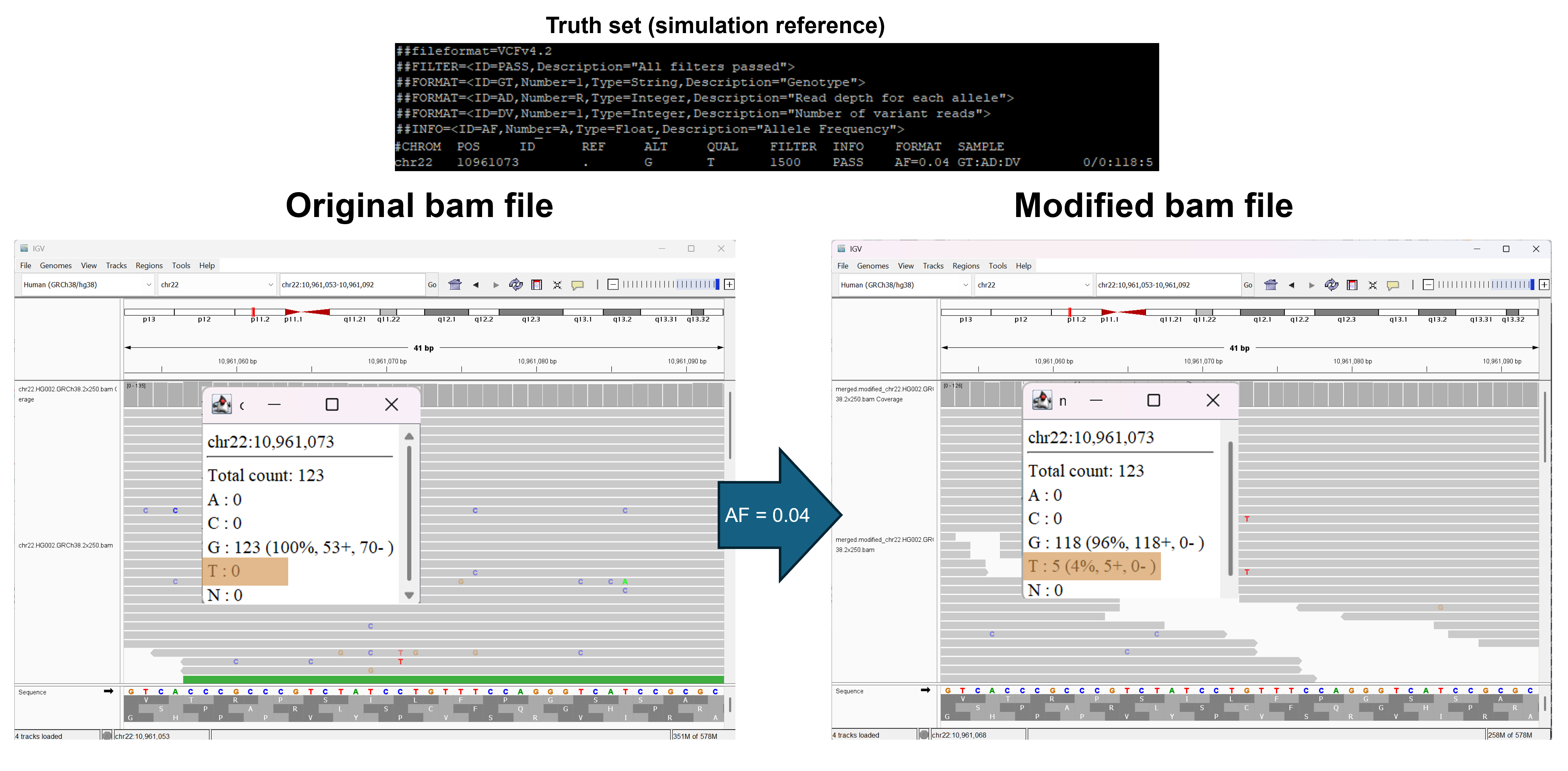

### TweakVarMosaicSNV2.png

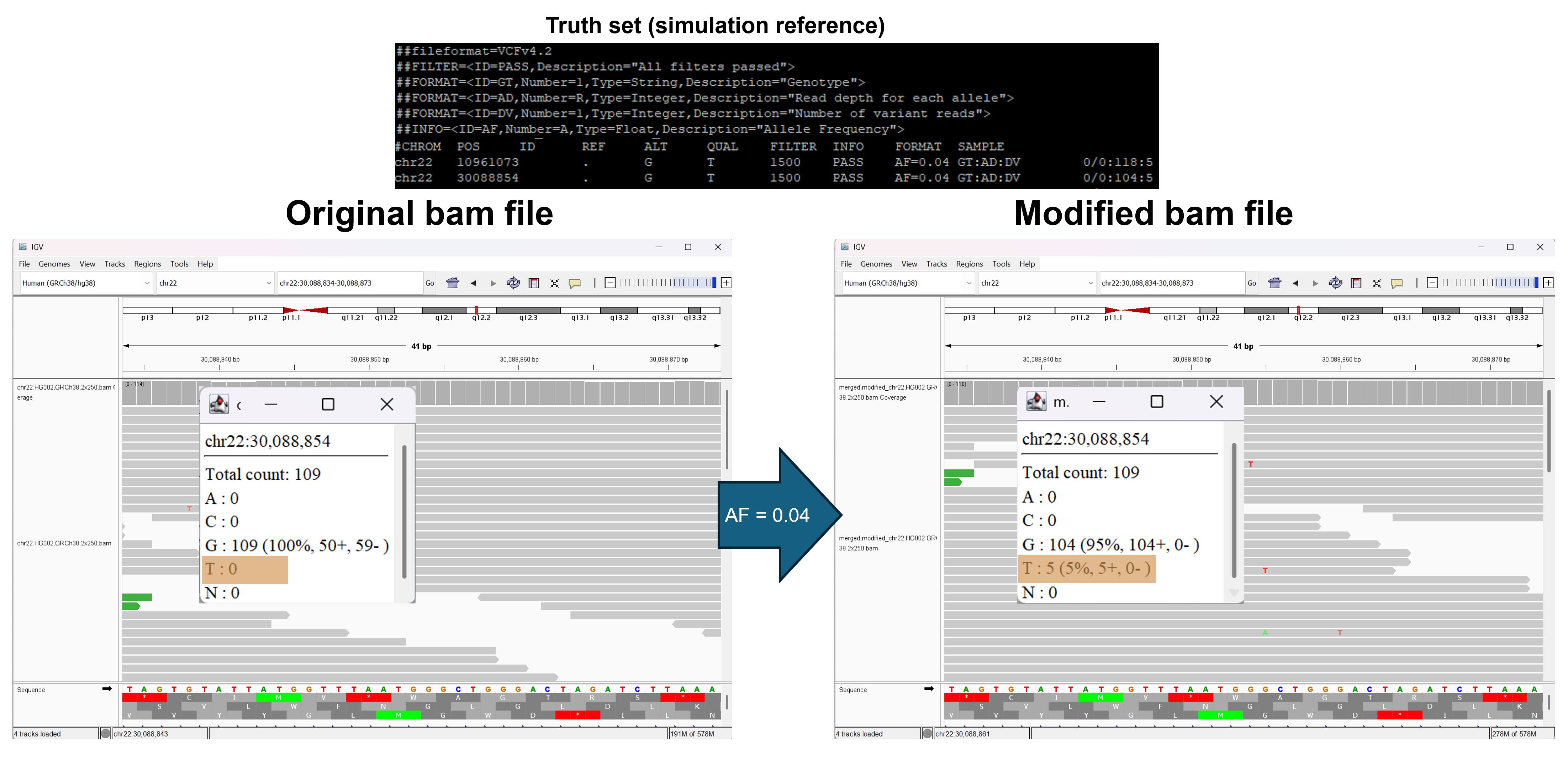

### TweakVarSimulator.png

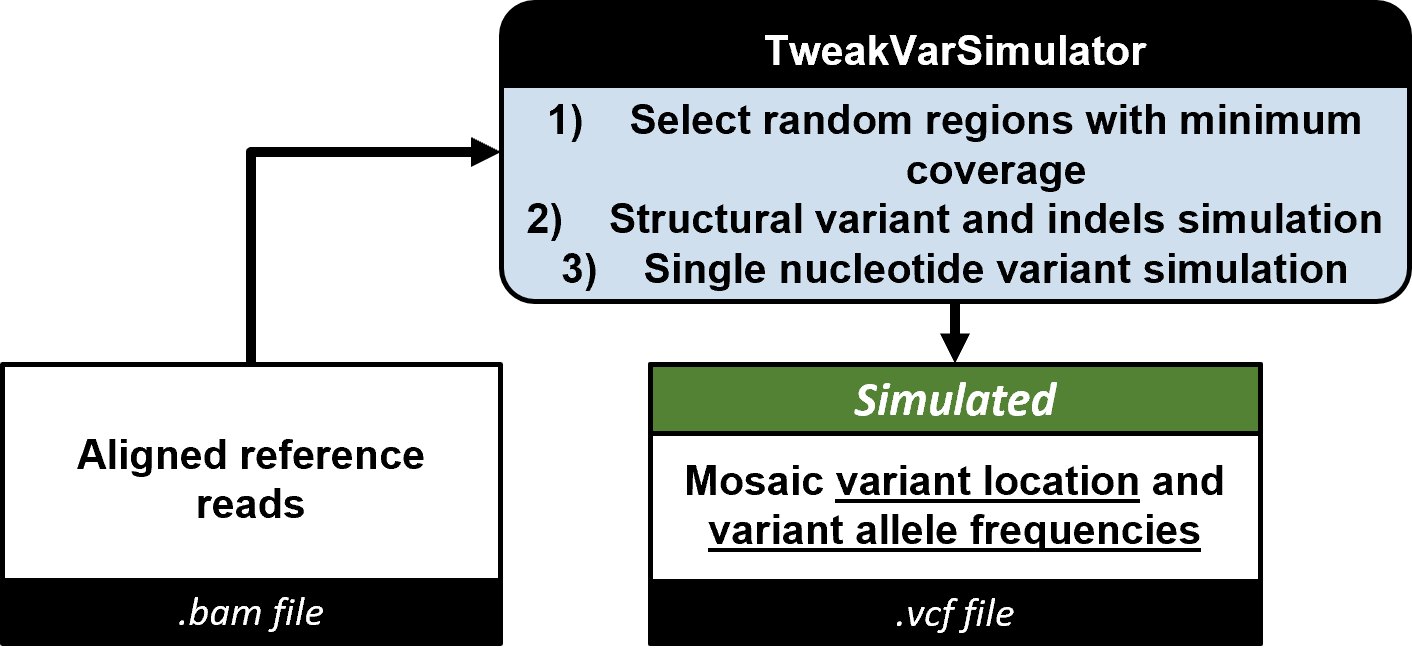

### Xinchang Zheng.jpg

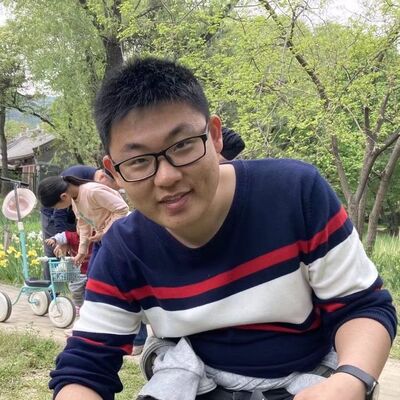
